# Elevated genetic risk for multiple sclerosis originated in Steppe Pastoralist populations

**DOI:** 10.1101/2022.09.23.509097

**Authors:** William Barrie, Yaoling Yang, Evan K. Irving-Pease, Kathrine E. Attfield, Gabriele Scorrano, Lise Torp Jensen, Angelos P. Armen, Evangelos Antonios Dimopoulos, Aaron Stern, Alba Refoyo-Martinez, Abigail Ramsøe, Charleen Gaunitz, Fabrice Demeter, Marie Louise S. Jørkov, Stig Bermann Møller, Bente Springborg, Lutz Klassen, Inger Marie Hyldgård, Niels Wickmann, Lasse Vinner, Thorfinn Sand Korneliussen, Morten E. Allentoft, Martin Sikora, Kristian Kristiansen, Santiago Rodriguez, Rasmus Nielsen, Astrid K. N. Iversen, Daniel J. Lawson, Lars Fugger, Eske Willerslev

**Author notes:** Joint first authors (William Barrie, Yaoling Yang, Evan K. Irving-Pease, Kathrine E. Attfield, Gabriele Scorrano, Lise Torp Jensen). Joint last authors (Astrid K. N. Iversen, Daniel J. Lawson, Lars Fugger, Eske Willerslev).

## Abstract

Multiple sclerosis (MS) is a modern neuro-inflammatory and -degenerative disease, which is most prevalent in Northern Europe. Whilst it is known that inherited risk to MS is located within or within close proximity to immune genes, it is unknown when, where and how this genetic risk originated ^1^. By using the largest ancient genome dataset from the Stone Age ^2^, along with new Medieval and post-Medieval genomes, we show that many of the genetic risk variants for MS rose to higher frequency among pastoralists located on the Pontic Steppe, and were brought into Europe by the Yamnaya-related migration approximately 5,000 years ago. We further show that these MS-associated immunogenetic variants underwent positive selection both within the Steppe population, and later in Europe, likely driven by pathogenic challenges coinciding with dietary, lifestyle, and population density changes. This study highlights the critical importance of this period as a determinant of modern immune responses and its subsequent impact on the risk of developing MS in a changing environment.

## INTRODUCTION

Multiple sclerosis (MS) is an autoimmune disease of the brain and spinal cord that currently affects more than 2.5 million people worldwide ^1^. The prevalence varies markedly with ethnicity and geographical location, with the highest prevalence observed in Europe (142.81 per 100,000 people), and Northern Europeans being particularly susceptible to developing the disease ^3^. The origins and reasons for the geographical variation are poorly understood, yet such biases may hold important clues as to why the prevalence of autoimmune diseases, including MS, has continued to rise during the last 50 years.

While still elusive, MS etiology is thought to involve gene-gene and gene-environmental interactions. Accumulating evidence suggests that exogenous triggers initiate a cascade of events involving a multitude of cells and immune pathways in genetically vulnerable individuals, which may ultimately lead to MS neuropathology ^1^.

Genome-wide association studies have identified 233 commonly occurring genetic variants that are associated with MS; 32 variants are located in the HLA region and 201 outside the HLA region ^4^. The strongest MS associations are found in the HLA region with the most prominent of these,

HLA-DRB1*15:01, conferring an approximately three-fold increase in the risk of MS in individuals carrying at least one copy of the allele. Collectively, genetic factors are estimated to explain approximately 30% of the overall disease risk, while environmental and lifestyle factors are considered the major contributors to MS. Such determinants may include geographically varying exposures like infections and low sun exposure/vitamin D deficiency. For instance, while infection with Epstein-Barr virus (EBV) frequently occurs in childhood and usually is symptomless, delayed infection into early adulthood, as typically observed in countries with high standards of hygiene, is associated with a 32-fold increased risk of MS ^5,6^. Lifestyle factors associated with increased MS risk such as smoking, obesity during adolescence, and nutrition/gut health also vary geographically ^7^.

Dysregulations including autoimmunity in modern immune systems could also result from an altered pressure of pathogens, creating a shift in the delicate balance of pro- and anti-inflammatory pathways^8^.

European genetic ancestry (henceforth “ancestry”) has been postulated to explain part of the global difference in MS prevalence in admixed populations ^9^. Specifically, MS cases in African Americans exhibit increased European ancestry in the HLA region compared to controls, with European haplotypes conferring more MS risk for most HLA alleles, including HLA-DRB1*15:01. Conversely, Asian American cases have decreased European ancestry in the HLA region compared to controls. Although Ancient European ancestry and MS risk in Europe are known to be geographically structured (Figure 1a-b), the effect of ancestry variation within Europe on MS prevalence is unknown.

**Figure 1:**
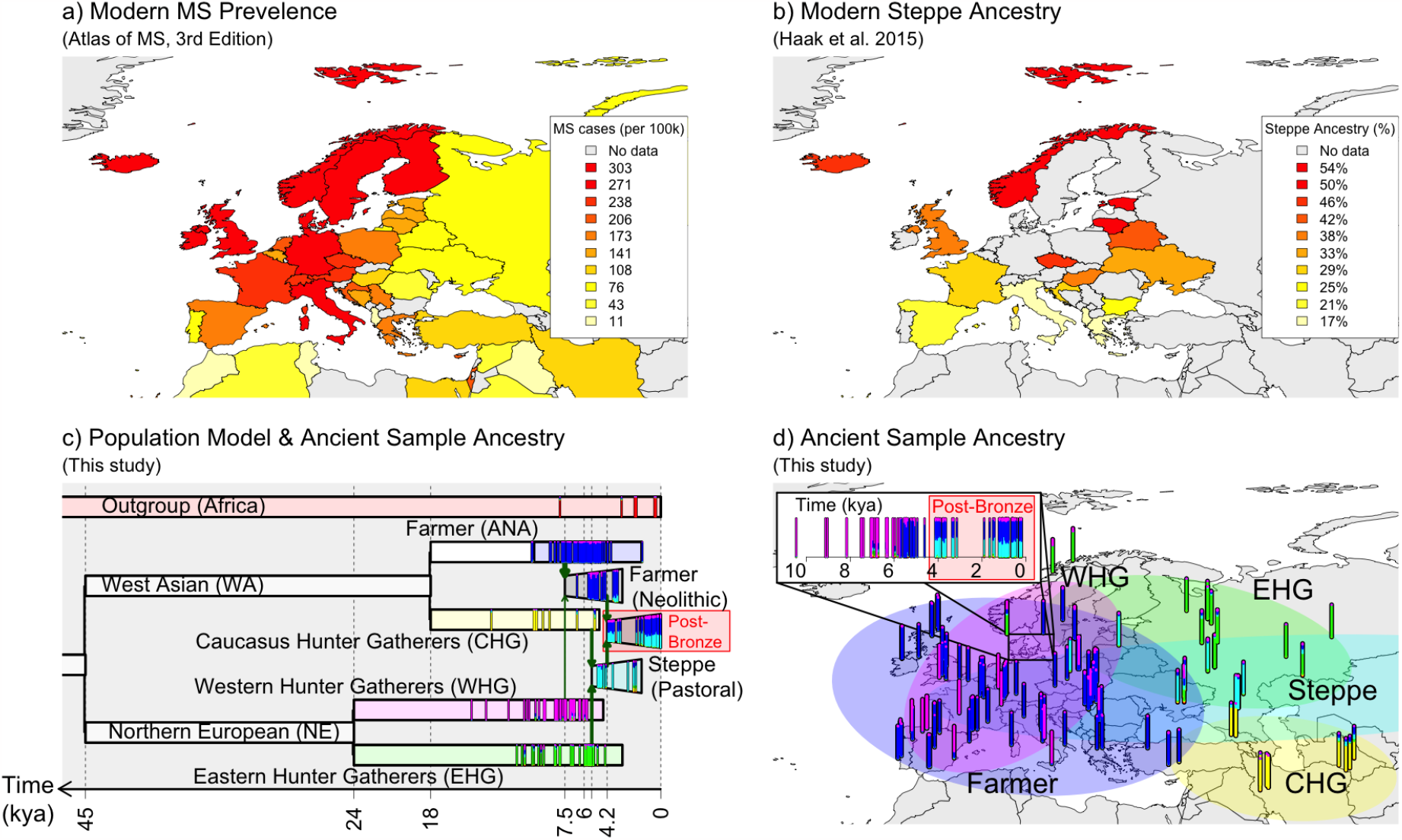
Population history of Europe is associated with modern-day distribution of MS. a) Modern-day geographical distribution of MS in Europe. Prevalence data for MS (cases per 100,000) was obtained from the Atlas of MS - 3rd edition^3^. b) Steppe ancestry in modern samples as estimated by ^10^. c-d) A model of European prehistory ^11^ onto which our reference samples have been projected using NNLS of the Population Painting (see Methods), and the same data represented spatially (kya = thousand years before present). Samples are vertical bars representing their “admixture estimate” estimated by NNLS (Methods) from six ancestries: Eastern Hunter-Gatherers (EHG; green), Western Hunter-Gatherers (WHG; pink), Caucasus Hunter-Gatherers (CHG; yellow), Farmer (ANA+Neolithic; blue), Steppe (cyan) or an Outgroup (represented by ancient Africans, red). Important population expansions are shown as growing bars and “recent” (post-Bronze age) non-reference admixed populations are shown for the Denmark time-transect (see Extended Data Figure 2 for details). Chronologically, WHG and EHG were largely replaced by Farmers amid demographic changes during the “Neolithic transition” around 9,000 years ago. Later migrations during the Bronze Age about 5,000 years ago brought a roughly equal Steppe ancestry component from the Pontic-Caspian Steppe to Europe, an ancestry descended from the EHG from the Middle Don River region and CHG ^2^. Steppe ancestry has been associated with the Yamnaya culture and then with the expansion westwards of the Corded Ware Complex and Bell Beaker culture, and the eastwards expansion in the form of the Afanasievo culture ^10,12^.

Present-day ancestral variation can be modelled as a mixture of genetic ancestries derived from ancient populations, who can be distinguished by their subsistence lifestyle: Western Hunter-Gatherers (WHG), Eastern Hunter-Gatherers (EHG), Caucasus Hunter-Gatherers (CHG), Farmers (ANA + Neolithic), and Steppe Pastoralists (Figure 1c-d). Using the largest ancient genome dataset from the Stone Age, presented in the accompanying study “Population Genomics of Stone Age Eurasia” ^2^, coupled with new Medieval and post-Medieval genomes, we quantified present-day European genetic ancestry with respect to these ancestral populations to identify signals of lifestyle-specific evolution. Then we determined whether the variants associated with an increased risk for MS have undergone positive selection. We asked when selection occurred and whether the targets of selection were specific to lifestyle. Finally, we examined the environmental conditions that may have caused selection for risk variants, including human subsistence practice and exposure to pathogens. An overview of the evidence provided by all methods used can be found in Extended Data Figure 1.

## RESULTS

To examine the ancestry patterns within modern genomes, we estimated ancestry at specific loci (“local ancestry” labels) for ∼410,000 self-identified “white British” individuals in the UK Biobank ^13^, using a reference panel of 318 ancient DNA (aDNA) samples (Figure 1; Extended Data Figure 2; ^14^) from the Mesolithic and Neolithic, including Steppe pastoralists (Methods). Comparing the ancestry at each labelled single nucleotide polymorphism (SNP, n = 549,323) to genome-wide ancestry in the UK Biobank provided an “anomaly score”. Two regions stood out as having the most extreme ancestry compositions (Figure 2a): the LCT/MCM6 region on chromosome 2, well-established as regulating lactase persistence ^14,15^, and the HLA region on chromosome 6.

**Figure 2.**
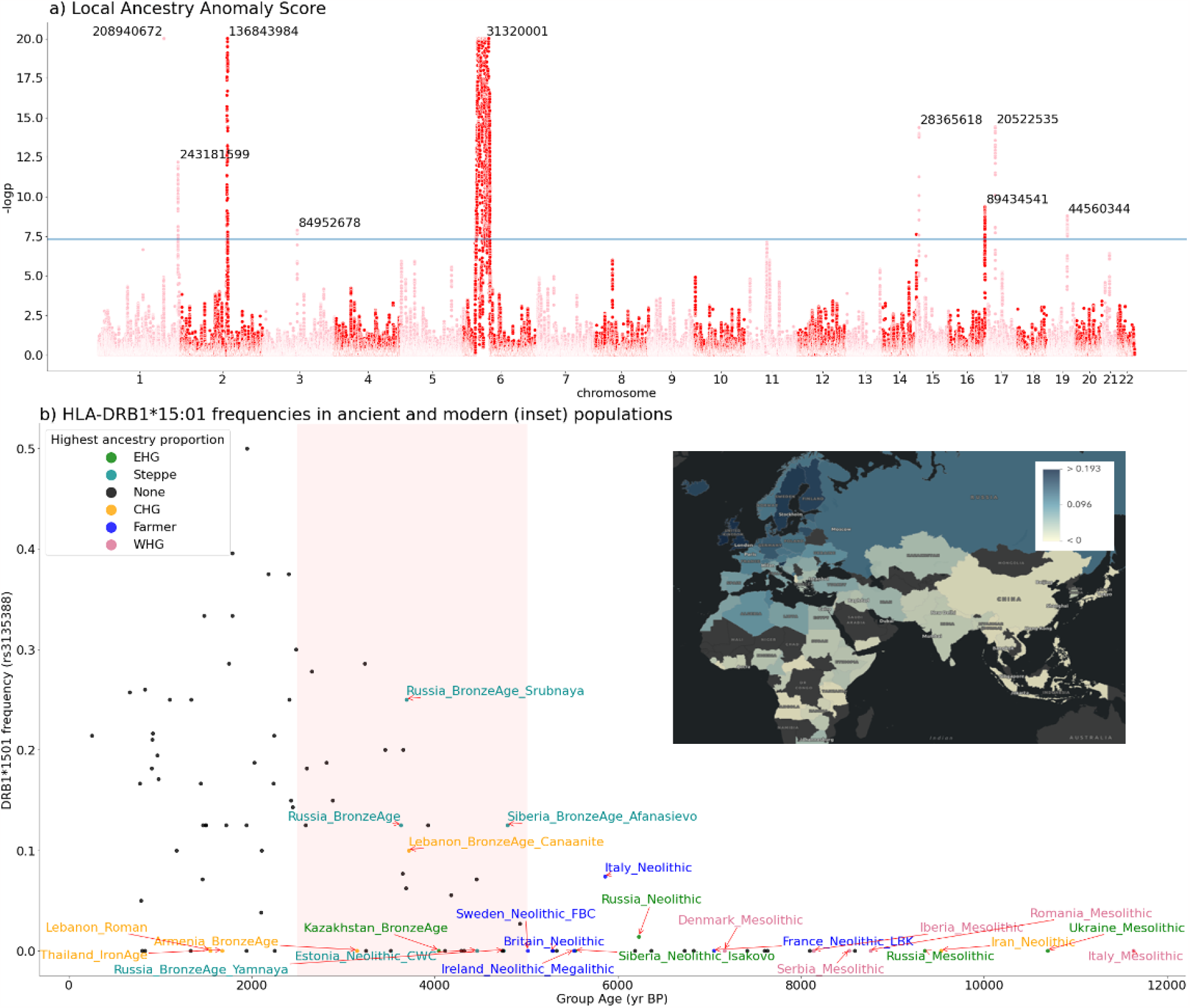
Areas of unusual local ancestry in the genome, and ancient and modern frequencies of DRB1*15:01. a): Local Ancestry Anomaly Score measuring the difference between the local ancestry and the genome-wide average (capped at -log10(p) = 20; see Methods). Peaks are labelled with chromosome position (build GRCh37/hg19). b) HLA-DRB1*15:01 frequencies (Y-axis) in ancient populations over time (X-axis, yr BP = years before present); this is the highest effect variant for MS risk (calculated using rs3135388 tag SNP). For each ancestry (CHG, EHG, WHG, ANA, Steppe), the five populations with the highest amount of that ancestry are coloured and labelled; other populations are shown as black points. DRB1*15:01 was present in one sample before the Steppe expansion, but rose to high frequency during the Yamnaya formation (approximate time period shaded red). The geographical distribution of DRB1*15:01 frequency in modern populations is also shown (inset).

The HLA region is strongly associated with autoimmune diseases ^16^, of which we examined multiple sclerosis (MS) and rheumatoid arthritis (RA), a common systemic inflammatory disease that characteristically affects the joints, in detail, using the largest ancient genome dataset from the Stone Age (full description in ^2^) coupled with 86 new Medieval and post-Medieval genomes from Denmark (Extended Data Figure 2, Supplementary Note 1, ST1). This dataset totals 1,750 imputed diploid shotgun-sequenced ancient genomes (ST13), of which 1,509 are from Eurasia; alongside modern data, with our newly published genomes we have an almost complete transect from approximately 10,000 years ago to the present.

The frequencies of alleles conferring the highest risk for MS in our ancient groups, all of which are within the HLA class II region, show striking patterns. In particular, the tag SNP (rs3135388-T) for HLA-DRB1*15:01, the largest genetic risk factor for MS ^1^, is first observed in an Italian Neolithic individual (sampleId R3 from Grotta Continenza, C14 dated to between 5,836-5,723 BCE, coverage 4.05 X) and rapidly increased in frequency around the time of the emergence of the Yamnaya culture around 5,300 years ago in Steppe and Steppe-derived populations (Figure 2). From risk allele frequencies of individuals in the UK Biobank born in, and of a “typical ancestral background” for, each country ^14^, we found HLA-DRB1*15:01 frequency peaks in modern populations of Finland, Sweden and Iceland, and in ancient populations with high Steppe ancestry (Figure 2b, inset).

To investigate the risk of a particular genetic ancestry at all MS-associated fine-mapped loci present in the UK Biobank imputed dataset (n = 205/233, ^4^, see Methods), we used the local ancestry dataset to calculate a risk ratio (see Methods: Weighted Average Prevalence) for each ancestry. For MS, Steppe ancestry has the highest risk ratio in nearly all HLA SNPs, while Farmer and “Outgroup” ancestry (represented by ancient Africans) are often the most protective (Figure 3a), meaning a Steppe-derived haplotype at these positions confers MS risk.

**Figure 3:**
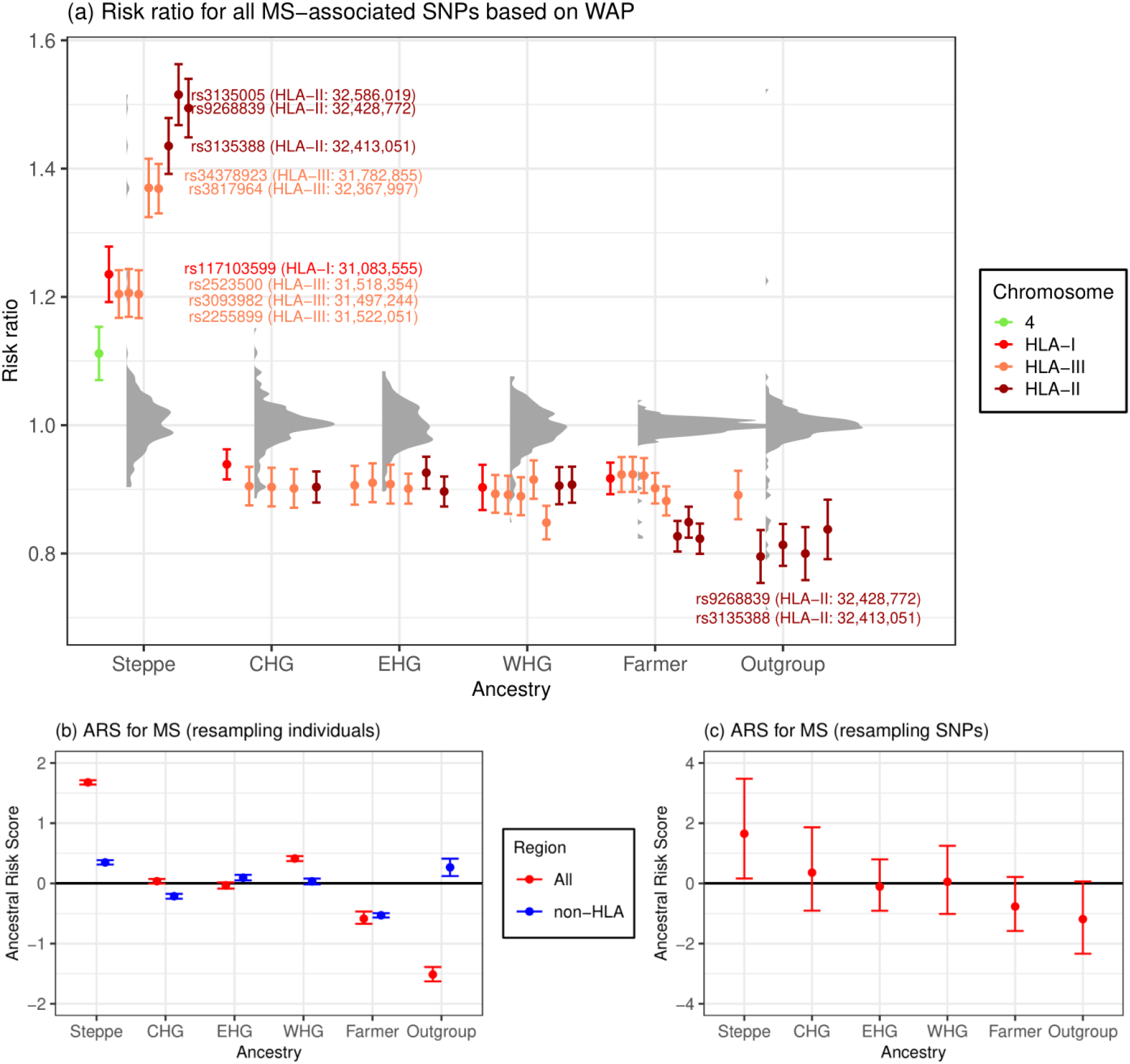
Associations between local ancestry at fine-mapped MS-associated SNPs and MS in a modern population. a) Risk ratio of SNPs for MS based on weighted average prevalence (WAP; see Methods), when decomposed by inferred ancestry. A mean and standard deviation were calculated for each ancestry based on bootstrap resampling, for each chromosome. The distribution of risk ratios at each ancestry is shown as a raincloud plot. SNPs significant at the 1% level are shown individually, coloured by chromosome or HLA region, and those with risk ratio >1.2 or <0.8 are annotated with rsID, HLA region and position (build GRCh37/hg19). b-c) Ancestral Risk Scores (ARS, see Methods) for MS. Confidence intervals are estimated by either bootstrapping over individuals (b, which can be interpreted as testing power to reject a null hypothesis of no association between MS and ancestry) and bootstrapping over SNPs (c, which can be interpreted as testing whether ancestry is associated with MS genome-wide). We show results for all associated SNPs (red) and non-HLA SNPs only (blue) when bootstrapping over individuals.

Having shown that some ancestries carry higher risk at particular SNPs, we wanted to calculate an aggregate risk score for each ancestry. We used a statistic, the Ancestral Risk Score (ARS, introduced in ^14^, which is equivalent to a polygenic risk score (PRS) for a modern individual consisting of entirely one ancestry. ARS offers an improvement on calculating PRS using ancient genotype calls directly, as it mitigates the effects of low aDNA sample numbers and bias ^17^, while being robust to intervening drift and selection. We used effect size estimates from previous association studies, under an additive model, with confidence intervals obtained via an accelerated bootstrap ^18^ (Supplementary Note 4). In the ARS for MS (Figure 3b), Steppe ancestry had the largest risk, followed by WHG, CHG and EHG; Farmer and Outgroup ancestry had the lowest ARS. Therefore, Steppe ancestry is contributing the most risk for MS across all associated SNPs. We tested for a genome-wide association by resampling loci, and found that Steppe risk is much reduced but still clearly exceeds Farmer (Figure 3c). Although most of the signal is driven by SNPs in the HLA region, this pattern persists even when excluding these SNPs (Figure 3b).

The fact that Steppe ancestry confers risk at all but two MS-associated HLA SNPs (Supplementary Information Figure S3.4b) implies that these alleles have a common evolutionary history. We therefore investigated whether ancestry could be used for phenotype prediction. We conducted three types of association analysis in the UK Biobank for disease-associated SNPs, controlling for age, sex and the first 18 PCs. The first was a regular SNP-based association analysis, as in a GWAS. The second tested for association with local ancestry probabilities instead of genotype values (Supplementary Note 3). The third was based on Haplotype Trend Regression (HTR), which is used to detect interactions between SNPs ^19^ by treating haplotypes as a set of features from which to predict a trait, instead of using SNPs as in a regular GWAS. We developed a new method called Haplotype Trend Regression with eXtra flexibility (HTRX, Supplementary Note 5, more details in ^20^ that searches for haplotype patterns that include single SNPs and non-contiguous haplotypes. To evaluate the performance of our models and prevent overfitting, we assessed its ability to predict out-of-sample data, which measures how well the model can generalise to new data. We showed by simulation (Supplementary Information Figure S5.1) that HTRX explains the same amount of variance as a regular GWAS when interactions are absent and more variance as interaction strength increases.

Although our cohort of self-identified “white British” individuals is relatively underpowered with respect to MS (cases = 1,949; controls = 398,049; prevalence = 0.487%), MS was associated with Steppe and Farmer ancestry (p < 1e-10) in the HLA region (Supplementary Information Figure S3.2). In 3 out of 4 main LD blocks within the HLA region (class I, two subregions of class II determined by LD blocks at 32.41-32.68Mb and 33.04-33.08Mb, and class III), local ancestry explains significantly more variation than genotypes (Figure 4; measured by average out-of-sample McFadden’s *R*^2^ for logistic regression, see Methods). While the increased performance of local ancestry in some regions over regular GWAS can be explained by tagging of SNPs outside the region, the increased HTRX performance over GWAS quantifies the total effect of a haplotype, including rare SNPs and epistasis. Across the entire HLA region, haplotypes explain more out-of-sample variation than GWAS (at least 2.90%, compared to 2.48%). Interaction signals are also observed within the HLA class I, within class II, and between class I and III regions.

**Figure 4:**
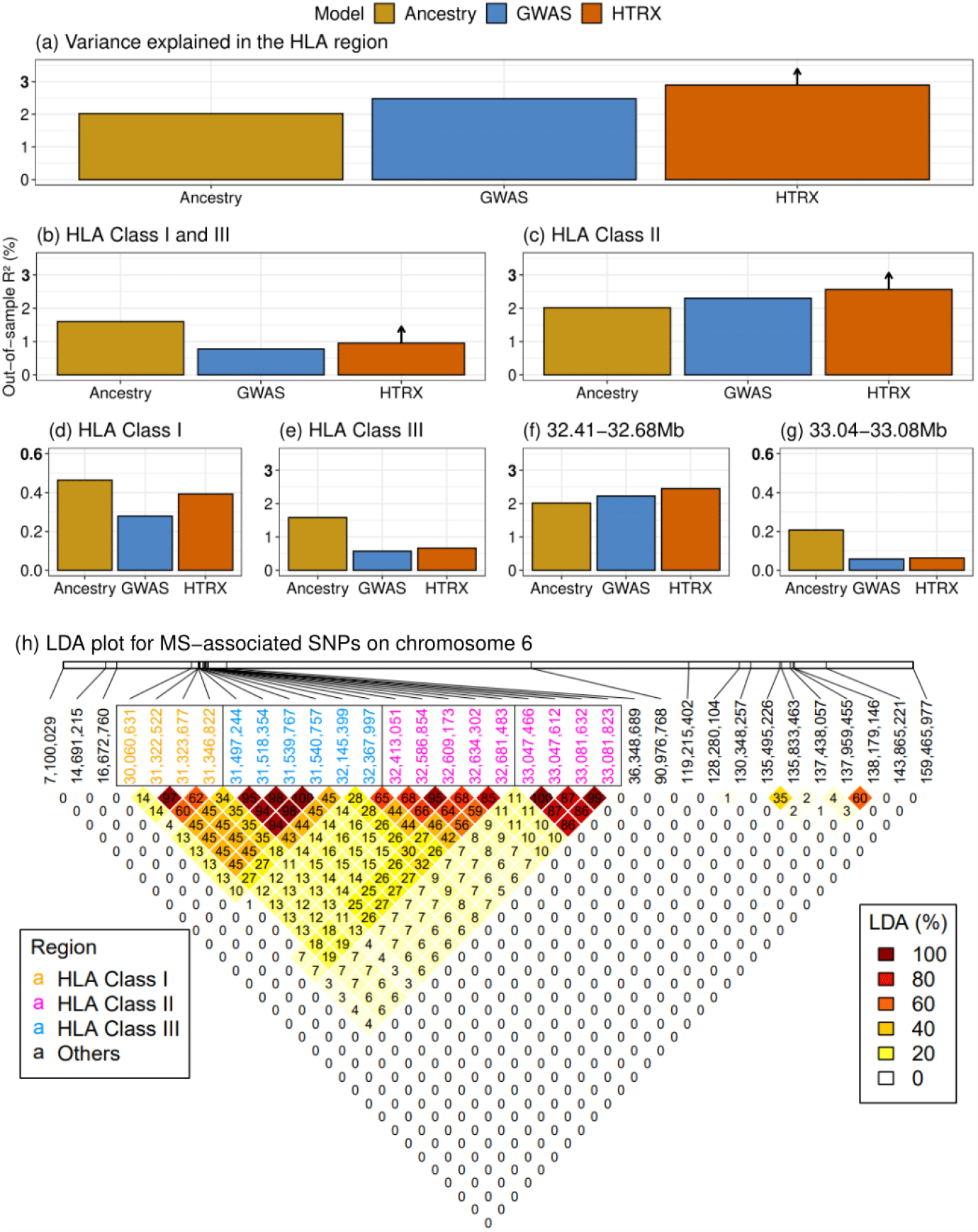
MS association in the HLA. Comparison of variance explained in MS within the UK Biobank, for all fine-mapped HLA SNPs with an independent contribution ^4^. The plots compare GWAS (treating SNPs as having independent effects), local ancestry at those SNPs, and HTRX (haplotypes), after accounting for covariates (Methods). a) is for fine-mapped MS-associated SNPs in the HLA. b) is HLA class I and -III, c) is HLA class II, d) is HLA class I, e) is HLA class III, f) and g) are subregions of HLA class II chosen from LD. HTRX has small “up-arrows” where these are lower bounds (Methods). h) Genetic correlations in the HLA region at our time-depth from Ancestry-based LD (LDA, see Methods) and Supplementary Information Figure S7.5 for LD.

Multiple SNPs at the 32.41-32.68Mb region are Steppe-associated, have high MS odds ratios, and are in LDA (Figure 4), which may explain the increased HTRX predictive performance. We further tested whether co-occurring ancestries at each loci were associated with MS (Methods; Supplementary Information Figure S3.3), but found no evidence that risk was associated with any ancestry other than Steppe.

Having established that Steppe ancestry contributes most of the HLA-associated risk for MS, we investigated whether MS risk evolved under selection. We tested for evidence of directional selection across all associated SNPs, decomposed by ancestry over time. This test uses a novel “pathway-based chromosome painting” technique (Methods: Pathway painting) based on inference of a sample’s nearest neighbours in the marginal trees of an ancestral recombination graph (ARG) that contains labelled individuals ^14,21^. The resulting ancestral path labels, for haplotypes in both ancient and modern individuals, allowed us to infer allele frequency trajectories for risk associated variants, while controlling for changes in admixture proportions through time. These paths extend backwards from the present day to approximately 15,000 years ago, and are labelled with the unique population that a path travels through (ANA: Anatolian Farmers; CHG: Caucasus Hunter-Gatherers, EHG: Eastern Hunter-Gatherers, WHG: Western Hunter-Gatherers). Because it uses distinct pathways, this approach does not use the labels of the relatively recent Steppe admixture or outgroup populations, and the path labels are not representative of a continuous population, but represent a path backwards in time that encompasses that population. For example, the CHG path originates in Caucasus Hunter-Gatherers, before merging with EHG to form the Steppe population, and then merges with other ancestries in later European populations (Figure 1).

For our ancestry path analysis, a substantial fraction of the fine-mapped MS variants were not imputed in our ancient dataset, due to quality control filtering and the difficulty of accurately inferring HLA alleles in ancient samples ^22^. To address this, we LD-pruned genome-wide significant summary statistics from the same study ^4^, for which we could reliably assign ancestry path labels (n = 62, see Methods). This allowed us to test for polygenic selection across disease-associated variants using CLUES ^23^ and PALM ^24^.

For MS, we found evidence that disease risk was selectively increased when considering all ancestries collectively (p = 1.02e-5; ω = 0.017), between 5,000-2,000 years ago (Figure 5). Conditioning on each of the four long-term ancestral paths (CHG, EHG, WHG, and ANA), we found a statistically significant signal of selection in the WHG (p = 7.22e-5; ω = 0.021), EHG (p = 2.60e-3; ω = 0.016) and CHG paths (p = 3.06e-2; ω = 0.009), but not in the ANA path (p = 0.64; ω = 0.004). Again, it is likely that selection occurred in the pastoralist population of the Steppe, as that population consists of approximately equal proportions of EHG and CHG ancestry ^11^ (Figure 1). The SNP driving the largest change in genetic risk over time in the pan-ancestry analysis was was rs3129934 (p = 1.31e-11; s = 0.018), which tags the HLA-DRB1*15:01 haplotype ^25^. We also tested three other SNPs that tag the HLA-DRB1*15:01 haplotype (rs3129889, rs3135388 and rs3135391) for evidence of selection, and found that the ancestry stratified signal was consistently strongest in CHG (Figure 5b).

**Figure 5:**
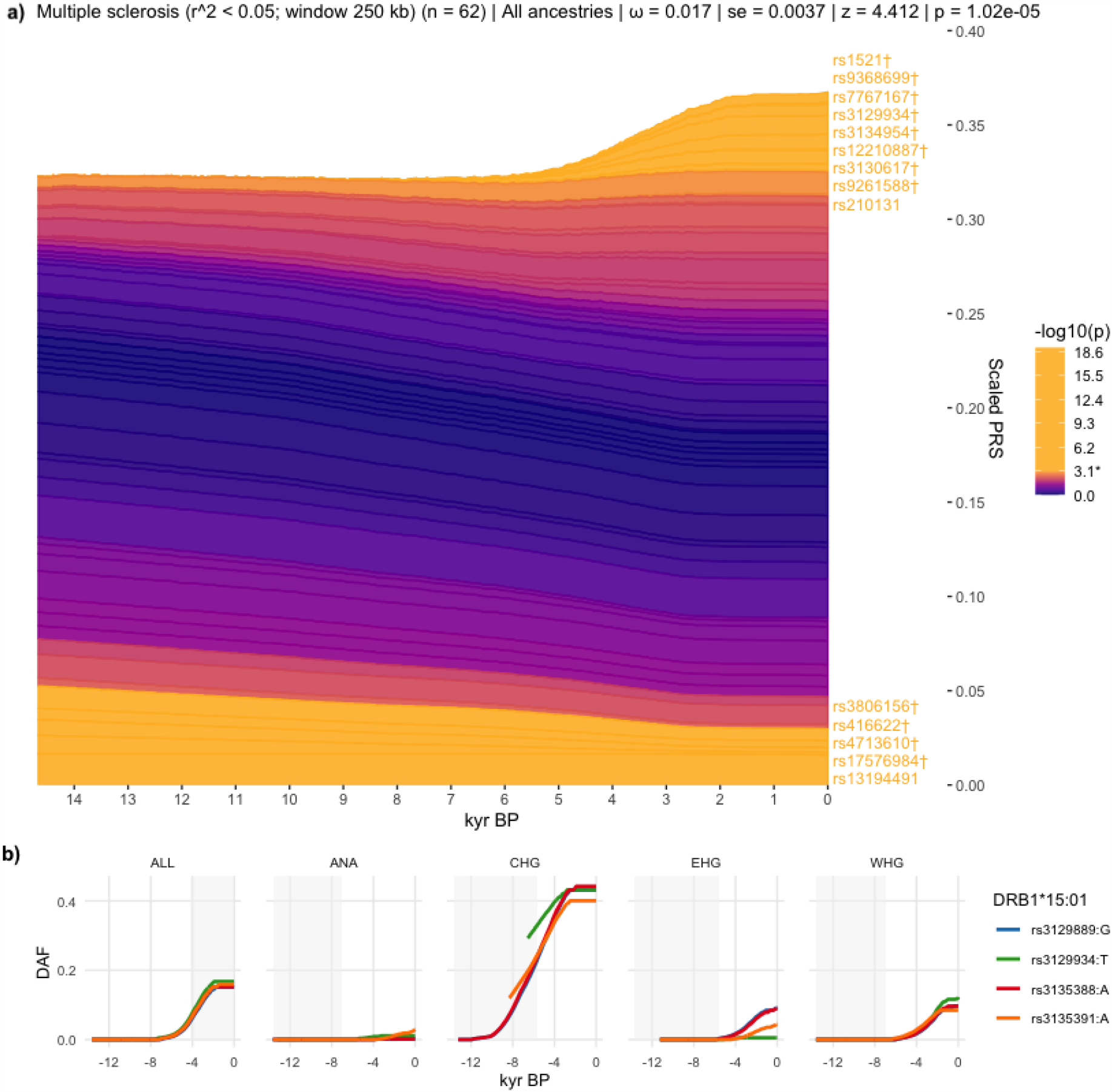
Evidence for selection on MS-associated SNPs. a) Stacked line plot of the pan-ancestry PALM analysis for MS, showing the contribution of SNPs to disease risk over time. SNPs are shown as stacked lines, the width of each line being proportional to the population frequency of the positive risk allele, weighted by its effect size. When a line widens over time the positive risk allele has increased in frequency, and vice versa. SNPs are sorted by the magnitude and direction of selection, with positively selected SNPs at the top, negatively selected SNPs at the bottom, and neutral SNPs in the middle. SNPs are coloured by their corresponding p-value in a single locus selection test. The asterisk marks the Bonferroni corrected significance threshold, and nominally significant SNPs are shown in yellow and labelled by their rsIDs. SNPs marked with the dagger symbol are located in the HLA locus. The Y-axis shows the scaled average polygenic risk score (PRS) in the population, ranging from 0 to 1, with 1 corresponding to the maximum possible average PRS (i.e. when all individuals in the population are homozygous for all positive risk alleles) and the X-axis shows time in units of thousands of years before present (kyr BP). b) Maximum likelihood trajectories for four SNPs tagging DRB1*15:01, for all ancestry paths combined (ALL) and for each path separately (see Extended Data Figure 1, and Methods: Pathway painting). Portions of the trajectories with high uncertainty (i.e., posterior density <0.08) have been masked. The background is shaded for the approximate time period in which the ancestry existed as an actual population. The Y-axis shows the derived allele frequency (DAF), and the X-axis shows time in units of thousands of years before present (kyr BP).

To further examine the nature of selection, we developed a new summary statistic, Linkage Disequilibrium of Ancestry (LDA). LDA is the correlation between local ancestries at two SNPs, measuring whether recombination events between ancestries are high compared to recombination events within ancestries. We subsequently defined the “LDA score” of a SNP as the total LDA of the SNP with the rest of the genome. A high LDA score indicates that the haplotype inherited from the reference population is longer than expected, while a low score indicates that the haplotype is shorter than expected (i.e. underwent more recombination). For example, the LCT/MCM6 region exhibits a high LDA score (Extended Data Figure 5), as expected from a relatively recent selective sweep ^26^.

The HLA has significantly *lower* LDA scores than the rest of chromosome 6 (Extended Data Figure 5). We simulated the LDA score under selection (Supplementary Information Figure S7.1; Methods), and found that single locus selection cannot explain this signal (Supplementary Information Figure S7.2-3). Instead, different loci in LD must have independently reached high frequency in different ancestral populations that admixed, with selection favouring haplotypes of mixed ancestry over single-ancestry haplotypes. Extending multi-SNP selection models ^27^, our explanation (Supplementary Information Figure S7.1) is that at least two separate loci rose selectively in separate populations that later admixed and remained selected in the HLA, justifying a new term, “recombinant favouring selection”. This means that there was selection for diverse ancestry in the HLA region, driven by recombination.

The HLA region contains the highest “Outgroup” ancestry anywhere on the genome (Figure 6), reflecting high nucleotide diversity. Unlike other measures of balancing selection such as Fst (Figure 6), LDA describes excess ancestry LD from specific, dated populations and therefore is an independent signal. For the HLA class II region, the selection measures all line up (LDA score, Fst, pi), but for class I, the LDA score has an additional non-diverse minimum at 30.8Mb, implying that here the genome is ancestrally diverse but genetically strongly constrained. The LDA score is thus informative about the type of selection being detected, and whether it has been subject to change.

**Figure 6:**
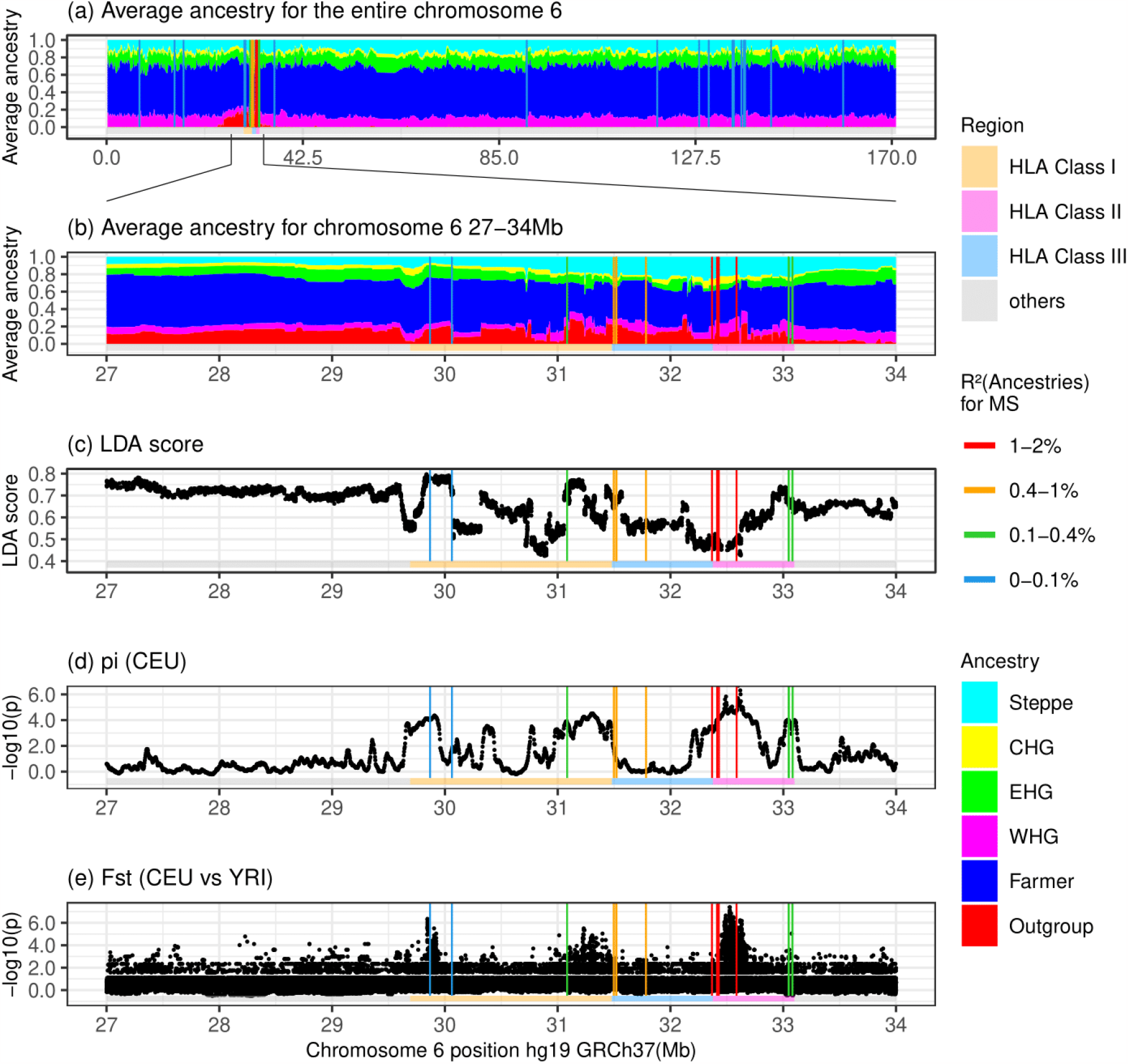
Signatures of selection at the HLA locus showing different regions of the HLA (horizontal coloured bar) and locations of MS-associated SNPs (vertical lines, coloured by the variance explained by 6 ancestries). a): Whole Chromosome 6 “local ancestry” decomposition by genetic position. b). HLA “local ancestry” decomposition by genetic position. c): LDA score; low values are indicative of selection for multiple linked loci, while high values indicate positive selection. d): pi scores (nucleotide diversity) for CEU (Northern and Western European ancestry). MS-associated SNPs fall in highly diverse regions of the HLA. e): Fst scores (divergence between two populations) for CEU vs YRI(Yoruba); locally higher scores indicate regions that have undergone differential selection between the two populations.

Because MS would not have conferred a fitness advantage on ancient individuals, it is likely that this selection was driven by traits with shared genetic architecture, of which increased risk for MS in the present is a pleiotropic by-product. We therefore looked at LD-pruned MS-associated SNPs that showed statistically significant evidence for selection using CLUES (n = 32), in one or more ancestries, and which also had a genome-wide significant trait association (p < 5e-8) in any of the 4,359 traits from the UK Biobank (^13^; UK Biobank Neale Lab, Round 2: http://www.nealelab.is/uk-biobank/) and any of the 2,202 traits in the FinnGen study ^28^. We observed that all selected SNPs were also associated with multiple other traits (Supplementary Information Figures S6.8 - S6.16). To determine if the observed signal of polygenic selection favouring MS risk could be better explained by selection acting on a genetically correlated trait, we performed a systematic analysis of traits in UK Biobank and FinnGen with at least 20% overlap among the MS-associated selected SNPs (n = 115 traits). Using a joint test in PALM, specifically designed for disentangling polygenic selection on correlated traits, we found no UK Biobank or FinnGen traits where the selection signal favouring MS risk was significantly attenuated by selection acting on a genetically correlated trait, when accounting for the number of tests (Supplementary Note 6). This demonstrates that the selection signal for MS could not be explained by selection acting on any genetically correlated trait that we tested.

Because both the UK Biobank and FinnGen are underpowered with respect to many traits and diseases, we also undertook a manual literature search (see Methods) for all LD-pruned MS-associated SNPs which showed statistically significant evidence for selection using CLUES (n = 32, of which 25 (78%) are in the HLA region). We found that most of the alleles under positive selection are associated with protective effects against specific pathogens (virus, bacteria, fungi and parasites) and/or infectious diseases within one or several ancestral paths (disease or pathogen associated/total selected in ancestry path: pan-ancestry 11/14; ANA 8/9; CHG 6/9; EHG 6/7; WHG 17/18, Supplementary Note 8, ST11, Extended Data Figure 6), although we note that GWAS data for many infectious diseases are not available. We observed that the selected alleles had protective associations with several chronic viruses (EBV, varicella-zoster virus (VZV), herpes simplex virus (HSV), and cytomegalovirus (CMV)) and to viruses or diseases not associated with transmission in small hunter-gatherer groups (e.g., mumps and influenza). Moreover, many selected alleles conferred a reduction of risk of parasites, of skin and subcutaneous tissue, gastrointestinal, respiratory, urinary tract, and sexually transmitted infections, or of pathogens associated with these or other infections (e.g. *Clostridium difficile, Streptococcus pyogene*s, *Mycobacterium tuberculosis*, and coronavirus) (Supplementary Note 8, ST11, Extended Data Figure 6). We emphasise that although this evidence is strongly suggestive, many of these putative associations may not be statistically robust due to underpowered GWASs and the bias in candidate gene studies.

We compared these findings for MS with results for RA, which in contrast to MS is a *systemic* inflammatory disease, though it is mostly known for its characteristic joint lesions ^16^. Our findings show a strikingly different ancestry risk profile. HLA-DRB1*04:01 is the largest genetic risk factor for RA; in the CLUES analysis, the tag SNP for this allele (rs660895) displayed evidence of continuous negative selection until approximately 3,000 years ago (p = 7.95e-7, Extended Data Figure 4). We found that WHG and EHG ancestries often confer the most risk at SNPs associated with RA (Relative Risk ratio of RA-associated SNPs based on WAP, see Methods); and these ancestries have contributed the greatest risk for RA on aggregate, reflected in a higher ARS for these ancestries (Supplementary Note 4), while Steppe and Outgroup ancestry have the lowest scores (Extended Data Figure 3). These results were recapitulated in the local ancestry GWAS (Supplementary Note 3).

We found that RA-associated SNPs have undergone negative polygenic selection (p = 3.26e-3, Extended Data Figure 4) over the last approximately 15,000 years. When decomposed by ancestry path, we found that all paths exhibited a negative selection gradient, but none achieved nominal significance; although the CHG (p = 6.33e-2; ω = -0.014) path came close.

These results demonstrate that genetic risk for RA was higher in the distant past, in contrast to MS, with RA-associated risk variants present at higher frequencies in European hunter-gatherer populations before the arrival of agriculture. In order to understand what caused the high risk in hunter-gatherer populations and subsequent negative selection, we again undertook a manual literature search for pleiotropic effects of LD-pruned SNPs which showed statistically significant evidence for selection (n = 55 of which 36 (65%) are in the HLA region). We found that the majority of selected SNPs were associated with protection against distinct pathogens and/or infectious diseases across all paths (disease or pathogen associated/total selected in ancestry path: pan-ancestry 16/20; ANA 12/16; CHG 8/13; EHG 14/20; WHG 16/21). We found that selected RA-risk alleles were typically linked to the same pathogens or diseases as in the MS analysis, although some SNPs were protective against pathogens or diseases not observed in the MS-risk analysis (e.g., *Entamoeba histolytica, measles, viral hepatitis, arthropod-borne viral fevers and viral haemorrhagic fevers*, and pneumococcal pneumonia) (Supplementary Note 8, ST12, Extended Data Figure 6).

## DISCUSSION

The last 10,000 years have seen some of the most extreme global changes in lifestyle, with the emergence of farming in some regions and pastoralism in others. While 5,000 years ago Farmer ancestry predominated across Europe, a relatively diverged genetic ancestry arrived with the Steppe migrations around this time ^10,12^. We have shown that this genetic ancestry contributes the most genetic risk for MS today, and that these variants were the result of positive selection coinciding with the emergence of a pastoralist lifestyle on the Pontic-Caspian Steppe, and continued selection in the subsequent admixed populations in Europe. These results address the long-standing debate around the north-south gradient in MS prevalence in Europe, and suggest that the Steppe ancestry gradient in modern populations - specifically at the HLA region - across the continent causes this phenomenon, in combination with environmental factors. Furthermore, while epistasis between MS-associated variants in the HLA region has been demonstrated before ^29,30,31,32^, we have shown that accounting for this explains more variance than independent SNPs effects alone. Many of the haplotypes carrying these risk alleles have ancestry-specific origins, which could be exploited for individual risk prediction and may offer a pathway from genetic ancestry associations into a mechanistic understanding of MS risk. We have compared these findings with results for RA, another HLA class II associated chronic inflammatory disease, and found that the genetic risk for RA exhibits a contrasting pattern; genetic risk was highest in Stone Age hunter-gatherer ancestry and decreased over time.

Our interpretation of this history is that co-evolution between pathogens and their human hosts has resulted in massive and divergent genetic ancestry-specific selection on immune response genes according to lifestyle and environment, driven by a range of pathogenic drivers, and “recombinant favouring selection” after these populations merged. Similar examples of pathogen-driven evolution have recently been published ^33,34^. The Late Neolithic and Bronze Age was a time of massively increased infectious diseases in human populations, due to increased population density as well as contact with, and consumption of, domesticated animals and their products. The most recent common ancestor of many disease-associated pathogens existed in this period, such as *Mycobacterium tuberculosis* (tuberculosis (TB)) ^35,36^, *Yersinia pestis* (plague) ^37,38,39^, measles-morbillivirus (MeV) (measles) ^40^, HSV (herpes) ^41^, and VZV (chickenpox, zoster) ^42,43^. While these diseases are common today, it is difficult to infer their geographic ranges in the past, which may have been more limited: for example, in the fifth century BC, Hippocrates provided the first description of an outbreak of mumps caused by the mumps virus (MuV) ^44^, and populations were not large enough to sustain continuous measles transmission ^40^, which suggests that these viruses were not necessarily endemic at this time.

We have shown that many of the MS- and RA-associated variants under selection confer some resistance to a range of infectious diseases and pathogens (Supplementary Note 8) (for example, HLA-DRB1*15:01 is associated with protection against TB ^45^ and increased risk for lepromatous leprosy ^46^). We are, however, underpowered to detect specific associations beyond this hypothesis due to poor knowledge of the distribution and diversity of past diseases, poor preservation of endogenous pathogens in the archaeological record, and a lack of well-powered GWASs for many infectious diseases, partly due to widespread vaccination programs. Together, this evidence suggests that population dispersals, changing lifestyles and increased population density resulted in high and sustained transmission of both new and old pathogens, driving selection in immune response genes which are now associated with autoimmune diseases.

A pattern that repeatedly appears is that of lifestyle change driving changes in risk and phenotypic outcomes. We have shown that in the past environmental changes driven by lifestyle innovation inadvertently drove an increase in genetic risk for MS. Today, with increasing prevalence of MS cases observed over the last five decades ^47,48^, we again observe a striking correlation with changes in our environment, including lifestyle choices and improved hygiene, which no longer favours the previous genetic architecture. Instead, the fine balance of genetically-driven cells within the immune system, which are needed to combat a broad repertoire of pathogens and parasites without harming self-tissue, has been met with new challenges, including a potential absence of requirement. For example, while a population of immune cells, CD4+ T helper 1 (Th1), direct strong cellular immune responses against intracellular pathogens, T helper 2 (Th2) cells mediate humoral immune responses against extracellular bacteria and parasites and further have the capacity to guide the restoring of homeostasis, thus preventing damage of the infected tissue via immune-regulatory cytokines. We have shown that the majority of selected MS-associated SNPs are associated with protection against a wide range of infectious challenges, consistent with selection for strong but balanced Th1/Th2 immunity in the Bronze Age, when an increase in exposure to viral and bacterial pathogens and parasites likely took place. Although MS pathogenesis is complex and multicellular in nature, CD4+ Th cells, in particular IFN-ɣ producing Th1 cells and IL-17-producing Th17 cells, play a key role in disease development ^1^. The skewed Th1/Th2 balance observed in MS may partly result from the developed world’s increased sanitation, which has led to a drastically reduced burden of parasites, which the immune system had evolved to efficiently combat ^49^.

Similarly, the new pathogenic challenges associated with agriculture, animal domestication, pastoralism, and higher population densities might have substantially increased the risk of triggering a systemic RA-associated inflammatory state in genetically predisposed individuals. This could lead to an increased risk of a serious outcome following subsequent infections ^50^, years before any potential joint lesions ^51^, resulting in negative selection and thus, might present a parallel between RA-associated inflammation in the Bronze Age and MS today, in which lifestyle changes have exposed previously favourable genetic variants as autoimmune disease risks.

More broadly, it is clear that the late Neolithic and Bronze Age was a critical period in human history during which highly genetically and culturally divergent populations evolved and mixed ^2^. These separate histories dictate the genetic risk and prevalence of several autoimmune diseases today. Surprisingly, the emergence of the pastoralist Steppe lifestyle may have had an impact on immune responses as great as or greater than the emergence of farming during the Neolithic transition, commonly held to be the greatest lifestyle change in human history.

## DATA AVAILABILITY

All collapsed and paired-end sequence data for novel samples sequenced in this study will be made publicly available on the European Nucleotide Archive, together with trimmed sequence alignment map files, aligned using human build GRCh37. Previously published ancient genomic data used in this study are detailed in ST13, and are all already publicly available.

## CODE AVAILABILITY

The modified version of CLUES used in this study is available from https://github.com/standard-aaron/clues. The pipeline and conda environment necessary to replicate the analysis of allele frequency trajectories and polygenic selection in Supplementary Note 6 are available on Github at https://github.com/ekirving/ms_paper. The code to create Ancestry Anomaly scores based on Chromosome painting is on Github at https://github.com/danjlawson/ms_paper. The code to compute LDA and LDA score is available on Github at https://github.com/YaolingYang/LDAandLDAscore. The code for HTRX is on Github at https://github.com/YaolingYang/HTRX. The code for ARS calculation is on Github at https://github.com/will-camb/ms_paper.

## ACKNOWLEDGEMENTS

We extend our thanks to all the former and current staff at the Lundbeck Foundation GeoGenetics Centre and the GeoGenetics Sequencing Core, and to colleagues across the many institutions detailed below. We are particularly grateful to Maria Madrona, Lærke Hansen and Julie Bitz-Thorsen for laboratory assistance; and to Julie Hansen, Sandra Mularczyk, Katja Thorø Michler, Emilie Neerup Nielsen for their help with sampling, and to Line Olsen as project manager for the Lundbeck Foundation GeoGenetics Centre project. We thank UK Biobank Ltd. for access to the UK Biobank genomic resource. We also thank and acknowledge the participants and investigators of the FinnGen study. We are thankful to Illumina Inc. for collaboration. E.W. thanks St. John’s College, Cambridge, for providing a stimulating environment of discussion and learning and the Lundbeck Foundation, the Novo Nordisk Foundation, the Wellcome Trust, the Carlsberg Foundation, and the Danish National Research Foundation for financial support. R.N. acknowledges NIH grant R01GM138634. K.E.A., A.P.A., A.K.N.I., and L.F. thank the OAK Foundation.

## AUTHOR CONTRIBUTIONS

W.B., Y.Y., E.K.I-P., K.E.A., G.S., and L.T.J. contributed equally to this work. A.K.N.I., D.J.L., L.F., and E.W. led the study.

W.B., A.R-M., L.F., R.N., and E.W. conceptualised the study. R.N., K.K., L.F., and E.W. acquired funding for research.

A.R., C.G., F.D., M.L.S.J., S.B.M., B.S., L.K., I.M.H., N.W., L.V., and T.S.K., were involved in sample collection and processing

W.B., Y.Y., E.K.I-P., A.S., S.R., and D.J.L. were involved in developing and applying methodology.

W.B., Y.Y., E.K.I-P., G.S., A.P.A., A.R., E.D., M.S., S.R., A.K.N.I., and D.J.L. undertook formal analyses of data.

W.B., Y.Y., E.K.I-P., K.E.A., and L.T.J., A.K.N.I., L.F., and E.W. drafted the main text (W.B. led this).

W.B., Y.Y., E.K.I-P., G.S., L.T.J., E.D., A.S., F.D., M.L.S.J., S.B.M., B.S., L.K., I.M.H., N.W., L.V., A.K.N.I., and D.J.L. drafted supplementary notes and materials.

W.B., Y.Y., E.K.I-P., K.E.A., L.T.J., A.P.A., K.K., R.N., A.K.N.I., D.J.L., L.F., and E.W. were involved in reviewing drafts and editing.

All co-authors read, commented on, and agreed upon the submitted manuscript.

## COMPETING INTERESTS

The authors declare no competing interests

**Extended Data Figure 1.**
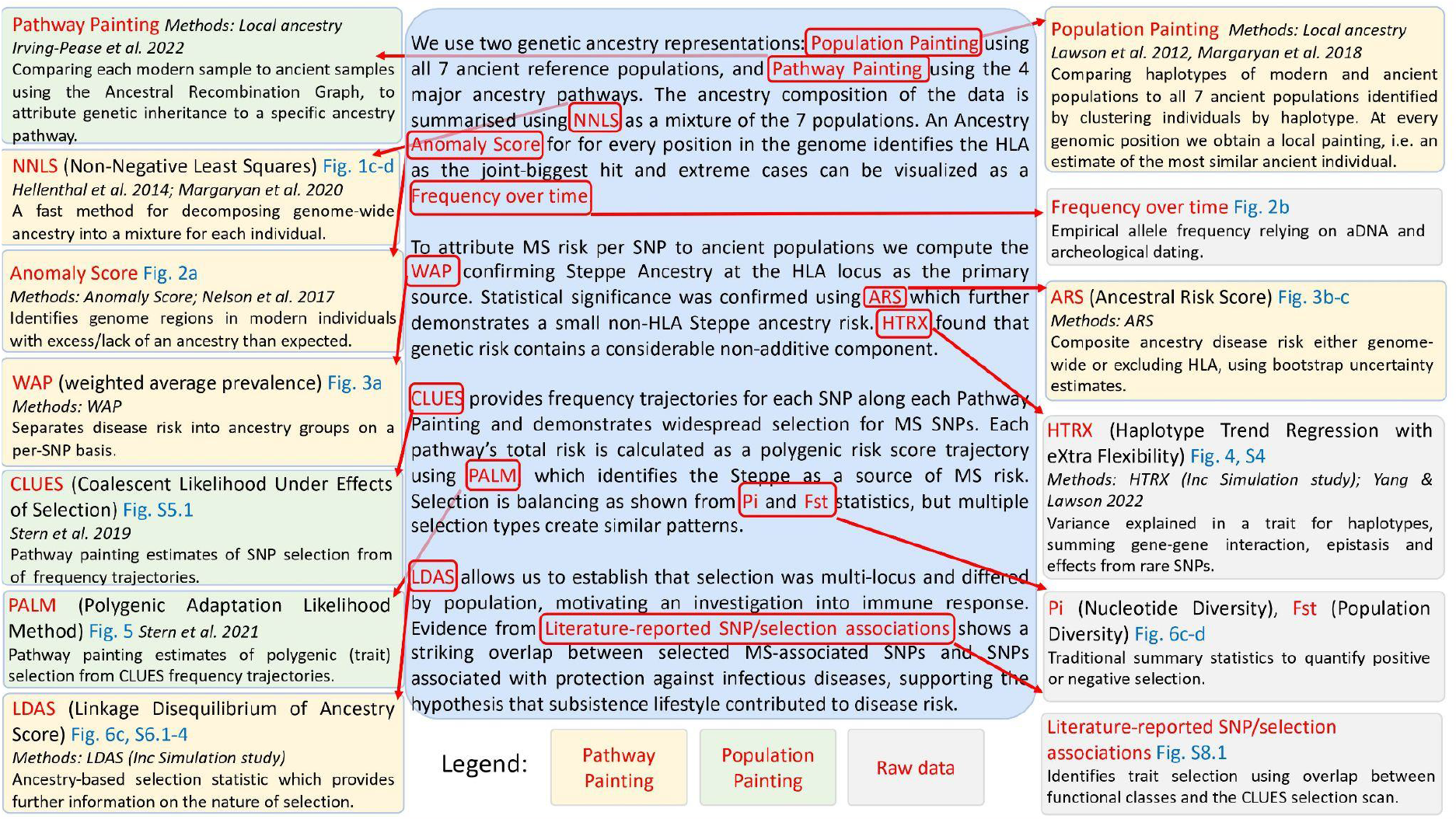
Methods map detailing datasets used, methods, and statistics. A narrative of the evidence used is provided in the centre, with boxes on each side detailing the methods used. Boxes are coloured by the dataset used.

**Extended Data Figure 2.**
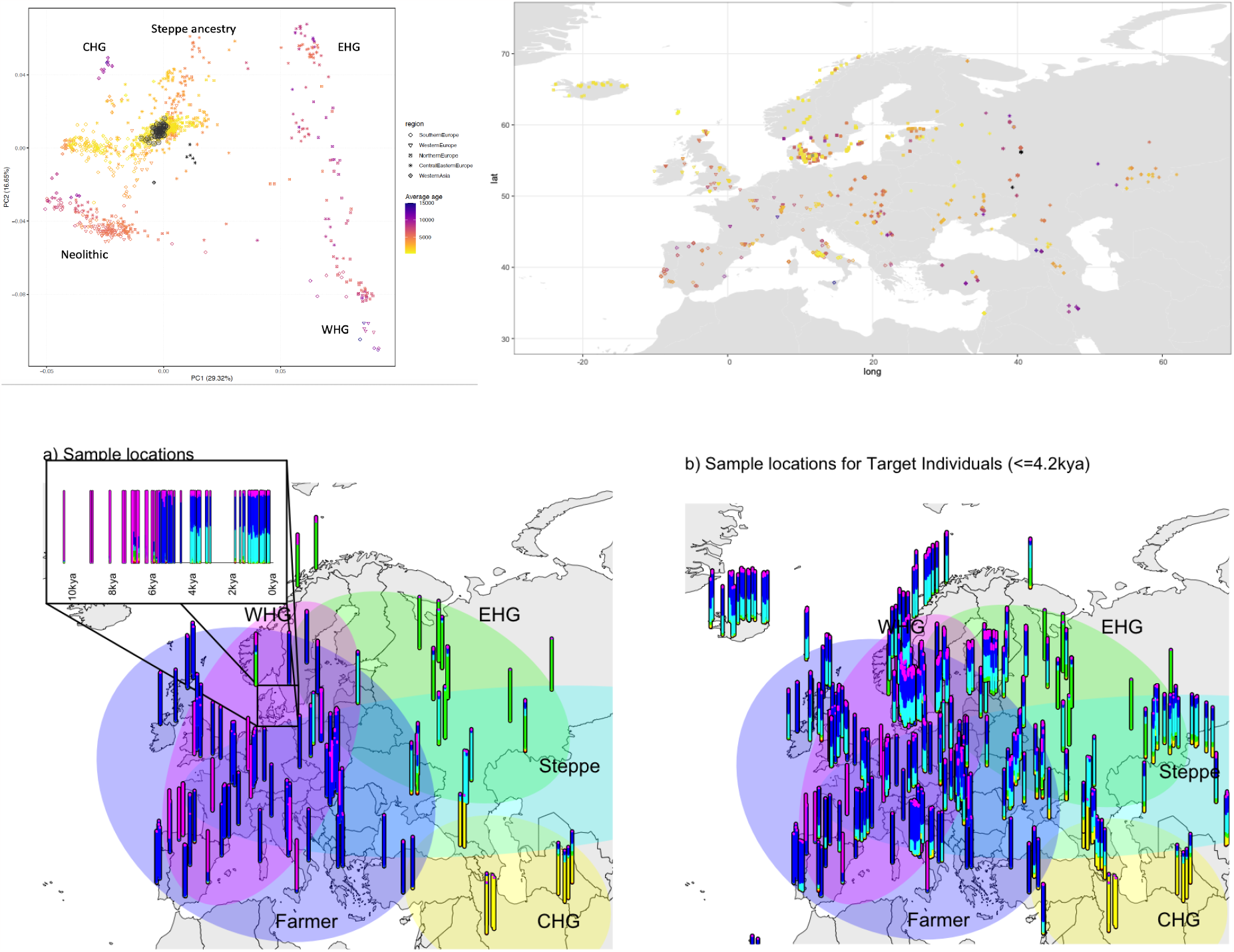
Ancient sample PCA, map, ancestry proportions through time for samples in Denmark. (1) PC1 vs PC2 of the filtered Western Eurasian ancient samples included in this study. Black circled points are Danish Medieval and post-Medieval samples published here for the first time. Major component ancestry locations are labelled. (2) Map of ancient filtered Western Eurasian ancient samples included in this study (3a) Map of reference data and time transect of Denmark as in Figure 1. (3b) More recent ancient data (samples < 4,200 years ago) not used as reference, showing the clines of the main ancestry components from (3a).

**Extended Data Figure 3.**
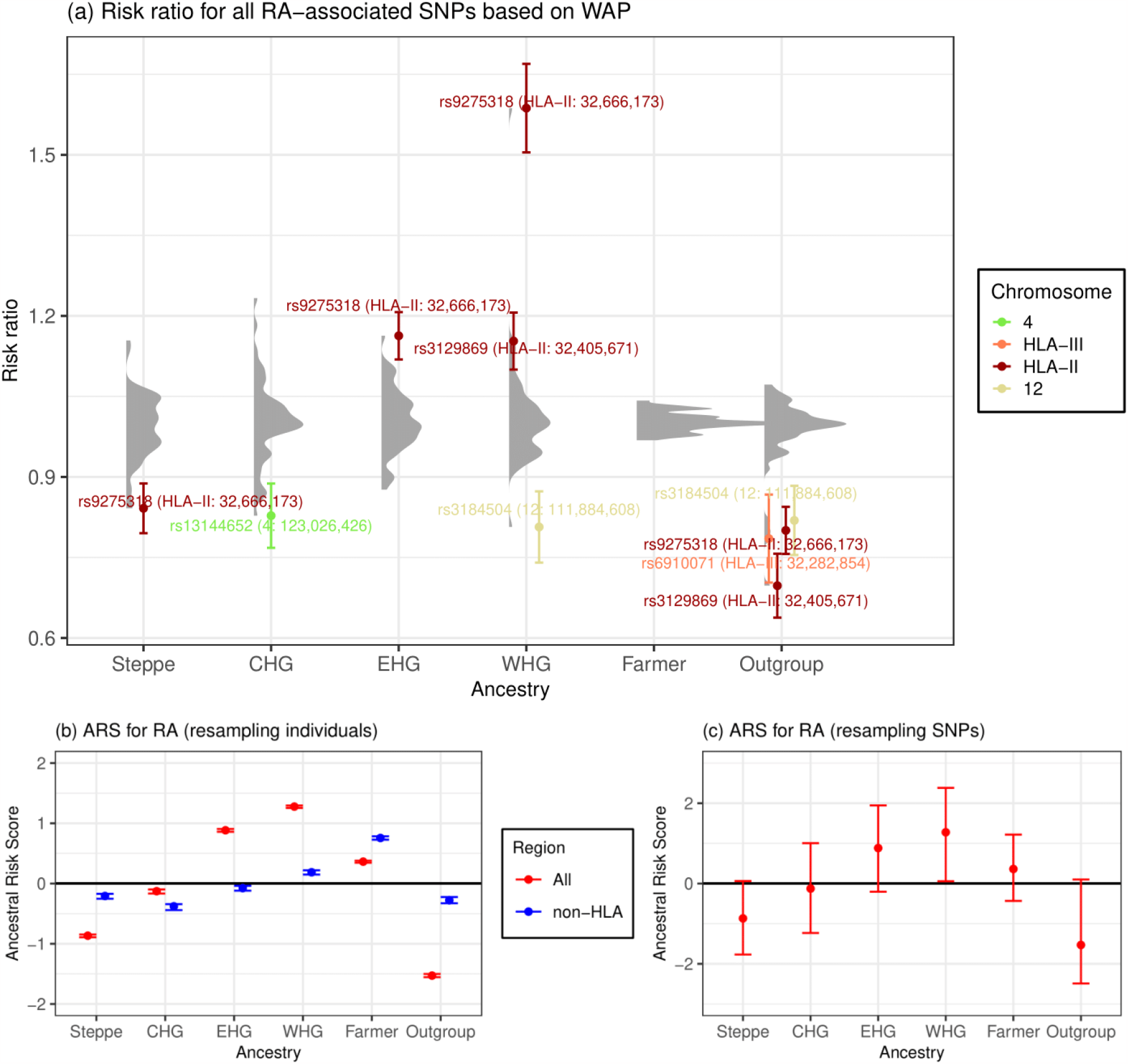
Associations between local ancestry at fine-mapped RA SNPs and RA in a modern population. a) Risk ratio of SNPs for RA based on weighted average prevalence (WAP; see Methods), when decomposed by inferred ancestry. A mean and standard deviation are calculated for each ancestry based on bootstrap resampling, for each chromosome. The distribution of risk ratios at each ancestry is shown as a raincloud plot. SNPs significant at the 1% level are shown individually, coloured by chromosome or HLA region, and those with risk ratio >1.1 or <0.9 are annotated with rsID, HLA region and position (build GRCh37/hg19). b-c) Genome-wide Ancestral Risk Scores (ARS, see Methods) for RA. Confidence intervals are estimated by either bootstrapping over individuals (b, which can be interpreted as testing power to reject a null hypothesis of no association between RA and ancestry) and bootstrapping over SNPs (c, which can be interpreted as testing whether ancestry is associated with RA genome-wide). We show results for all associated SNPs (red) and non-HLA SNPs only (blue) when bootstrapping over individuals.

**Extended Data Figure 4.**
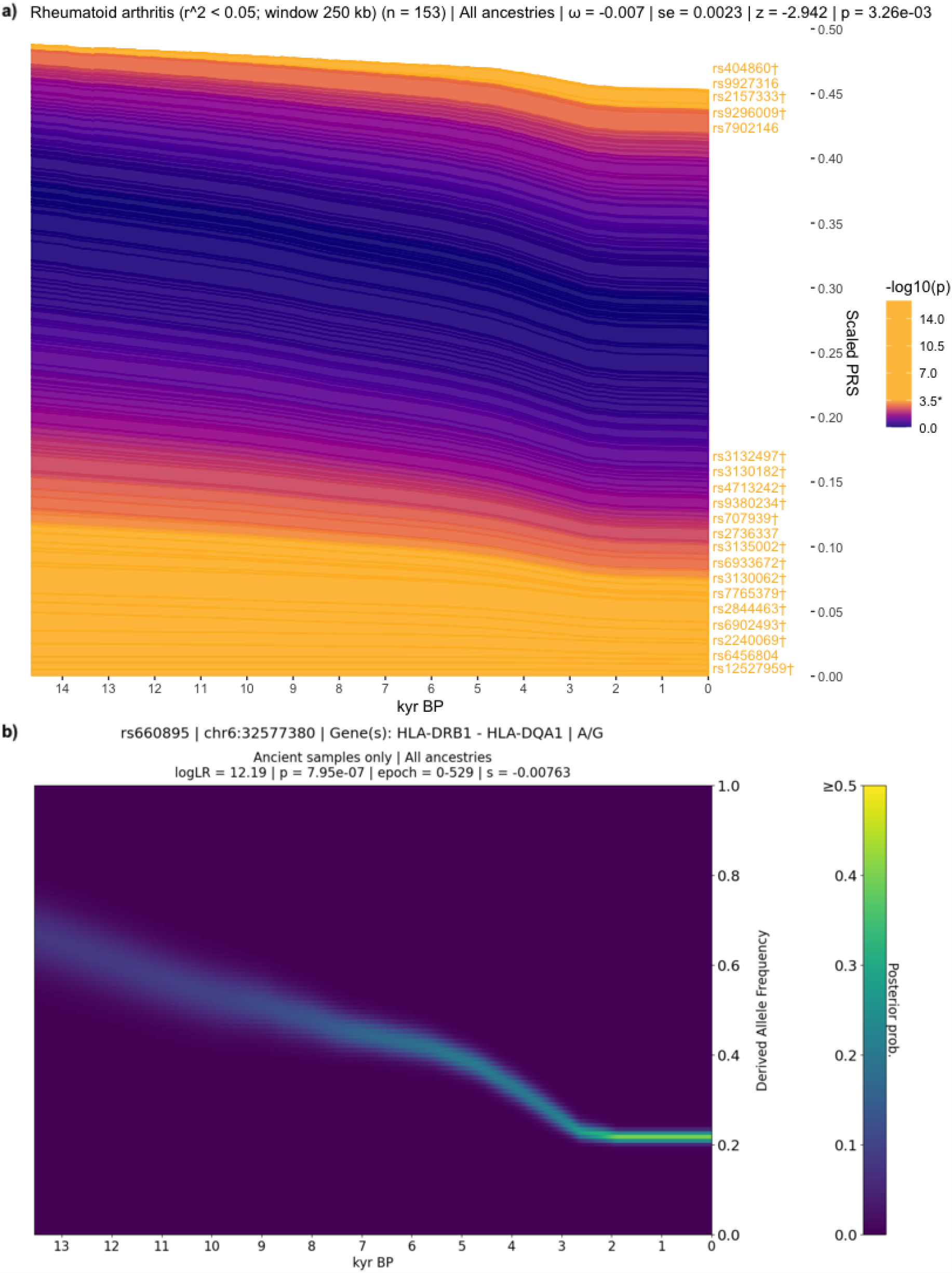
Evidence for selection on RA-associated SNPs. a) Stacked line plot of the pan-ancestry PALM analysis for RA, showing the contribution of SNPs to disease risk over time. SNPs are shown as stacked lines, the width of each line being proportional to the population frequency of the positive risk allele, weighted by its effect size. When a line widens over time the positive risk allele has increased in frequency, and vice versa. SNPs are sorted by the magnitude and direction of selection, with positively selected SNPs at the top, negatively selected SNPs at the bottom, and neutral SNPs in the middle. SNPs are coloured by their corresponding p-value in a single locus selection test. The asterisk marks the Bonferroni corrected significance threshold, and nominally significant SNPs are shown in yellow and labelled by their rsIDs. SNPs marked with the dagger symbol are located in the HLA locus. The Y-axis shows the scaled average polygenic risk score (PRS) in the population, ranging from 0 to 1, with 1 corresponding to the maximum possible average PRS (i.e. when all individuals in the population are homozygous for all positive risk alleles) and the X-axis shows time in units of thousands of years before present (kyr BP). b) Posterior likelihood trajectory for rs660895, tagging HLA-DRB1*04:01, inferred by CLUES.

**Extended Data Figure 5.**
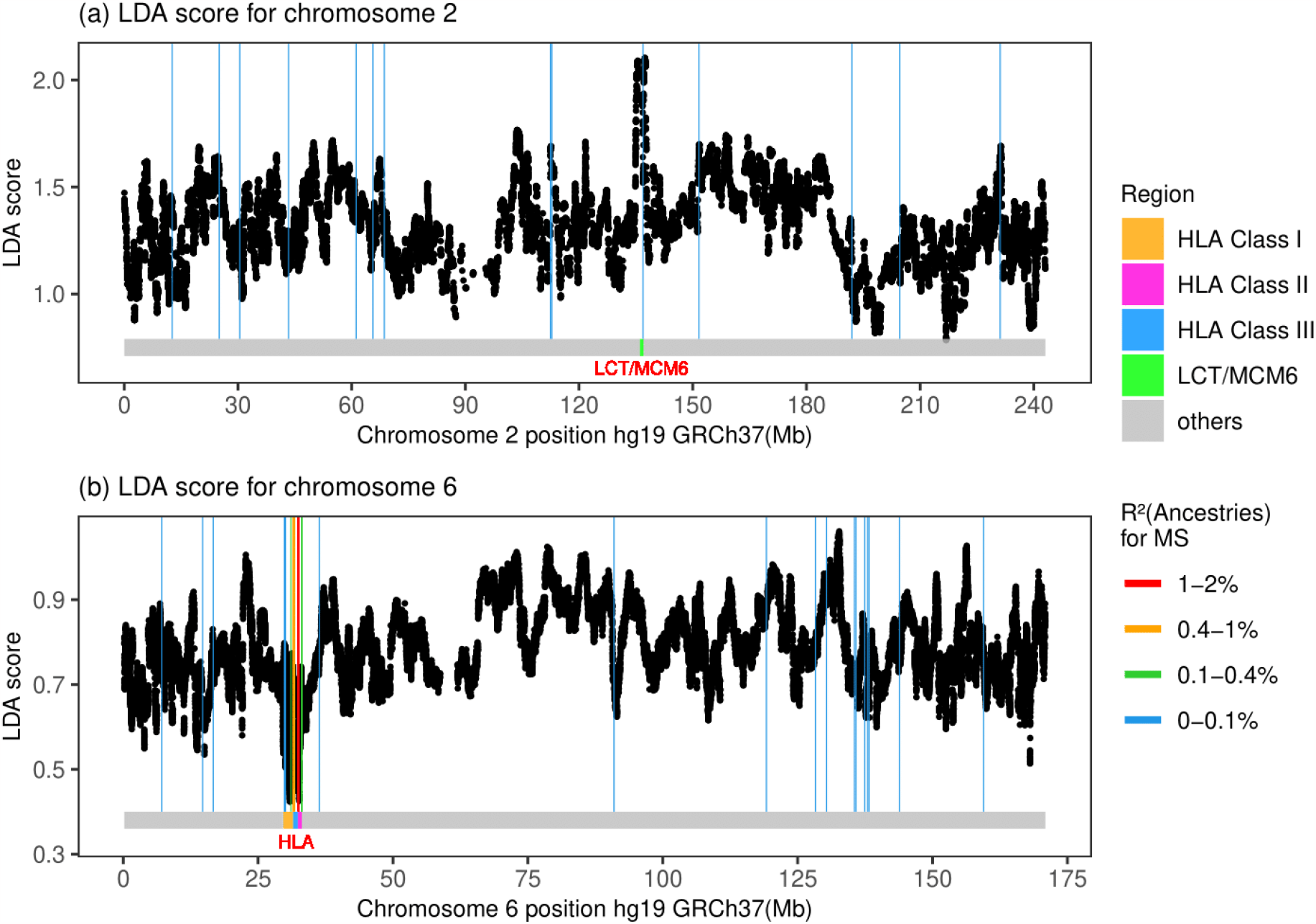
LDAS on chromosome 6 and 2. LDA score is a) high in the LCT/MCM6 region while is b) low in the HLA region.

**Extended Data Figure 6.**
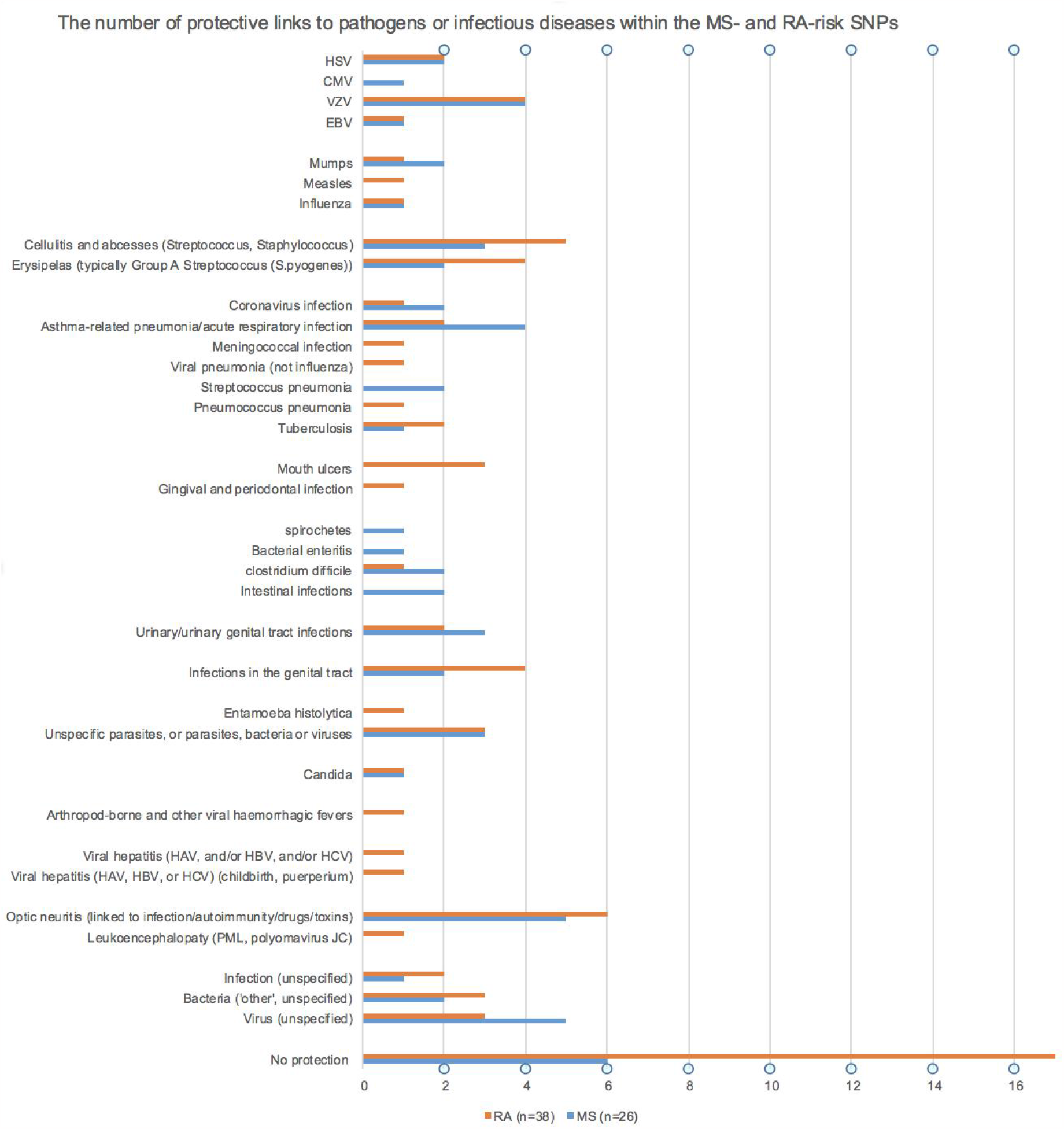
The number of protective associations with pathogens or infectious diseases for the MS- and RA-associated selected SNPs. The number of protective associations to specific pathogens and/or diseases associated with the MS- and RA-SNPs that showed statistically significant evidence for selection using CLUES. One SNP can have a link to more than one pathogen and/or disease (see ST11 and ST12 for details on each SNP). Eight and twenty SNPs had no detectable links to any pathogen or infectious disease in the MS and RA SNP sets, respectively.

## METHODS

### Data Generation Overview

In order to examine variants associated with phenotypes backwards in time, we assembled a large ancient DNA dataset. Here we present new genomic data from 86 ancient individuals from Medieval and post-Medieval periods from Denmark (Extended Data Figure 2, Supplementary Note 1, ST1). The samples range in age from around the 11th to the 18th century. We extracted ancient DNA from tooth cementum or petrous bone and shotgun sequenced the 86 genomes to a depth of genomic coverage ranging from 0.02 X to 1.6 X (mean = 0.39 X and median = 0.27 X). The genomes of the new 86 individuals were imputed using the 1,000 Genomes phased data as a reference panel by an imputation method designed for low coverage genomes (GLIMPSE, ^53^), and we also imputed 1,664 ancient genomes presented in the accompanying study “Population Genomics of Stone Age Eurasia” ^2^.

Depending on the specific data quality requirements for the downstream analyses, we filtered out samples with poor coverage, variant sites with low minor allele frequency (MAF) and with low imputation quality (average genotype probability < 0.98). Our dataset of ancient individuals spans approximately 15,000 years across Eurasia (Extended Data Figure 2).

### Ancient data DNA extraction and library preparation

Laboratory work was conducted in the dedicated ancient DNA clean-room facilities at the Lundbeck Foundation GeoGenetics Centre (Globe Institute, University of Copenhagen). A total of 86 Medieval and post-Medieval human samples from Denmark (ST2) were processed using semi-automated procedures. Each sample was processed in parallel. For each extract non USER-treated and USER-treated (NEB) libraries were built ^54^. All libraries were sequenced on the NovaSeq6000 instrument at the GeoGenetics Sequencing Core, Copenhagen, using S4 200 cycles kits version 1.5. A more detailed description of DNA extraction and library preparation can be found in Supplementary Note 1.

### Basic bioinformatics

The sequencing data was demultiplexed using the Illumina software BCL Convert (https://emea.support.illumina.com/sequencing/sequencing_software/bcl-convert.html, Illumina Inc.). Adapter sequences were trimmed and overlapping reads were collapsed using AdapterRemoval (2.2.4 ^55^). Single-end collapsed reads of at least 30bp and paired-end reads were mapped to the human reference genome build 37 using BWA (0.7.17 ^56^) with seeding disabled to allow for higher sensitivity. Paired- and single-end reads for each library and lane were merged, and duplicates were marked using Picard MarkDuplicates (2.18.26, http://picard.sourceforge.net) with a pixel distance of 12000. Read depth and coverage were determined using samtools (1.10 ^57^) with the all sites used in the calculation (-a). Data was then merged to sample level and duplicates were marked again.

### DNA authentication

In order to determine the authenticity of the ancient reads, post-mortem DNA damage patterns were quantified using mapDamage2.0 ^58^. Next, two different methods were used to estimate the levels of contamination. Firstly, we applied ContamMix in order to quantify the fraction of exogenous reads in the mitochondrial reads by comparing the mtDNA consensus genome to possible contaminant genomes ^59^. The consensus was constructed using an in-house perl script that used sites with at least 5 X coverage, and bases were only called if observed in at least 70% of reads covering the site. Lastly, we applied ANGSD (0.931 ^60^) to estimate nuclear contamination by quantifying heterozygosity on the X chromosome in males. Both contamination estimates only used filtered reads with a base quality of ≥20 and mapping quality of ≥30.

### Imputation

We combined the 86 newly sequenced Medieval and post-Medieval Danish individuals with 1,664 previously published ancient genomes ^2^. We then excluded individuals showing: contamination (more than 5%); low autosomal coverage (less than 0.1 X); low genome-wide average imputation genotype probability (less than 0.98), and we chose the best quality sample in a close relative pair (first or second degree relative). A total of 1,557 individuals passed all filters, and were used in downstream analyses. We restricted the analysis to SNPs with imputation INFO score ≥ 0.5 and MAF ≥ 0.05.

### Kinship analysis and uniparental haplogroup inferences

READ ^61^ was used to detect the degree of relatedness between pairs of individuals.

The mtDNA haplogroups of the Medieval and post-Medieval individuals were assigned using HaploGrep2 ^62^. Y chromosome haplogroup assignment was inferred following the workflow already published ^63^. More details can be found in Supplementary Note 2.

### Standard Population genetic analyses

The main population-genetics approach we base our inference on is Population-based painting (detailed below). However, to robustly understand population structure, we applied other standard techniques. Firstly, we used principal component analysis (PCA) (Extended Data Figure 2) to investigate the overall population structure of the dataset. We used PLINK ^64^, excluding SNPs with MAF < 0.05 in the imputed panel. Based on 1,210 ancient western Eurasia imputed genomes, the Medieval and post-Medieval samples cluster very close to each other, displaying a relatively low genetic variability and situated within the genetic variability observed in the post-Bronze Age western Eurasian populations.

We then used two additional standard methods to estimate ancestry components in our ancient samples. Firstly, we used model-based clustering (ADMIXTURE) ^65^ (Supplementary Note 1, Figure S1.1) on a subset of 826,248 SNPs. Secondly, we used qpAdm ^66^ (Supplementary Note 1 Figure S1.2 and Table S1.1) with a reference panel of three genetic ancestries (WHG, ANA, and Steppe) on the same 826,248 SNPs. We performed qpAdm applying the option “allsnps: YES” and a set of 7 outgroups was used as “right populations”: Siberia_UpperPaleolithic_UstIshim, Siberia_UpperPaleolithic_Yana, Russia_UpperPaleolithic_Sunghir, Switzerland_Mesolithic, Iran_Neolithic, Siberia_Neolithic, USA_Beringia. We set a minimum threshold of 100,000 SNPs and only results with p > 0.05 only have been considered.

### Population painting

Our main analysis uses chromosome painting ^67^ with a panel of 6 ancient ancestries (as on the UK Biobank, see below). This allows fine-scale estimation of ancestry as a function of those populations. We ran chromosome painting on all ancient individuals not in the reference panel, using a reference panel of ancient donors grouped into populations to represent specific ancestries: Western Hunter-Gatherer (WHG), Eastern Hunter-Gatherer (EHG), Caucasus Hunter-Gatherer (CHG), Farmer (ANA+Neolithic), Steppe, and African (method described in ^14^). Painting followed the pipeline of ^68^ based on GLOBETROTTER ^69^, with admixture proportions estimated using Non-Negative Least squares (NNLS). NNLS explains the genome-wide haplotype matches of an individual as a mixture of the genome-wide haplotype-matches of the reference populations. This setup allows both the reference panel and any additional samples (i.e. modern) to be described using these 6 ancestries (Figure 1).

We then painted individuals born in Denmark of a typical ancestry (typical based on density-based clustering of the first 18 PCs, ^2^). The reference panel used for chromosome painting was designed to capture the various components of European ancestry only, and so we urge caution in interpreting these results for non-European samples.

This dataset provides the opportunity to study the population history of Denmark from the Mesolithic to the post-Medieval period, covering around 10,000 years, which can be considered a typical Northern European population. Our results clearly demonstrate the impact of previously described demographic events, including the influx of Neolithic Farmer ancestry ∼9,000 years ago and Steppe ancestry ∼5,000 years ago ^10,12^. We highlight genetic continuity from the Bronze Age to the post-Medieval period (Supplementary Note 1 Figure S1.1), although *qpAdm* detected a small increase in the Farmer component during the Viking Age (Supplementary Note 1 Figure S1.2 and Table S1.1), while the Medieval period marked a time of increased genetic diversity, likely reflecting increased mobility across Europe. This genetic continuity is further confirmed by the haplogroups identified in the uniparental genetic makers (Supplementary Note 2). Together, these results suggest that after the Bronze Age Steppe migration there was no other major gene flow into Denmark from populations with significantly different Neolithic and Bronze Age ancestry compositions, and therefore no changes in these ancestry components in the Danish population.

### Local ancestry from Population painting

Chromosome Painting provides an estimate of the probability that an individual from each reference population is the closest match to the target individual at every position in the genome. This provides our first estimate of local ancestry from ^2^: the population of the first reference individual to coalesce with the target individual, as estimated by Chromopainter ^67^. This was estimated for all “White British” individuals in the UK Biobank, using the population painting reference panel described above. We refer to this henceforth as “local ancestry”, though note that the closest relative in the sample may not represent ancestry in the conventional sense.

### Pathway painting

An alternative approach is to identify which of the four major ancestry pathways (ANA Farmer, CHG, EHG, WHG) each position in the genome best matches to. This has the advantage of not forcing haplotypes to choose between “Steppe” ancestry and its ancestors, but the disadvantage of being more complex to interpret. To do this, we modelled ancestry path labels in GBR, FIN and TSI 1000G populations ^70^ and 1015 ancient genomes generated using a neural network to assign ancestry paths based on a sample’s nearest neighbours at the first five informative nodes of a marginal tree sequence, where an informative node is defined as one which has at least one leaf from the reference set of ancient samples described above (^14^ Supplementary Note S1c). We refer to this henceforth as “ancestry path labels”.

### SNP associations

We aimed to generate SNP associations from previous studies for each phenotype in a consistent approach. To generate a list of SNPs associated with multiple sclerosis (MS) and rheumatoid arthritis (RA), we used two approaches: in the first, we downloaded fine-mapped SNPs from previous association studies. For each fine-mapped SNP, if the SNP did not have an ancestry path label, we found the SNP in highest LD that did, with a minimum threshold of *r*^2^ ≥ 0.7 in the GBR, FIN and TSI 1000G populations using LDLinkR ^71^. The final SNPs used for each phenotype can be found in ST4 (MS), and ST5 (RA).

For MS, we used data from ^4^. For non-MHC SNPs, we used the “discovery” SNPs with P(joined) and OR(joined) generated in the replication phase. For MHC variants, we searched the literature for the reported HLA alleles and amino-acid polymorphisms (ST3). In total, we generated 205 SNPs which were either fine-mapped or in high LD with a fine-mapped SNP (15 MHC, 190 non-MHC).

For RA, we downloaded 57 genome-wide significant non-MHC SNPs for seropositive RA in Europeans ^72^. We retrieved MHC associations separately (^73^, with associated ORs and p-values from ^74^). In total, we generated 51 SNPs which were either fine-mapped or in high LD with a fine-mapped SNP (3 MHC, 48 non-MHC).

Secondly, because we could not always find LD proxies for fine-mapped SNPs that were present in our ancestry path labels dataset, we found that we were losing significant signal from the HLA, therefore we generated a second set of SNP associations. We downloaded full summary statistics for each disease (MS: ^4^; RA: ^75^), restricted to sites present in the ancestry path labels dataset, and ran PLINK’s (v1.90b4.4 ^76^) clump method (parameters: --clump-p1 5e-8 --clump-r2 0.05 --clump-kb 250 as in ^77^) using LD in the GBR, FIN and TSI 1000G populations ^70^ to extract genome-wide significant independent SNPs.

In the main text we report results for the first set of SNPs (“fine-mapped”) for analyses involving local ancestry in modern data, and the second set of SNPs (“pruned”) for analyses involving polygenic measures of selection (CLUES/PALM).

### Anomaly Score: Regions of Unusual Ancestry

To assess which regions of ancestry were unusual, we converted the ancestry estimates to Z-scores by standardizing to the genome-wide mean and standard deviation. Specifically, let *A*(*i, j, j*) denote the probability of the *k*th ancestry (*k* = 1,…, *K*) at the *j*th SNP (*j* = 1,…, *J*) of a chromosome for the *i*th individual (*i* = 1,…, *N*). We first computed the mean painting for each SNP, 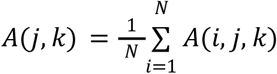. From this, we estimated a location parameter μ_*k*_ and scale parameter σ_*k*_ using a block-median approach. Specifically, we partitioned the genome into 0.5Mb regions, and within each, computed the mean and standard deviation of the ancestry. The parameter estimates are then the median values over the whole genome. We then computed an anomaly score for each SNP for each ancestry Z(*j, k*) = (*A*(*j, k*) −μ_*k*_)/σ_*k*_. This is the normal-distribution approximation to the Poisson-binomial score for excess ancestry, for which a detailed simulation study is presented in ^78^.

To arrive at an anomaly score for each SNP aggregated over all ancestries, we also had to account for correlations in the ancestry paintings. Instead of scaling each ancestry deviation *A*^*\**^ (*j, k*) = *A*(*j, k*) − μ_*k*_ by its standard deviation, we instead “whitened” them, i.e. rotated the data to have an independent signal. Let 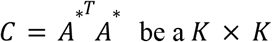 be a *K* × *K* covariance matrix, and let *C*^−1 *T*^ = *UDV*^*T*^ be its Singular Value Decomposition. Then *W* = *UD*^½^ is the whitening matrix from which *Z* = *A*^*^ *W* are normally distributed with covariance matrix diag(1) under the null hypothesis that *A*^*^ is normally distributed with mean 0 and unknown covariance Σ. The “ancestry anomaly score” test statistic for each SNP is *t* 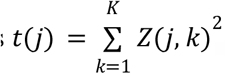, which is chi-squared distributed with *K* degrees of freedom under the null, and we reported p-values from this.

To test for gene enrichment, we formed a list of all SNPs reaching genome-wide significance (*p* < 5^−8^) and using the R package *gprofiler2*^79^ converted these to a unique list of genes. We then used *gost* to perform an enrichment test for each GO term, for which we used default p-value correction via the *g:Profiler SCS* method. This is an empirical correction based on performing random lookups of the same number of genes under the null, to control the error rate and ensure that 95% of reported categories (at p = 0.05) are correct.

### Allele Frequency over Time

To investigate how effect allele frequencies have changed over time, we extracted high effect alleles for each phenotype from the ancient data. We excluded all non-Eurasian samples, grouped them by “groupLabel”, excluded any group with fewer than 4 samples, and coloured points by ancestry proportion according to genome-wide NNLS based on chromosome painting (above).

### Weighted Average Prevalence (WAP)

In order to understand whether risk-conferring haplotypes evolved in the Steppe population, or in a pre- or post-dating population, we developed a statistic that could account for the origin of risk to be identified with multiple ancestry groups, which do not have to be the same set for each SNP.

We first applied k-means clustering to the dosage of each ancestry for each associated SNP and investigated the dosage distribution of clusters with significantly higher MS prevalence. For the target SNPs, the elbow method ^80^ suggested selecting around 5-7 clusters, of which we chose 6. After performing the k-means cluster analysis, we calculated the average probability for each ancestry for case individuals. Furthermore, we calculated the prevalence of MS in each cluster, and performed a one-sample t-test to investigate whether it differs from the overall MS prevalence (0.487%). This tests whether any particular combinations of ancestry are associated with the phenotype at a SNP. Clusters with high MS risk ratios have high Steppe components (Supplementary Information Figure S3.3), leading to the conclusion that Steppe ancestry alone is driving this signal.

We can then compute the Weighted Average Prevalence (WAP), which summarises these results into the ancestries. For the *j*th SNP, let 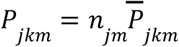 denote the sum of the *k*th ancestry probabilities of all the individuals in the *m*th cluster (*k, m* = 1,…, 6), where *n*_*jm*_ is the cluster size of the *m*th cluster. Let π_*jm*_ denote the prevalence of MS in the *m*th cluster, the weighted average prevalence for the *k*th ancestry is defined as:

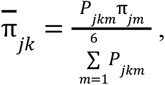

where *P*_*jkm*_ is defined as the weight for each cluster.

The standard deviation of 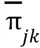 is computed as 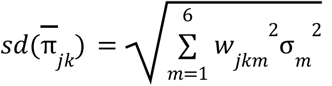, where 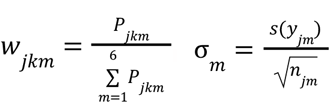 and *s*(*y*_*jm*_) is the standard deviation of the outcome for the individuals in the *m*th cluster. We also test the hypothesis that 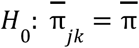 against 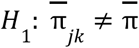, and compute the p-value as 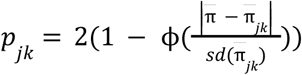.

For each ancestry, WAP measures the association of that ancestry with MS risk across all clusters. To make a clear comparison, we calculated the risk ratio (compared to the overall MS prevalence) for each ancestry at each SNP, and assigned a mean and confidence interval for the risk ratios of each ancestry at each chromosome (Figure 3, Extended Data Figure 3).

### PCA/UMAP of WAP/Average Dosage

To sort risk-associated SNPs into ancestry patterns according to that risk, we performed PCA on the average ancestry probability and WAP at each MS-associated SNP (Supplementary Information Figure S3.4). The former shows that all of the HLA SNPs except three from HLA class II and III have much larger Outgroup components compared with the others. The latter analysis indicates a strong association between Steppe and MS risk. Also, Outgroup ancestry at rs10914539 from chromosome 1 exceptionally reduces the incidence of MS, while Outgroup ancestry at rs771767 (chromosome 3) and rs137956 (chromosome 22) significantly boosts MS risk.

### Ancestral Risk Score (ARS)

To assign risk to ancient ancestries by computing the equivalent of a polygenic score for each, we followed methods developed in ^14^. We calculated the effect allele painting frequency for a given ancestry *f*_*{anc,i}*_ for SNP *i* using the formula:

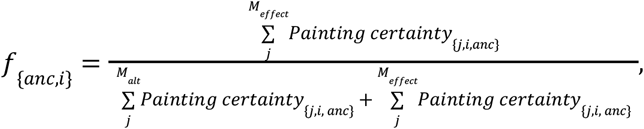

where there are *M*_*effect*_ individuals homozygous for the effect allele, *M*_*alt*_ individuals homozygous for the other allele, and 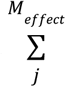 is the sum of the painting probabilities for that ancestry *anc* in individuals homozygous for the effect allele at SNP *i*. This can be interpreted as an estimate of an ancestral contribution to effect allele frequency in a modern population. The per-SNP painting frequencies can be found in ST4, ST5, and ST6.

To calculate the ancestral risk score (ARS) we summed over all *I* pruned SNPs in an additive model:

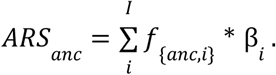

We then ran a transformation step as in ^81^, centering results around the ancestral mean (i.e. all ancestries) and reporting as a Z-score. To obtain 95% confidence intervals, we ran an accelerated bootstrap over loci, which accounts for the skew of data to better estimate confidence intervals ^82^.

### GWAS of Ancestry and Genotypes

The total variance of a trait explained by genotypes (SNP values), ancestry, and haplotypes (described below) is a measure of how well each captures the causal factors driving that trait. We therefore computed the variance explained for each data type in a “head-to-head” comparison, either at specific SNPs or SNP sets. In this section, we describe the model and covariates accounted for.

We used the UK Biobank to fit GWAS models for local ancestry values and genotype values separately, using only SNPs known to be associated with the phenotype (“fine-mapped” SNPs). We used the following phenotype codes for each phenotype: MS: Data-Field 131043; RA: Data-Field 131849 (seropositive).

Let Y_*i*_ denote the phenotype status for the *i*th individual (*i* = 1,…, 399998), which takes value 1 for a case and 0 for control, and let π_*i*_ = *Pr*(*Y*_*i*_ = 1) denote the probability that this individual is a case. Let *X*_*ijk*_ denote the *k*th ancestry probability (*k* = 1,…, *K*) for the *j*th SNP (*j* = 1,…, 205) of the *i*th individual. *C*_*ic*_ is the *c*th predictor (*c* = 1,…, *N*_*c*_) for the *i*th individual. We used the following logistic regression model for GWAS, which assumes the effects of alleles are additive:

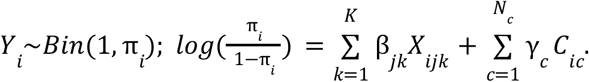

We used *N*_*c*_ = 20 predictors in the GWAS models, including sex, age and the first 18 PCs, which are sufficient to capture most of the population structure in the UK Biobank ^83^.

First, we built the model with *K* = 1. By using only one ancestry probability in each model, we aimed to find the statistical significance of each SNP under each ancestry. Then, we built the model with *K* = 5, i.e. using all 6 local ancestry probabilities which sum to 1. We calculated the variance explained by each SNP by summing up the variance explained by *X*_*ijk*_ (k = 1,…,5).

We considered fitting the multivariate models by using all the SNPs as covariates. However, the dataset only contains 1,982 cases. Even though only one ancestry is included, the multivariate model contains 191 predictors, which could result in overfitting problems. Therefore, the GWAS models are preferred over multivariate models.

We also fitted a logistic regression model for GWAS using the genotype data as follows:

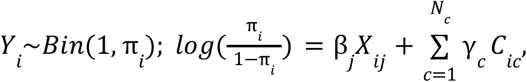

where *X*_*ijk*_ ∈ {0, 1, 2} denotes the number of copies of the reference allele of the jth SNP (*j* = 1,…, 205) that the *i*th individual has, and *C*_*ic*_ (*c* = 1,…, *N*_*c*_) denotes the covariates including age, sex and first 18 PCs for the ith individual, where *N*_*c*_ = 20. Due to the UK Biobank being underpowered compared to the case-control study from which these SNPs were found, the only statistically significant (*p* < 10^−5^) association is for the HLA class II SNP tagging HLA-DRB1*15:01.

### GWAS comparison for trait-associated SNPs

In this section, we describe how we moved from associations between SNPs (either genotype values or ancestry) and a trait, to total variance explained.

We compared the variance explained by SNPs from the GWAS model using the painting data (all 6 local ancestry probabilities; the 7th is a linear combination of the first 6) with that from the GWAS model using the genotype data. McFadden’s pseudo-R-squared measure ^84^ is widely used for estimating the variance explained by the logistic regression models. McFadden’s pseudo-R-squared is defined as

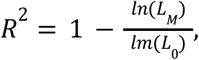

where *L*_*M*_ and *L*_0_ are the likelihoods for the fitted and the null model, respectively. Taking overfitting into account, we use the adjusted McFadden’s pseudo-R-squared by penalizing the number of predictors:

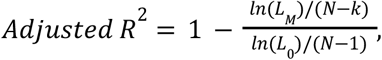

where N is the sample size and k is the number of predictors.

Specifically, *R*^2^ (*SNPs*) is calculated as the extra variance in addition to sex, age and 18 PCs that can be explained by SNPs:

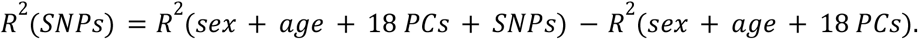

Notably, two SNPs stand out for explaining much larger variance than others when fitting the GWAS model using the genotype data, but overall more SNPs from GWAS painting explain more than 0.1% variance, which indicates the painting data are probably more efficient for estimating the effect sizes of SNPs and detecting significant SNPs. Also, some SNPs from GWAS models using painting data explain almost the same amount of variance, suggesting that these SNPs consist of very similar ancestries.

### Haplotype Trend Regression with eXtra flexibility (HTRX)

Ancestry is a strong predictor of MS, but we wanted to understand whether it was tagging some causal factor that was not in our genetic data, or whether it was tagging either interactions or rare SNPs. To address this, we propose Haplotype Trend Regression with eXtra flexibility (HTRX), which searches for haplotype patterns that include single SNPs and non-contiguous haplotypes. HTRX is an association between a template of *n* SNPs and a phenotype. A template gives a value for each SNP taking values of “0” or “1”, reflecting whether the reference allele of each SNP is present or absent, or an “X” meaning either value is allowed. For example, haplotype “1X0” corresponds to a 3-SNP haplotype where the first SNP is the alternative allele and the third SNP is the reference allele, while the second SNP can be either the reference or the alternative allele. Therefore, haplotype “1X0” is essentially only a 2-SNP haplotype.

To examine the association between a haplotype and a binary phenotype, we replace the genotype term with a haplotype from the standard GWAS model:

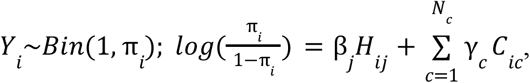

where *H* denotes the *j*th haplotype probability for the *i*th individual:

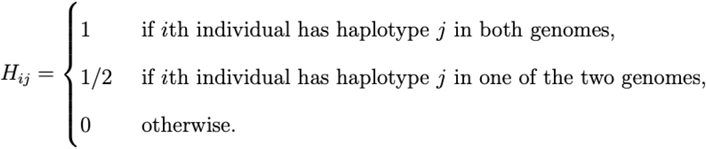

HTRX can identify gene-gene interactions, and is superior to HTR not only because it can extract combinations of significant SNPs within a region, leading to improved predictive performance, but the haplotypes are more interpretable as multi-SNP haplotypes are only reported when they lead to increased predictive performance.

### HTRX Model selection procedure for shorter haplotypes

Fitting HTRX models directly on the whole dataset can lead to significant overfitting, especially when the number of SNPs increases. When overfitting occurs, the models experience poorer predictive accuracy against unseen data. Further, HTRX introduces an enormous model space which must be searched.

To address these problems, we implemented a two-step procedure:

Step 1: Select candidate models. This step aims at addressing the model search problem by obtaining a set of models more diverse than traditional bootstrap resampling ^85^.

1. Randomly sample a subset (50%) of data. Specifically, when the outcome is binary, stratified sampling is used to ensure the subset has approximately the same proportion of cases and controls as the whole data.
2. Start from a model with fixed covariates (18 PCs, sex and age), and perform forward regression on the subset, i.e. iteratively choose a feature (in addition to the fixed covariates) to add whose inclusion enables the model to explain the largest variance, and select *s* models with the lowest Bayesian Information Criterion (BIC) ^86^ to enter the candidate model pool.
3. Repeat (1)-(2) *B* times, and select all the different models in the candidate model pool as the candidate models.

Step 2: Select the best model using 10-fold cross-validation.

1. Randomly split the whole data into 10 groups with approximately equal sizes, using stratified sampling when the outcome is binary.

(2)In each of the 10 folds, use a different group as the test dataset, and take the remaining groups as the training dataset. Then, fit all the candidate models on the training dataset, and use these fitted models to compute the additional variance explained by features (out-of-sample *R*^2^) in the test dataset. Finally, select the candidate model with the biggest average out-of-sample *R*^2^ as the best model.

### HTRX Model selection procedure for longer haplotypes (Cumulative HTRX)

Longer haplotypes are important for discovering interactions. However, there are 3^*k*^ − 1 haplotypes in HTRX if the region contains *k* SNPs, making it unrealistic for regions with large numbers of SNPs. To address this issue, we proposed cumulative HTRX to control the number of haplotypes, which is also a two-step procedure.

Step 1: Extend haplotypes and select candidate models.

1. Randomly sample a subset (50%) of data, using stratified sampling when the outcome is binary. This subset is used for all the analysis in (2) and (3).
2. Start with *L* randomly chosen SNPs from the entire *k* SNPs, and keep the top *M* haplotypes that are chosen from the forward regression. Then add another SNP to the *M* haplotypes to create 3*M* + 2 haplotypes. There are 3*M* haplotypes obtained by adding “0”, “1” or “X” to the previous *M* haplotypes, as well as 2 bases of the added SNP, i.e. “XX…X0” and “XX…X1” (as “X” was implicitly used in the previous step). The top *M* haplotypes from them are then selected using forward regression. Repeat this process until obtaining *M* haplotypes which include *k* − 1 SNPs.

(3)Add the last SNP to create 3*M* + 2 haplotypes. Afterwards, start from a model with fixed covariates (18 PCs, sex and age), perform forward regression on the training set, and select *s* models with the lowest BIC to enter the candidate model pool.

(4)Repeat (1)-(3) *B* times, and select all the different models in the candidate model pool as the candidate models.

Step 2: Select the best model using 10-fold cross-validation, as described in “HTRX Model selection procedure for shorter haplotypes”.

We note that because the search procedure in Step 1(2) may miss some highly predictive haplotypes, cumulative HTRX acts as a lower bound on the variance explainable by HTRX.

As a model criticism, only common and highly predictive haplotypes (i.e. those with the greatest adjusted *R*^2^) are correctly identified, but the increased complexity of the search space of HTRX leads to haplotype subsets that are not significant on their own but are significant when interacting with other haplotype subsets being missed. This issue would be eased if we increase all the parameters *s, l, M* and *B* but with higher computational cost, or improve the search by optimizing the order of adding SNPs. This leads to a decreased certainty that the exact haplotypes proposed are “correct”, but together reinforces the inference that interaction is extremely important.

### Simulation Study for HTRX

To investigate how the total variance explained by HTRX compare to GWAS and HTR, we used a simulation study comparing:

1. linear models (denoted by “lm”) and generalized linear models with a logit link-function (denoted by “glm”);
2. models with or without actual interaction effects;
3. models with or without rare SNPs (frequency smaller than 5%);
4. remove or retain rare haplotypes when rare SNPs exist.

We started from creating the genotypes for 4 different SNPs *G*_*ijk*_ (*i* = 1,…, 100000 denotes the index of individuals, *j* = 1 (“1*XXX*”), 2 (“*X*1*XX*”), 3 (“*XX*1*X*”) *and* 4 (“*XXX*1”) represents the index of SNPs, and *q* = 1, 2 for two genomes as individuals are diploid). If no rare SNPs were included, we sampled the frequency *F*_*j*_ of these 4 SNPs from 5% to 95%; otherwise, we sampled the frequency of the first 2 SNPs from 2% to 5% (in practice, we obtained *F*_1_ = 2. 8%and *F*_2_ = 3. 1%under our seed) while the last 2 SNPs from 5% to 95%. For the *i*th individual, we sampled *G*_*ijq*_ ∼*Bin*(1, *F*) for the *q*th genome of the *j*th SNP, and took the average value of two genomes as the genotype for the *j*th SNP of the *i*th individual: 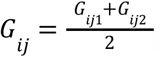. Based on the genotype data, we obtained the haplotype data for each individual, and we considered removing haplotypes rarer than 0.1% or not when rare SNPs were generated. In addition, we sampled 20 fixed covariates (including sex, age and 18 PCs) *C*_*ic*_, where *c* = 1,…, 20 from UK Biobank for 100000 individuals.

Next, we sampled the effect sizes of SNPs 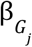 and covariates 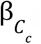, and normalized them by their standard deviations: 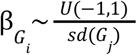 and 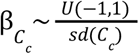 for each fixed *j* and *c*, respectively. When an interaction exists, we created a fixed effect size for haplotype “11XX” as twice the average absolute SNP effects: 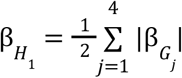 where *H*_1_ refers to “11XX”; otherwise, *H*_1_ = 0. Note that 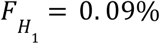 when rare SNPs are included.

Finally, we sampled the outcome based on the outcome score (for the *i*th individual)

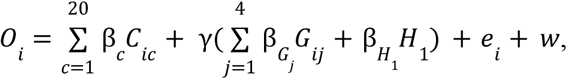

where γ is a scale factor for the effect sizes of SNPs and haplotype “11XX”, *e*_*i*_ ∼*N*(0, 0. 1) is the random error, and *w* is a fixed intercept term. For linear models, the outcome *Y*_*i*_ = *0*_*i*_ ; for generalized linear models, we sampled the outcome from the binomial distribution: *Y*_*i*_ ∼*Bin*(1, π_*i*_), where 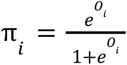 is the probability that the *i*th individual is a case.

As the simulation is intended to compare the variance explained by HTRX, HTR and SNPs (GWAS) in addition to fixed covariates, we tripled the effect sizes of SNPs and haplotype “11XX” (if an interaction exists) by setting γ = 3. In “glm”, to ensure a reasonable case prevalence (e.g. below 5%), we set *w* =− 7, which was also applied in “lm”.

We applied the procedure described in “HTRX Model selection procedure for shorter haplotypes” for HTRX, HTR and GWAS, and visualized the distribution of the out-of-sample *R*^2^ for each of the best models selected by each method in Supplementary Information Figure S5.1. In both “lm” and “glm”, HTRX has equal predictive performance to the true model. It performs as well as GWAS when the interaction effects are absent, explains more variance when an interaction is present, and is significantly more explanatory than HTR. When rare SNPs are included, the only effective interaction term is rare. In this case the difference between GWAS and HTRX becomes smaller as expected, and removing the rare haplotypes hardly reduces the performance of HTRX.

In conclusion, we demonstrated through simulation that our HTRX implementation a) searches the haplotype space effectively and b) protects against overfitting. This makes it a superior approach compared to HTR and GWAS to integrate SNP effects with gene-gene interactions. Its robustness is also retained when there are rare effective SNPs and haplotypes.

### Quantifying selection via historical allele frequencies from Pathway Painting

The historical trajectory of SNP frequencies is a strong signal of selection when ancient DNA data are available. This is the main purpose of our Pathway Painting method, and can be used to infer selection at individual loci and combined into a polygenic score by analysing sets of SNPs associated with a trait.

Firstly, we inferred allele frequency trajectories and selection coefficients for a set of LD-pruned genome-wide significant trait associated variants using a modified version of CLUES (Coalescent Likelihood Under Effects of Selection) ^23^. To account for population structure in our samples, we applied a novel chromosome painting technique based on inference of a sample’s nearest neighbours in the marginal trees of an ARG that contains labelled individuals ^14^. We ran CLUES using a time-series of imputed aDNA genotype probabilities obtained from 1,015 ancient West Eurasian samples that passed all quality control filters. We produced four additional models for each trait associated variant by conditioning the analysis on one of the four ancestral path labels from our chromosome painting model: either Western Hunter-Gatherers (WHG), Eastern Hunter-Gatherers (EHG), Caucasus Hunter-Gatherers (CHG), or Anatolian farmers (ANA).

Secondly, we were able to infer polygenic selection gradients (ω) and p-values for each trait, i.e. of MS and RA, in all ancestral paths, using PALM (Polygenic Adaptation Likelihood Method) ^24^. Full methods and results can be found in Supplementary Note 6.

### Linkage Disequilibrium of Ancestry (LDA) and LDA Score (LDAS)

In population genetics, linkage disequilibrium (LD) is defined as the non-random association of alleles at different loci in a given population ^87^. Just like the values of the genotype, ancestries can be correlated along the genome, and further, deviations from the expected length distribution for a particular ancestry is a signal of selection, dated by the affected ancestry. We propose an ancestry linkage disequilibrium (LDA) approach to measure the association of ancestries between SNPs, and an LDA Score (LDAS) to quantify deviations from the null hypothesis that ancestry is inherited at random across loci.

LDA is defined in terms of local ancestry. Let *A*(*i, j, k*) denote the probability of the *k*th ancestry (*k* = 1,…, *K*) at the *j*th SNP (*j* = 1,…, *J*) of a chromosome for the *i*th individual (*i* = 1,…, *N*).

We define the distance between SNP *l* and *m* as the average *L*_2_ norm between ancestries at those SNPs. Specifically, we compute the *L*_2_ norm for the *i*th genome as

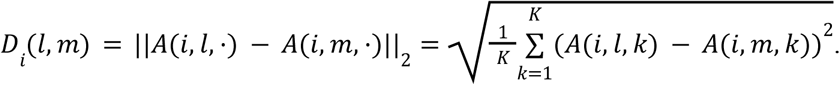

Then we compute the distance between SNP *l* and *m* by averaging *D*_*i*_ *(l, m)*

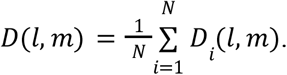

We define *D*^*^(*l, m*) as the theoretical distance between SNP *l* and *m* if there were no linkage disequilibrium of ancestry (LDA) between them. *D*^*^(*l, m*) is estimated by

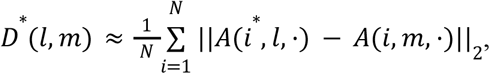

where *i*^*^ ∈ {1,…, *N*} are resampled without replacement at SNP *l*. Using the empirical distribution of ancestry probabilities accounts for variability in both the average ancestry and its distribution across SNPs. Ancestry assignment can be very precise in regions of the genome where our reference panel matches our data, and uncertain in others where we only have distant relatives of the underlying populations.

The LDA between SNP *l* and *m* is a similarity, defined in terms of the negative distance − *D*(*l, m*) normalized by the expected value *D*^*^(*l, m*) under no LD, as:

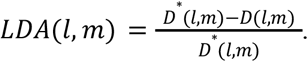

LDA therefore takes an expected value 0 when haplotypes are randomly assigned at different SNPs, and positive values when the ancestries of haplotypes are correlated.

LDA is a pairwise quantity. To arrive at a per-SNP property, we define the LDA score (LDAS) of SNP *j* as the total LDA of this SNP with the rest of the genome, i.e. the integral of the LDA for that SNP. Because this quantity decreases to zero as we move away from the target SNP, this is in practice computed within an *X*cM-window (we use *X* = 5 as LDA is approximately zero outside this region in our data) on both sides of the SNP. Note that we measure this quantity in terms of the genetic distance, and therefore LDAS is measuring the length of ancestry-specific haplotypes compared to individual-level recombination rates.

As a technical note, when the SNPs approach either end of the chromosome, they no longer have a complete window, which results in a smaller LDAS. This would be appropriate for measuring total ancestry correlations, but to make LDAS useful for detecting anomalous SNPs, we use the LDAS of the symmetric side of the SNP to estimate the LDAS within the non-existent window.

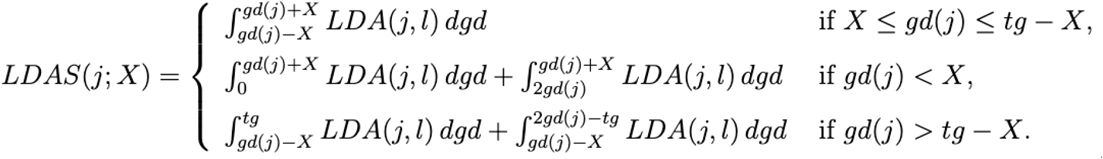

where *gd*(*l*) is the genetic distance (i.e. position in cM) of SNP *l*, and *tg* is the total genetic distance of a chromosome. We also assume the LDA on either end of the chromosome equals the LDA of the SNP closest to the end: *LDA*(*j, gd* = 0) = *LDA*(*j, l*_*min(gd)*_) and *LDA*(*j, gd* = *td*) = *LDA*(*j, l*_*max (gd)*_), where *gd* is the genetic distance, *l*_*min(gd)*_ and *l*_*max(gd)*_ are the indexes of the SNP with the smallest and largest genetic distance, respectively.

The integral 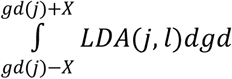 is computed assuming linear interpolation of the LDA score between adjacent SNPs.

LDA thus quantifies the correlations between the ancestry of two SNPs, measuring the proportion of individuals who have experienced a recombination leading to a change in ancestry, relative to the genome-wide baseline. The LDA score is the total amount of genome in LDA with each SNP (measured in recombination map distance).

### Simulation study for LDA and LDAS

An ancient population *P*_0_ evolved for 2200 generations before splitting into two sub-populations *P*_1_ (Steppe) and *P*_2_ (Farmer). After evolving for 400 generations, we added mutations “*m*_1_” and “*m*_2_” at different loci in *P*_1_ and *P*_2_. Both added mutations were then positively selected in the following 300 generations, after which they merged to *P*_3_, where both added mutations experienced strong positive selection for 20 generations. Finally, we sampled 1000 individuals from *P*_3_ to compute their ancestry proportions of *P*_1_ and *P*_2_ using the “chromosome painting” technique, and calculated the LDA score of the simulated chromosome positions.

The above describes the simulation in Supplementary Information Figure S7.1.

We investigated balancing selection at 2 loci as well. The balancing selection in *P*_1_ and *P*_2_ ensured the mutated allele reaches around 50% frequency, while positive selection made the mutated allele become almost the only allele. In *P*_3_, if *m*_1_ or *m*_2_ was positively selected, its frequency reached more than 80% regardless of whether the allele experienced balancing or positive selection in *P*_1_ or *P*_2_, because we set a strong positive selection. If *m*_1_ or *m*_2_ was balancing selected in *P*_3_, its frequency slightly increased, e.g. if *m*_1_ underwent balancing selection in *P*_1_, it had 25% frequency when *P*_3_ was created, and the frequency reached around 37.5% after 20 generations of balancing selection in *P*_3_.

The results (Supplementary Information Figure S7.2) show that positive selection in *P*_3_ resulted in low LDA scores around the selected locus, if this allele was not uncommon (i.e. had 50% (balancing selection) or 100% frequency (positive selection) in subpopulation *P*_1_ or *P*_2_). Note that the balancing selection in *P*_1_ or *P*_2_ worked the same as “weak positive selection”, because *m* and *m* were rare when they first occurred, which were positively selected until 50% frequency.

We also performed simulations for selection at a single locus (Supplementary Information Figure S7.2 and S7.3).

Stage 1: We added a mutation *m*_1_ in the 1600 generation in *P*_0_, which then underwent balancing selection until generation 2200, when *P*_0_ split into *P* _1_ and *P*_2_, where the frequency of *m* was around 50%.

Stage 2: Then we explored different combinations of positive, balancing and negative selection of *m*_1_ in *P*_1_ and *P*_2_. the frequency of *m*_1_ reached 80%, 50% and 20% when it was positively, balancing or negatively selected, respectively, until generation 2899. Here we sampled 20 individuals each in *P*_1_ and *P*_2_ as the ancient samples.

Stage 3: Then *P*_1_ and *P*_2_ merged into *P*_3_ in generation 2900. In *P*_3_, for each combination of selection in Stage 2, we simulated positive, balancing and negative selection for *m*_1_. The selection lasted for 20 generations, and then we sampled 4000 individuals from *P*_3_ as the modern population.

Results: when *m*_1_ was positively selected in only one of *P*_1_ and *P*_2_, and it experienced negative selection in *P*_3_, the LDA scores around the locus of *m*_1_ were low. Otherwise, no abnormal LDA scores were found at *m*_1_.

## Notes

### Competing Interest Statement

The authors have declared no competing interest.

### Summary of Updates

The article has been significantly re-written to improve clarity. The CLUES/PALM results have been updated with an improved model, but none of the headline results have changed.

