## Supplementary Information for "Elevated genetic risk for multiple sclerosis originated in Steppe Pastoralist populations"

#### Contents

**Contents** **1**
Key to Supplementary Data Tables 1
**1) Archaeological Sites and Data Generation** **2**
Archaeological sites 3
The churchyard of Our Lady/Vor Frue Kirkegård ÅHM6093 3
Ahlgade 15-17, MHO 71/85, Holbæk Parish, Holbæk, Sealand. 6
Tjærby, Gimming Parish, Randers, Jutland. KHM 0899. 14.09.04-20, GPS coordinates: 567982, 6259161. 9
Data Generation 10
Sampling, lab work and sequencing 10
Libraries authentication 11
References 12
**2) Analysis of mtDNA sequence, Sex determination and Y chromosome analysis** **13**
Analysis of mtDNA sequence 13
Methods 13
Results 13
Conclusion 17
Sex determination and Y chromosome analysis 19
Methods 19
Results 19
Sex determination 19
Phylogenetic placement 19
Conclusion 20
References 20
**3) Cluster Analysis, Weighted Average Prevalence and Local Ancestry GWAS** **22**
Methods 22
Cluster Analysis 22
Weighted Average Prevalence 23
PCA/UMAP of WAP/average dosage 24

GWAS 25
Local ancestry and genotype GWAS 25
Comparison of local ancestry and genotype gwas 26
Results 27
Weighted average prevalence 27
Association with MS risk at externally ascertained SNPs for ancestries and genotypes 27
Cluster analysis comparing between MS-risk and local ancestry for 3 example SNPs 27
In the HLA Class-II region, all SNPs share a pattern in which high Steppe ancestry is associated with high MS-risk. The risk decreases monotonically and is not present in the Steppe precursor populations (Hunter Gathers), but is with the admixed Bronze-age European populations (Steppe + Farmer). 27
PCA plots for average ancestry and weighted average prevalence of MS-associated SNPs 27
References 28
**4) Ancestral risk scores 28**
Introduction 28
Methods 29
Imputation of local ancestry 30
Ancestral risk score 30
Results 32
References 33
**5) Haplotype Trend Regression with eXtra flexibility (HTRX) 33**
Methods 33
Definition 33
HTRX Model selection procedure for shorter haplotypes 34
HTRX Model selection procedure for longer haplotypes (Cumulative HTRX) 36
Simulation for HTRX 37
Results 39
HTRX simulation 39
References 39
**6) Polygenic selection analysis for auto-immune disease risk 39**
Introduction 39
Methods 40
Sample data 40
GWAS ascertainment 40
Selection analysis 42
Polygenic selection analysis 43
Pleiotropic trait analysis 43
Joint polygenic selection analysis 44
Results 45
Multiple sclerosis 45
Pan-ancestry analysis 45
Western hunter-gatherer ancestral path 45
Eastern hunter-gatherer ancestral path 46
Caucasus hunter-gatherer ancestral path 46
Anatolian farmer ancestral path 47

Cross ancestry comparisons 48
Pleiotropic UK Biobank traits 49
Pleiotropic FinnGen traits 51
Joint polygenic selection analysis 55
Rheumatoid arthritis 56
Pan-ancestry analysis 57
Western hunter-gatherer ancestral path 58
Eastern hunter-gatherer ancestral path 60
Caucasus hunter-gatherer ancestral path 63
Anatolian farmer ancestral path 64
Cross ancestry comparisons 65
Pleiotropic UK Biobank traits 66
Pleiotropic FinnGen traits 68
Discussion 72
References 78
**7) Ancestry linkage disequilibrium (LDA) and Ancestry linkage disequilibrium score (LDAS) 80**
Methods 80
Definition 80
Simulation for selection: LDA 83
Results 85
Simulation for LDA scores with selection in one or two loci 85
LDA score for chromosome 2 and 6 85
LD plot for chromosome 6 MS-associated SNPs 85
References 85
**8) Pleiotropic effects of selected SNPs associated with multiple sclerosis or rheumatoid arthritis 86**
Introduction 86
Methods 89
Pleiotropic trait analysis 89
Results 92
Pleiotropic trait analysis of SNPs associated with multiple sclerosis 92
Pleiotropic trait analysis of SNPs associated with rheumatoid arthritis 95
Discussion 98
References 105

### Key to Supplementary Data Tables

| No. | Description |
|---|---|
| Supplementary Table 1 | Metadata for 87 new Danish Medieval and post-Medieval samples |
| Supplementary Table 2 | Data for each library analysed |
| Supplementary Table 3 | rsIDs used for fine-mapped variants in Patsopoulos 2019 ST11 |
| Supplementary Table 4 | ‘Fine-mapped’ and proxy SNPs used for MS, with painting effect allele frequencies |
| Supplementary Table 5 | Fine-mapped’ and proxy SNPs used for RA, with painting effect allele frequencies |
| Supplementary Table 6 | Fine-mapped’ and proxy SNPs used for CD, with painting effect allele frequencies |
| Supplementary Table 7 | CLUES results for LD-pruned SNPs for MS |
| Supplementary Table 8 | CLUES results for all SNPs associated with MS showing evidence of selection |
| Supplementary Table 9 | CLUES results for LD-pruned SNPs for RA |
| Supplementary Table 10 | CLUES results for all SNPs associated with RA showing evidence of selection |
| Supplementary Table 11 | Protective pleiotropic effects of MS-associated SNPs showing statistically significant evidence of selection |
| Supplementary Table 12 | Protective pleiotropic effects of RA-associated SNPs showing statistically significant evidence of selection |
| Supplementary Table 13 | Metadata for all ancient samples used in this study, including original publications |
| Supplementary Table 14 | Full results from Joint polygenic selection analysis |

### 1) Archaeological Sites and Data Generation

Gabriele Scorrano[1], Abigail Ramsøe[1], Charleen Gaunitz[1], Lasse Vinner[1], Thorfinn Sand Korneliussen[1], Fabrice Demeter[1], Marie Louise S. Jørkov[2], Stig Bermann Møller[3], Bente Springborg[3], Lutz Klassen[4], Inger Marie Hyldgård[4], Niels Wickmann[5]

[1]Lundbeck Foundation GeoGenetics Centre, GLOBE Institute, University of Copenhagen, Østervoldgade 5-7, DK-1350K, Copenhagen, Denmark

[2]Laboratory of Biological Anthropology, Department of Forensic Medicine, University of Copenhagen, Denmark

[3]Ålborg Historiske Museum, Nordjyske Museer, Vang Mark 25, 9380 Vestbjerg, Denmark

[4]Museum Østjylland. Stemannsgade 2, DK-8900 Randers C, Denmark

[5]Museum Vestsjælland, Forten 10, 4300 Holbæk, Denmark

#### Archaeological sites

##### The churchyard of Our Lady/Vor Frue Kirkegård ÅHM6093

Ålborg Sogn, Hornum Herred, Ålborg Amt Sted- og lokalitetsnr: 120516-107
Marie Louise Jørkov, Stig Bergmann Møller, Bente Springborg

The urban medieval churchyard of Our Lady (Vor Frue) and associated building structures were excavated by Aalborg Historiske Museum/Nordjyske Museer between 2011 to 2013. The churchyard belonged to the church and convent of Our Lady and is located in the eastern part of the medieval town of Aalborg. Approximately 900 graves were recovered of which 272 could be sampled for DNA analysis. The churchyard was excavated in connection with a large sewerage project, and only parts of the churchyard was exhumed.

The church of Our Lady was a parish church as well as a convent for nuns, probably of the Order of Saint Augustine, and date to the first half of the 12th century. A few radio-carbon dates open up a slight possibility that the churchyard, with an associated wooden church, can be dated back to the 11th century[1].

The convent was abandoned after the reformation in 1536. The area of the churchyard was used for burials from 1000/1050 to 1806. Burials were dated based on arm-positions (A to D), and all were represented[2]. However, since dating from arm-positions is somewhat uncertain, dating was also determined based on stratigraphy, dendrochronology, archaeological findings, as well as few radio-carbon dates[3,4].

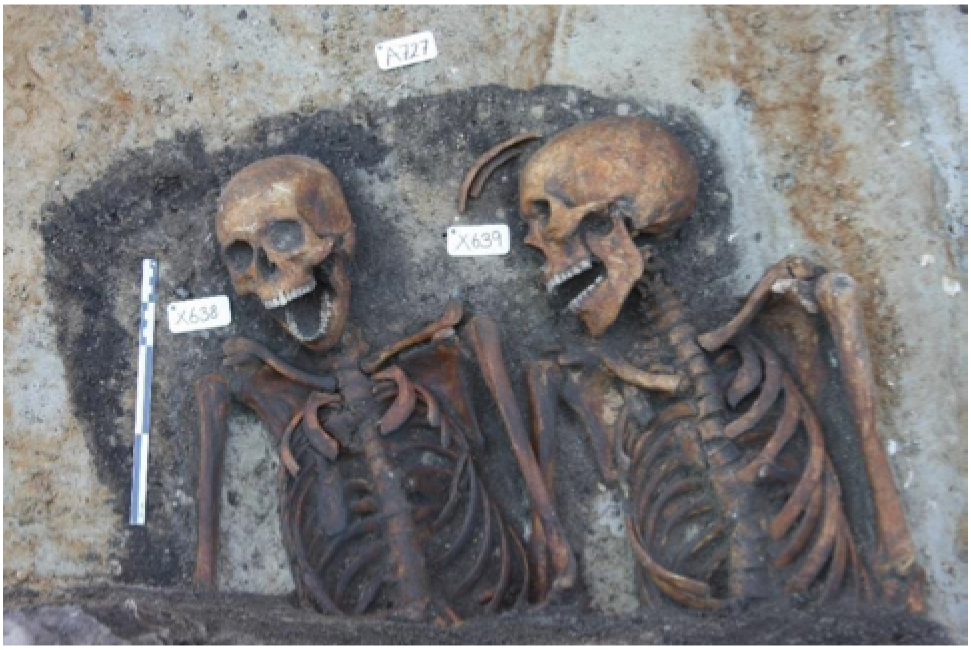


Photo from Springborg and Møller (2016): Two adults from grave A727 (x638: CGG_2_100494 and x639: CGG_2_100495) dated to ca. 1429-1496.

#### Ahlgade 15-17, MHO 71/85, Holbæk Parish, Holbæk, Sealand.

Marie Louise Jørkov and Niels Wickman

The cemetery Ahlgade 15-17 is an urban cemetery in the center of Holbæk, north of the main street of Ahlgade and adjacent to the fjord and the harbour. It was excavated in 1985 to 1986 by Museum Vestsjælland, previously called Museet for Holbæk og Omegn.

The cemetery belonged to the former parish church of St. Nicolai and date from the late 12th century to 1573 when the church was abandoned. However, the cemetery is thought to have been taken out of use shortly after the reformation in 1536.

Holbæk was a small medieval town that came into existence around 1200. It had two Parish churches: St. Nicolai and the contemporary Parish of Our Lady, located in the eastern part of town. There was also a castle, which according to written sources dates from the beginning of the 13th century[1]. The town was inhabited by families of craftsmen, farmers, fishermen and tradesmen.

The cemetery of St. Nicolai church is the first fully excavated urban cemetery in Denmark. It covered an area of ca. 480m2 where a minimum of 583 graves have been excavated[1,2]. The majority of the graves were dated based on arm positions of the skeletons (i.e. A-B-C and D) when lying in supine position and often with the head towards the east. These were identified in 309 graves but represents some degree of uncertainty[3] as there are considerable overlap between the use of arm positions and the time periods. The most commonly found arm position was position B, most likely corresponding to the time period 1300-1350. Selected graves have been radio carbon dated to confirm the dating range of the arm positions[1]. Only few burials could be dated from archaeological findings. The graves contained burials of adults and subadults and the distribution of adults were more or less equally divided in males and females in all periods. The majority of the subadult skeletons, however were found in the upper lays of the cemetery. This is most likely a reflection of poor preservation in the lower layers rather than specific burial practice.

Scallopshells from Santiago de Compostella were found in five of the graves 10 scallopshells in total. Two of these graves are included in this study (EG272 (CGG 2_101751) and EG620 (CGG 2_101617; 2_101588). The skeleton in grave EG272, an old adult individual of 50+ years, was buried with arm position B. The skeleton in grave EG620, a male aged 40-60 years, was found resting on the right side with bended legs and is also assumed to date to the 12th century[1].

A large dietary stable isotope study was carried out by Jørkov which showed that the population subsisted largely on a mixed diet of terrestrial animal plant protein as well as marine/brackish food sources[4,5].


#### Tjærby, Gimming Parish, Randers, Jutland. KHM 0899. 14.09.04-20, GPS coordinates: 567982, 6259161.


Marie Louise Jørkov, Lutz Klassen, Inger Marie Hyldgård


Rural cemetery ca. 5 km east of Randers on the north side of Randers fjord. It was excavated by Kulturhistorisk Museum Randers in 1998 to 2010. The excavation area revealed a stone church, and a cemetery containing ca. 1200 graves from which 351 individuals were sampled for DNA analysis in this project. The cemetery dates to ca. 1050 to late 1536, but skeletal remains were only preserved from graves dating after 1200. Remnants of a farmhouse and a wooden church predating the cemetery (900-1100) were also recovered[1,2]. The surrounding area consisted of forest and meadows.

The cemetry area measured ca. 45x42 meters (1900m2) and has been fully excavated. In nearly half of the graves, no skeletons were preserved. The remaining burials (673 in total) contained skeletons of varying degree of preservation from few poorly preserved fragments to well preserved and nearly complete skeletons due to the varying soil conditions in the area. The best preserved skeletons were found south of the church. Evidence of wooden coffins were found in some of the graves, but the majority of the individuals had been buried without. 57% of the graves with preserved bones contained adults (male and female), 10% subadults and 33% children.

The graves were dated based on the arm position of the skeletons[3] as well as from C-14 dating on selected individuals and from few archaeological finds such as coins and ceramics. More than half of the skeletons were found with arm position A (arms and hands down the side) representing period AD 1200-1300. One third had arm position B (arms along the side but hands on thigh or hips), which starts to appear in the 13th century, dominates in the 14th century and disappears around AD 1400. The remaining skeletons were buried with arm position C (elbows bended and lower arms on stomach) or D (arms bended and hands crossed over chest) representing the latest period AD 1400-1536. The majority of the graves were found south of the church. There was a significant decrease in burials during the period of use in particular in the latest period.

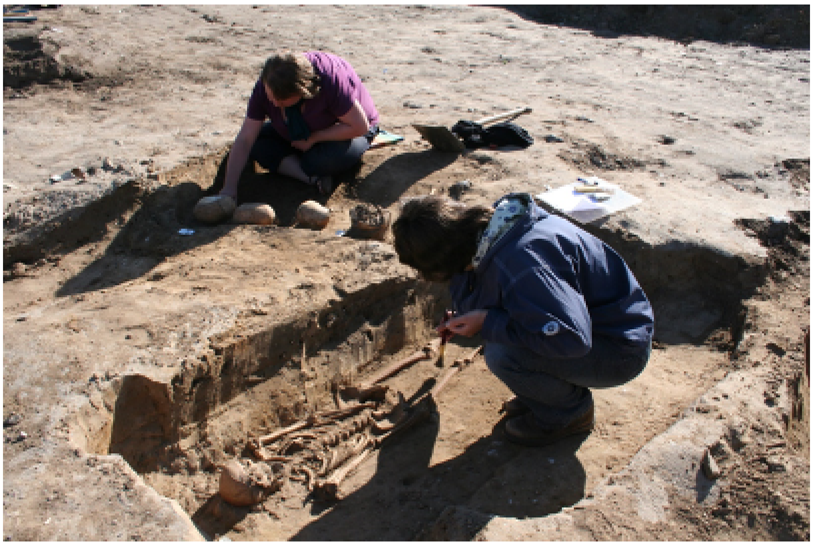

Excavation of Tjærby cemetery. Photo: Museum Østjylland.

### Data Generation

#### Sampling, lab work and sequencing

Charleen Gaunitz, Lasse Vinner

Sequencing data was generated from a total of 86 Medieval samples (ST1), using semi-automated laboratory procedures. Laboratory work on aDNA was conducted in the dedicated ancient DNA clean-room facilities at the Lundbeck Foundation GeoGenetics Centre (Globe Institute, University of Copenhagen).

In brief, two parallel sub-samples of <150 mg were obtained from human skeletal material and demineralized as described earlier, using pre-digestion for 30 min (Damgaard et al., 2015). Two aDNA extractions were performed per subsample, using a 96 well format, combining 150 µl of demineralized material with 1.5 ml binding buffer (500 ml Qiagen PB, supplemented with 15 ml Sodium acetate 3M, and 1.25 ml 5M NaCl, phenol red, adjusted to pH=5) and 10 µl of paramagnetic beads (G-Bioscience, #786-915) for 15 minutes (Rohland et al., 2018). Pelleted beads were washed twice in 450 µl and 100 µl 80% ethanol + 20% 10mM Tris-HCl, respectively, and eluted in 10 mM Tris-HCl + 0.05% Tween-20. From each subsample one extract (35 µl) was incubated with 10 µl USER enzyme (NEB #M5505) for 3h at 37°. DNA shotgun sequencing libraries were prepared in 96-well format essentially as described elsewhere (Meyer and Kircher 2010), using a small (25ul) or large (50ul) total reaction volume for non-USER and USER-treated extracts, respectively, including 21.25 µl or 42.5 µl DNA template. Clean-up procedures after end-repair and adapter-ligation were performed with 10 µl of paramagnetic beads (G-bioscience) in 10 volumes of the binding buffer described above. The requirement for PCR amplification was evaluated by qPCR using 1µl of pre-amplified library. Indexing PCR, using 8-bp unique dual indexing (Illumina TruSeq UDI0001-0096) in 50 or 100 µl reaction volumes, with KAPA HiFi HotStart Uracil+ (KapaBiosystems #KR0413) according to manufacturer's recommendations, with typically 14 amplification cycles. Final purification of libraries was performed using a 1:1.6 ratio of library to HighPrep™ PCR beads (MagBio, #AC-60250). Length distribution and concentration of individual purified libraries was controlled using the Fragment Analyzer (High Sensitivity kit). Libraries were pooled equimolar before sequencing. Sequencing was performed on Illumina NovaSeq6000 at the GeoGenetics Sequencing Core, Copenhagen, using S4 200 cycles kits version 1.5.

#### Libraries authentication

Gabriele Scorrano, Abigail Ramsøe, Thorfinn Sand Korneliussen

All libraries with at least 1X MT coverage for each sample were separately evaluated for contamination using ContamMix[1] before merging to sample level (Supplementary Table XX). ContamMix quantifies the fraction of exogenous reads in the mitochondrial reads by comparing the mtDNA consensus genome to possible contaminant genomes.

For each library, an in-house perl script was used to construct the endogenous mitochondrial genome. The parameters to do this depended on the coverage of each library: for libraries below 5X (but above 1X), sites were called if it was at least 2X coverage, whereas for libraries above 2X this value was set to a 5X cutoff. For both methods, bases were only called if 70% of reads at a site agreed with the consensus call.

We further investigated libraries showing over 5% contamination (<95% map authentic value). Two libraries showed over 5% contamination with a very low MT depth coverage (1.96X and 2.40X, Supplementary Table XX) however low coverage, below 3 X, can return unreliable results[1]. One library with coverage ~ 14 X had a map authenticity close to the edge (94,5%). However, merging these libraries with the other ones from the same samples did not result in a meaningful level of contamination (Supplementary Table X). The slight levels of contamination reported by ContamMix are thus most probably due to errors while constructing the consensus genome caused by a combination of very low depth and the characteristic C->T damage of non-USER treated ancient samples. Then all the libraries have been merged and included in the analysis.


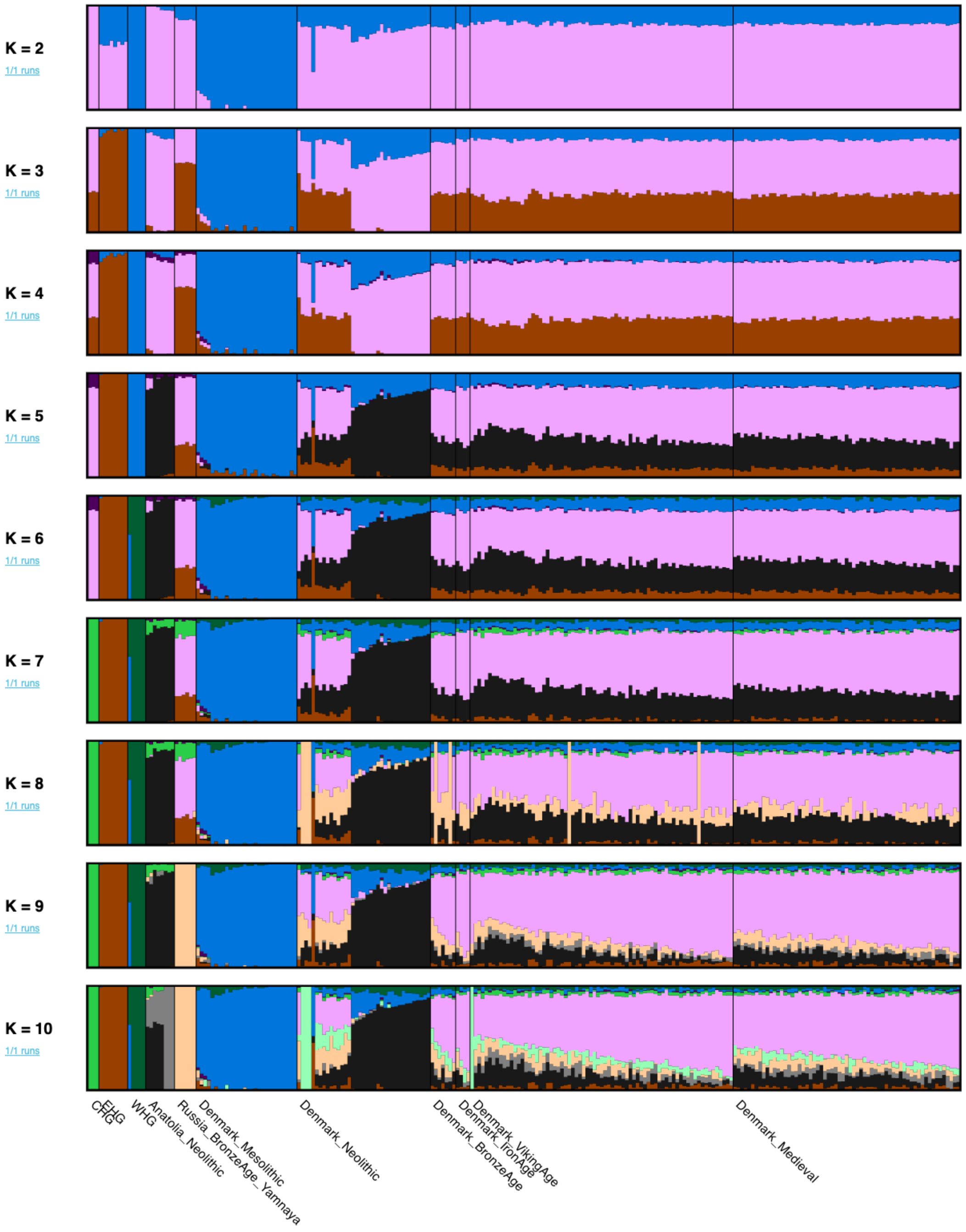


**Figure S1.1**: ADMIXTURE results performed with Caucasina hunter-gatherers (CHG), Eastern hunter-gatherers (EHG), Western hunter-gatherers (WHG), Anatolian Neolithic and Russia Bronze Age Yamnaya (Steppe ancestry) as the source populations from k=2 to k=10.

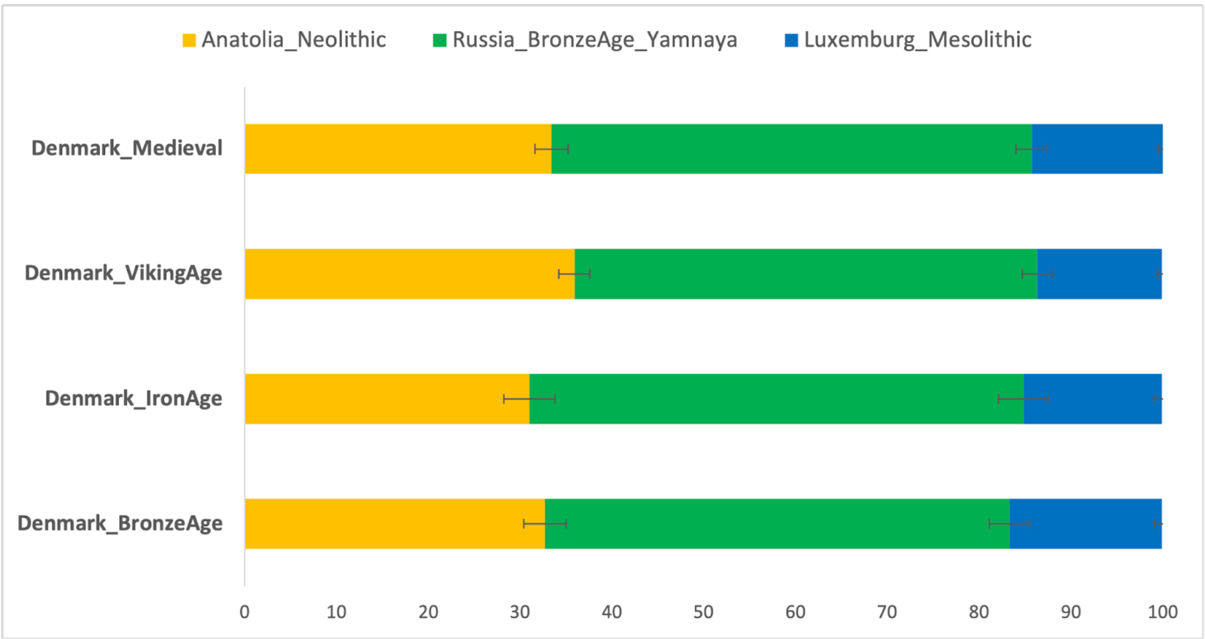


**Figure S1.2:** Three-way *qpAdm* model with the source populations Luxemburg_Mesolithic (WHG), Russia_Bronze_Age_Yamnaya (Steppe ancestry) and Anatolia Neolithic. Error bars represent standard errors of the proportion of each component. The complete results are reported in Table SXq1.

**Table S1.1:** Results obtained by *qpAdm* analysis for the three-way admixture model.

| | **Component** | | | **Standard Deviation** | | | **nSNPs** | **p-value** |
|---|---|---|---|---|---|---|---|---|
| | Anatolia_Neolithic | Russia_BronzeAge_Yamnaya | Luxemburg_Mesolithic | Anatolia_Neolithic | Russia_BronzeAge_Yamnaya | Luxemburg_Mesolithic | 838230 | |
| Denmark_Medieval | 33.4 | 52.3 | 14.3 | 1.8 | 1.7 | 0.5 | 838230 | 0.07 |
| Denmak_Viking Age | 35.9 | 50.4 | 13.6 | 1.7 | 1.6 | 0.5 | 838230 | 0.30 |
| Denmar | 31 | 53.8 | 15.1 | 2.8 | 2.7 | 0.9 | 838230 | 0.61 |

| | | | | | | | | |
|---|---|---|---|---|---|---|---|---|
| k_IronAge | | | | | | | | |
| Denmark_BronzeAge | 32.7 | 50.6 | 16.6 | 2.3 | 2.2 | 0.8 | 838230 | 0.53 |

### 2) Analysis of mtDNA sequence, Sex determination and Y chromosome analysis

Gabriele Scorrano[1]

[1]Lundbeck Foundation GeoGenetics Centre, GLOBE Institute, University of Copenhagen, Østervoldgade 5-7, DK-1350K, Copenhagen, Denmark

#### Analysis of mtDNA sequence

Gabriele Scorrano

##### Methods

We carried out a phylogenetic tree analysis using the reconstructed mitochondrial genomes from the individuals presented in this study. We reconstructed the mitochondrial consensus sequence from genome-wide bam files by ANGSD v.0.928 filtering out reads with a mapping quality lower than 20. Moreover, we limited the minimum depth of coverage to 5x and used -doFasta 2, -doCounts 1 and -trim 5 (which trims the first and last five nucleotides from each read) options (Korneliussen et al. 2014). All fasta files have finally been aligned using mafft and following phylotree recommendations (van Oven and Kayser, 2009). The phylogenetic tree analysis was performed by BEASTv1.10.4 (Drummond et al. 2012) and BEAUti v1.10.4 was used to generate the input file for the analysis. After having analysed the aligned fasta file with jModelTest 2.1.10 (Darriba et al., 2012), we applied the TrN (Tamura and Nei, 1993) substitution model with gamma plus invariant sites and strict clock with a prior of 2.74 $10^{-8}$ µ/site/year (Posth et al., 2016). We used the Bayesian Skyline coalescent with 10 as group members and piecewise-linear as the Skyline model (Posth et al., 2016). We performed two MCMC runs with $50^{8}$ states and sampled every $10^{5}$ states and evaluated the runs by Tracer (v1.6) (http://tree.bio.ed.ac.uk/software/tracer/). The two independent runs were combined using LogCombiner v1.8.1 and summarized in a Maximum Clade Credibility (MCC) tree using TreeAnnotator v1.8.1 (both programs included in the BEAST package). The resulting trees were annotated by TreeAnnotator v1.8.4 and visualized by FigTreev1.4.3 (http://tree.bio.ed.ac.uk/software/figtree/).

##### Results

The phylogenetic tree gives an overview of the distribution of mitochondrial haplogroups in our dataset (Figure S2.1). The frequency of haplogroups is similar to that found in the modern population (Bybjerg-Grauholm et al., 2018), H is the haplogroup with highest frequency (33.7%).

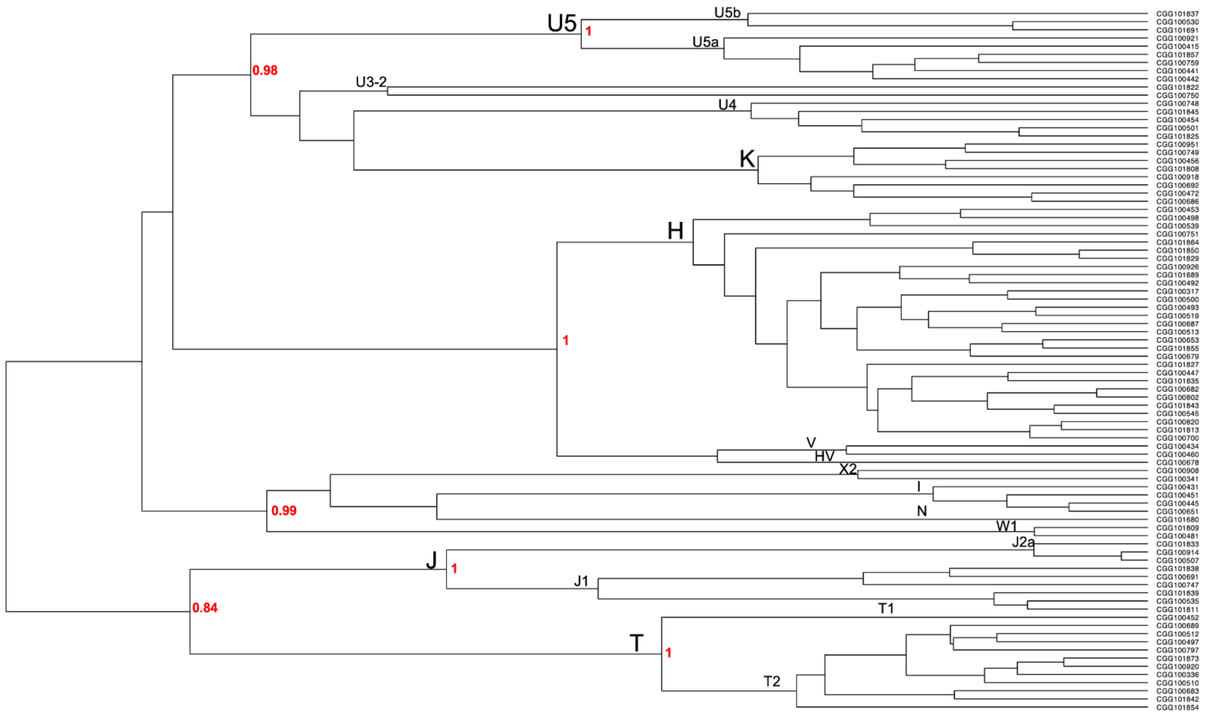


**Figure S2.1: Mitochondrial phylogenetic tree**: Phylogenetic tree of mitochondrial haplogroups found in the 86 Danish Medieval and post-Medieval samples. Estimated bootstrap values for principal nodes are reported in red.

Among the most common haplogroups identified we find haplogroup U, of which the subclade U5 which is commonly observed among Western European Hunter-Gatherers.

The most common subclades of U5 are U5a and U5b. Most of the Medieval and post-Medieval individuals belong to the sub-haplogroups U5a1. U5a1 has eastern origin, and has been found among Mesolithic individuals from Russia, Ukraine and also from Sweden (Lazaridis et al., 2014). It is also found in high frequencies among the Yamnaya population (Allentoft et al., 2016), suggesting that the spread of this sub-haplogroup in Europe could be associated with the spread of the Steppe ancestry from Eastern Europe. U5a1 shows high frequency in Denmark during the Viking age (Margyaran et al., 2020). This result is congruent with the autosomal structure of Danish populations through time (Figure S1.1 and Figure S1.2) where a genetic continuity from the Viking age to the post-Medieval period has been highlighted.

The first individual with U5b comes from 30 kya, U5b is in fact one of the most ancient mitochondrial European haplogroups and it spreads around 20 kya (Fu et al. 2016; Scorrano et al., 2021) but today it appears with low frequencies among the modern populations (De Angelis et al. 2018; Scorrano et al. 2021a).

The genetic influence of the spread of Early Farmers into Europe is shown by the great frequencies of the haplogroups K (9.3%) and J (10.5%). In particular the sub-clades K1a and K1b are mainly found in Neolithic populations across Europe (Antonio et al., 2019; Brunel et al., 2020; Gamba et al., 2014; Hofmanova et al., 2016). Moreover, they had great frequencies across the Scandinavian Viking age people (Margyaran et al., 2020).

#### Conclusion

In conclusion we find most of the haplogroups still present in the modern-day Danes suggesting that there have been no major migrations in Denmark since the Medieval period. The sub-clades identified are in accordance with the autosomal results showing a great frequency of the mitochondrial haplogroups associated with the spread of Steppe ancestry, but at the same time also a mitochondrial Neolithic contribution. Moreover, the mitochondrial results show a mitochondrial genetic continuity from the Viking age to the post-Medieval period.

### Sex determination and Y chromosome analysis

Gabriele Scorrano

#### Methods

Genetic sex for each individual was determined following the method suggested by Skoglund et al. (2013). For the male individuals Y chromosome haplogroup assignment was inferred and a clustering tree has been performed following an already published workflow (Scorrano et al 2021). EPA-ng (Barbera et al., 2019) has been used to place the ancient Y-chromosome sequences in the reference tree.

#### Results

##### Sex determination

We unambiguously determined genetic sex for all 86 individuals (37 female, 49 males; ST1). In Figure S2.2 the individuals form two clearly separated clusters corresponding to male (XY) and female (XX) using the normalised sequencing coverage across the sex chromosomes: X and Y.

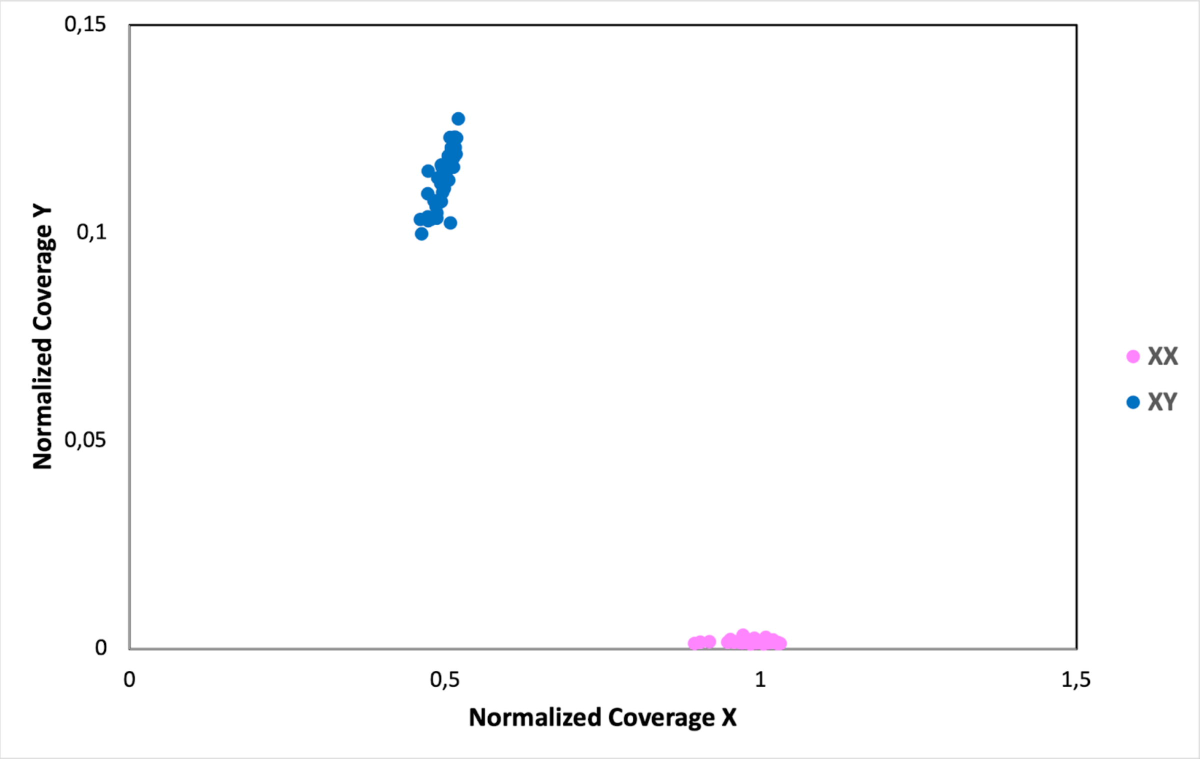


**Figure S2.2: Sex determination:** plot showing the normalised X and Y coverage by autosomal coverage for each individual reported in this study. Symbol colors indicate the inferred sex chromosome karyotype.

#### Phylogenetic placement

We built a clustering tree to analyse the Y-chromosome diversity across the 49 Medieval and post-Medieval male individuals from Denmark (Figure S2.3) and different haplogroups and sub-haplogroups have been identified.

One individual belongs to the sub-haplogroup E1b1b, identified for the first time in Mesolithic Israeli samples of the Natufian culture, and exists today at a low frequency in northern Europe (Lazaridis et al., 2016). One sample belongs to the sub-haplogroups N1a1a, a subclade that has higher average frequency in northern Europe (Derenko et al., 2007; Lappalainen et al., 2008). However, most of the samples fall within two main haplogroups: I and R.

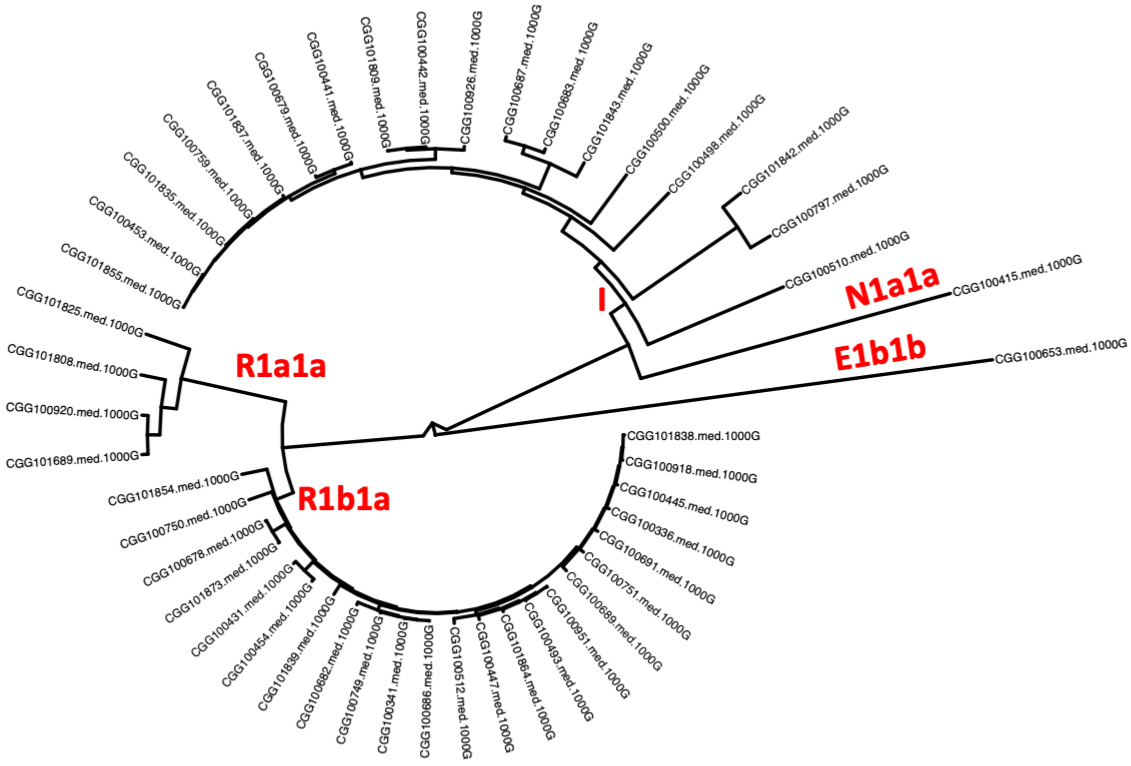


**Figure S2.3: Y chromosome phylogenetic tree**: Phylogenetic tree with most likely placements of ancient samples here presented.

Haplogroup I is the oldest major haplogroup in Europe, haplogroup IJ is thought to have arrived from the Middle East to Europe around 35,000 years ago, developing later into haplogroup I.

It was speculated that I1 lineage was among the Mesolithic European hunter-gatherers (Szécsényi-Nagy et al. 2015) and spread in Scandinavia from northern Germany. The sub-haplogroup I1a is the most common Y-chromosome I haplogroup in Northern Europe and it was already identified in Danish individuals from the Viking age (Margyaran et al., 2020).

The sub-haplogroup I2 originated during the Late Paleolithic, the oldest I2 sample was recovered from Bichon (13,500 year-old), a Late Palaeolithic man from Switzerland (Jones et al., 2015). Loschbour from Mesolithic Luxembourg is the oldest I2a1b individual (Lazaridis et al., 2014), while in Scandinavia the first evidence of I2a1b is with Motala (7,800 year-old) (Mittnik et al., 2018). However, most of the individuals belong to the sub-lineages of the haplogroup R1: R1a and R1b. The sub-lineages R1a-M417 (R1a1a1) is the most prevalent male lineage in the extant eastern and northern European people, and it was associated with the spread of Steppe ancestry in Europe (Allentoft et al.,

2015; Haak et al., 2015). The R1a identified lineages have been present in Denmark since the Viking Age (Margyaran et al., 2020).

Appreciable frequency of male sub-lineage R1b was identified in northern Europe starting from the Mesolithic (Jones et al., 2017), through the Neolithic (Allentoft et al., 2015) and the Viking age (Margyaran et al., 2020) until the Medieval. Today it is the most common Y-chromosome haplogroup in Western Eurasia.

#### Conclusion

In conclusion male haplogroups found with high frequencies are already identified during the Viking age, suggesting a Y-chromosome genetic continuity from the Viking Age to the post-Medieval period. Moreover, the haplogroups observed with high frequency are still the most abundant male lineage of Western Europe.

### 3) Cluster Analysis, Weighted Average Prevalence and Local Ancestry GWAS

Yaoling Yang[1,2], Daniel Lawson[1,2]

[1]Department of Statistical Sciences, School of Mathematics, University of Bristol, UK
[2]Integrative Epidemiology Unit, Population Health Sciences, University of Bristol, UK

#### Methods

##### Cluster Analysis

In order to understand whether risk-conferring haplotypes evolved in the Steppe population, or in a pre- or post-dating population in which Steppe ancestry is high, we used k-means clustering on the dosage of each ancestry for each selected significant SNP and investigated the dosage distribution of clusters with significantly higher MS prevalence. For the target SNPs, the Elbow method (Thorndike 1953). suggested selecting around 5-7 clusters, of which we chose 6. After performing the k-means cluster analysis, we calculated the average probability for each ancestry for case individuals. Furthermore, we calculated the prevalence of MS in each cluster, and performed a one-sample t-test to investigate whether it differs from the overall MS prevalence (0.487%). This tests whether particular combinations of ancestry are associated with the phenotype at a SNP. Clusters with high MS risk-ratios have high Steppe components (Figure S3.3), leading to the conclusion that Steppe ancestry alone is driving this signal.

##### Weighted Average Prevalence

In order to quantify the risk of each ancestry for each SNP, we calculated the weighted average prevalence (WAP) for each ancestry based on the result of k-means clustering (above).

For the $j$th SNP, let $P_{jkm} = n_{jm}\overline{P}_{jkm}$ denote the sum of the $k$th ancestry probabilities of all the individuals in the $m$th cluster ($k, m = 1,..., 6$), where $n_{jm}$ is the cluster size of the $m$th cluster. Let $\pi_{jm}$ denote the prevalence of MS in the $m$th cluster, the weighted average prevalence for the $k$th ancestry is defined as:

$$\overline{\pi}_{jk} = \frac{P_{jkm}\pi_{jm}}{\sum_{m=1}^{6} P_{jkm}},$$

where $P_{jkm}$ is defined as the weight for each cluster.

The standard deviation of $\bar{\pi}_{jk}$ is computed as $sd(\bar{\pi}_{jk}) = \sqrt{\sum_{m=1}^{6} w_{jkm}^2 \sigma_m^2}$, where $w_{jkm} = \frac{P_{jkm}}{\sum_{m=1}^{6} P_{jkm}}$,

$\sigma_m = \frac{s(y_{jm})}{\sqrt{n_{jm}}}$ and $s(y_{jm})$ is the standard deviation of the outcome for the individuals in the $m$th cluster. We also test the hypothesis that $H_0$: $\bar{\pi}_{jk} = \bar{\pi}$ against $H_1$: $\bar{\pi}_{jk} \neq \bar{\pi}$, and compute the p-value as $p_{jk} = 2(1 - \phi(\frac{|\bar{\pi} - \bar{\pi}_{jk}|}{sd(\bar{\pi}_{jk})}))$.

For each ancestry, WAP measures the association of that ancestry with MS risk across all clusters. To make a clear comparison, we calculated the risk ratio (compared to the overall MS prevalence) for each ancestry at each SNP, and assigned a mean and confidence interval for the risk ratios of each ancestry at each chromosome (Figure 3, Figure S3.1).

#### PCA/UMAP of WAP/average dosage

To sort risk-associated SNPs into ancestry patterns according to that risk, we performed PCA on the average ancestry probability and WAP at each MS-associated SNP (Figure S3.4). The former shows that all of the HLA SNPs except three from HLA class II and III have much larger Outgroup components compared with the others. The latter analysis indicates a strong association between Steppe and MS risk. Also, Outgroup ancestry at rs10914539 from chromosome 1 exceptionally reduces the incidence of MS, while Outgroup ancestry at rs771767 (chromosome 3) and rs137956 (chromosome 22) significantly boosts MS risk.

#### GWAS

##### Local ancestry and genotype GWAS

We used the UK Biobank to fit GWAS models for local ancestry values and genotype values separately, using only SNPs known to be associated with the phenotype (‘fine-mapped’ SNPs). We used the following phenotype codes for each phenotype: MS: Data-Field 131043; RA: Data-Field 131849 (seropositive).

Let $Y_i$ denote the phenotype status for the $i$th individual ($i = 1,..., 399998$), which takes value 1 for a case and 0 for control, and let $\pi_i = Pr(Y_i = 1)$ denote the probability that this individual has the event. Let $X_{ijk}$ denote the $k$th ancestry probability ($k = 1,..., K$) for the $j$th SNP ($j = 1,..., 205$) of the

$i$th individual. $C_{ic}$ is the $c$th predictor ($c = 1, ..., N_c$) for the $i$th individual. We used the following logistic regression model for GWAS, which assumes the effects of alleles are additive:

$$Y_i \sim Bin(1, \pi_i);\ log(\frac{\pi_i}{1-\pi_i}) = \sum_{k=1}^{K} \beta_{jk} X_{ijk} + \sum_{c=1}^{N_c} \gamma_c C_{ic}.$$

We used $N_c$=20 predictors in the GWAS models, including sex, age and the first 18 PCs, which are sufficient to capture most of the population structure in the UK Biobank (Sarmanova et al., 2020[1]).

First, we built the model with $K = 1$. By using only one ancestry probability in each model, we aimed to find the statistical significance of each SNP under each ancestry. Then, we built the model with $K = 5$, i.e. using all 6 local ancestry probabilities which sum to 1. We calculated the variance explained by each SNP by summing up the variance explained by $X_{ijk}$(k=1,...,5).

We considered fitting the multivariate models by using all the SNPs as covariates. However, the dataset only contains 1,982 cases. Even though only one ancestry is included, the multivariate model contains 191 predictors, which could result in overfitting problems. Therefore, the GWAS models are preferred over multivariate models.

We also fitted a logistic regression model for GWAS using the genotype data as follows:

$$Y_i \sim Bin(1, \pi_i);\ log(\frac{\pi_i}{1-\pi_i}) = \beta_j X_{ij} + \sum_{c=1}^{N_c} \gamma_c C_{ic},$$

where $X_{ij} \in \{0, 1, 2\}$ denotes the number of copies of the reference allele of the jth SNP ( $j = 1, ..., 205$) that the $i$th individual has, and $C_{ic}$ ($c = 1, ..., N_c$) denotes the covariates including age, sex and first 18 PCs for the ith individual, where $N_c$=20. Due to the UK Biobank being underpowered compared to the Case-Control study from which these SNPs were found, the only statistically significant (at $p < 10^{-5}$) association is in the HLA class II tagging HLA-DRB1*15:01.

#### Comparison of local ancestry and genotype gwas

We compared the variance explained by SNPs from the GWAS model using the painting data (all 6 local ancestry probabilities) with that from GWAS model using the genotype data. McFadden's pseudo R squared measure (McFadden et al., 1973[2]) is widely used for estimating the variance explained by the logistic regression models. McFadden's pseudo R squared is defined as

$$R^2 = 1 - \frac{ln(L_M)}{lm(L_0)},$$

where $L_M$ and 0 are the likelihoods for the fitted and the null model, respectively. Taking overfitting into account, we propose the adjusted McFadden's pseudo R squared by penalizing the number of predictors:

$$Adjusted\,R^2 = 1 - \frac{ln(L_M)/(N-k)}{ln(L_0)/(N-1)},$$

where N is the sample size and k is the number of predictors.

Specifically, $R^2(SNPs)$ is calculated as the extra variance in addition to sex, age and 18 PCs that can be explained by SNPs:

$$R^2(SNPs) = R^2(sex + age + 18\,PCs + SNPs) - R^2(sex + age + 18\,PCs).$$

Notably, two SNPs stand out for explaining much larger variance than others when fitting the GWAS model using the genotype data, but overall more SNPs from GWAS painting explain more than 0.1% variance, which indicates the painting data are probably more efficient for estimating the effect sizes of SNPs and detecting significant SNPs. Also, some SNPs from GWAS models using painting data explain almost the same amount of variance, suggesting that these SNPs consist of very similar ancestries.

### Results

#### Weighted average prevalence

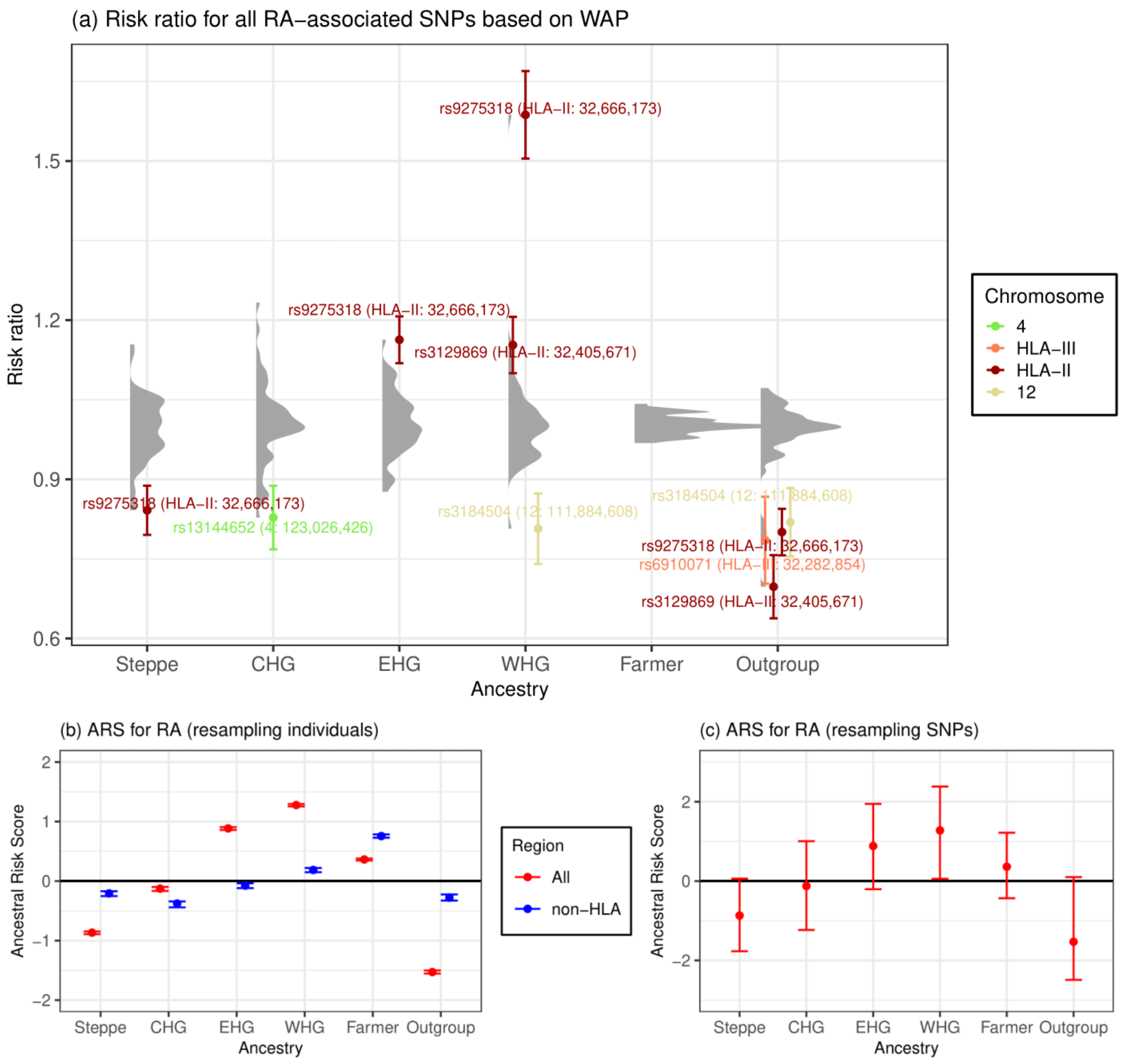


**Figure S3.1. Associations between local ancestry at fine-mapped RA SNPs and RA in a modern population.**

a) Risk ratio of SNPs for RA based on weighted average prevalence (WAP; see Methods), when decomposed by inferred ancestry. A mean and standard deviation are calculated for each ancestry based on bootstrap resampling, for each chromosome. The distribution of risk ratios at each ancestry is shown as a raincloud plot. SNPs significant at the 1% level are shown individually, coloured by chromosome or HLA region, and those with risk ratio >1.1 or <0.9 are annotated with rsID, HLA region and position (build GRCh37/hg19). b-c) Genome-wide Ancestral Risk Scores (ARS, see Methods) for RA. Confidence intervals are estimated by either bootstrapping over individuals (b, which can be interpreted as testing power to reject a null hypothesis of no association between RA and ancestry) and bootstrapping over SNPs (c, which can be interpreted as testing whether ancestry is associated with RA genome-wide). We show results for all associated SNPs (red) and non-HLA SNPs only (blue) when bootstrapping over individuals.

#### Association with MS risk at externally ascertained SNPs for ancestries and genotypes

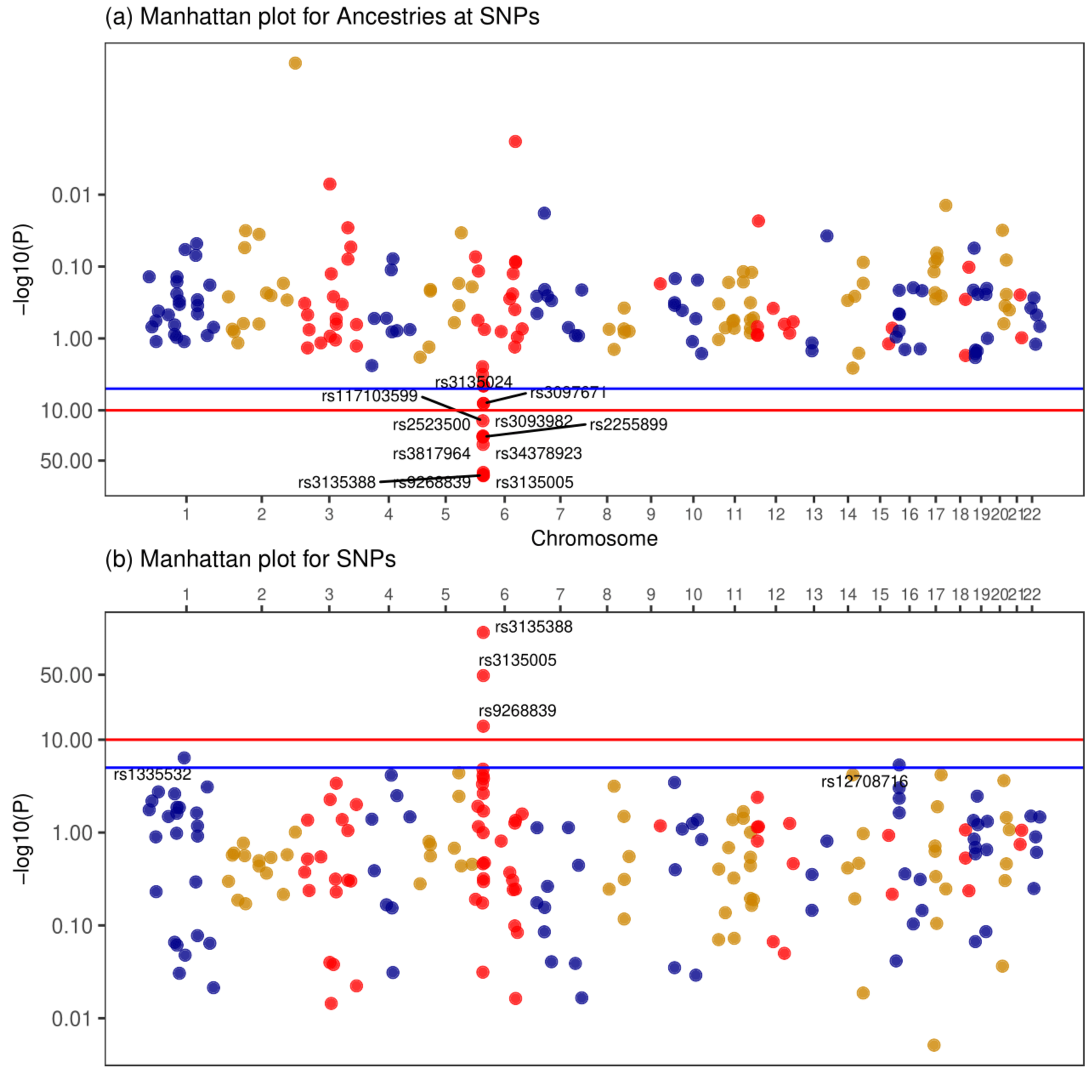


**Figure S3.2: Association with MS risk at externally ascertained SNPs, for (top) ancestry, and (bottom) SNPs**.

Due to the UK Biobank being less powered (having fewer cases) than the Case-Control study from which these SNPs were found, the only statistically significant association is in the HLA.

#### Cluster analysis comparing between MS-risk and local ancestry for 3 example SNPs

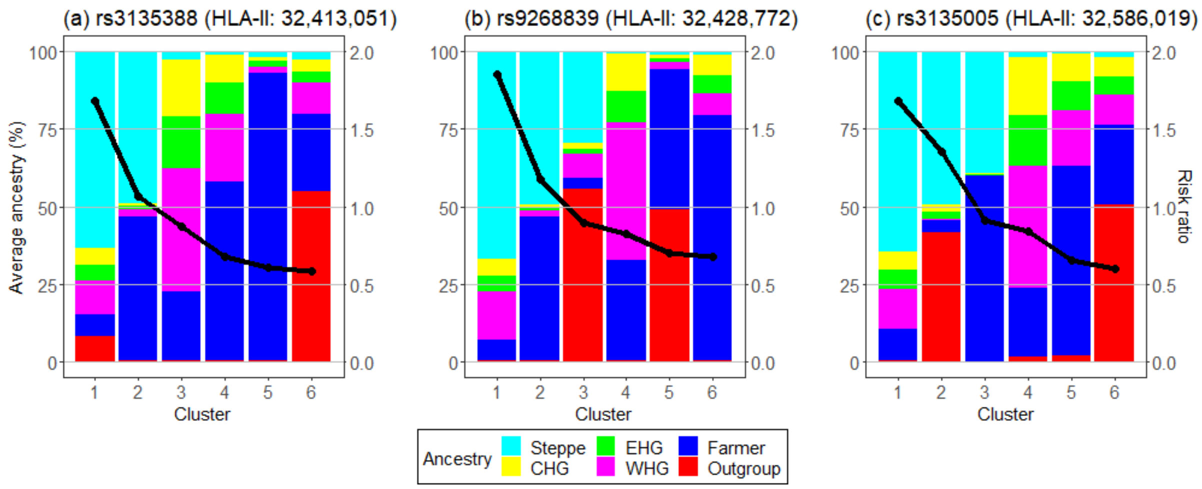


**Figure S3.3: Comparison between MS-risk and local ancestry for 3 example SNPs**.

In the HLA Class-II region, all SNPs share a pattern in which high Steppe ancestry is associated with high MS-risk. The risk decreases monotonically and is not present in the Steppe precursor populations (Hunter Gathers), but is with the admixed Bronze-age European populations (Steppe + Farmer).

### PCA plots for average ancestry and weighted average prevalence of MS-associated SNPs

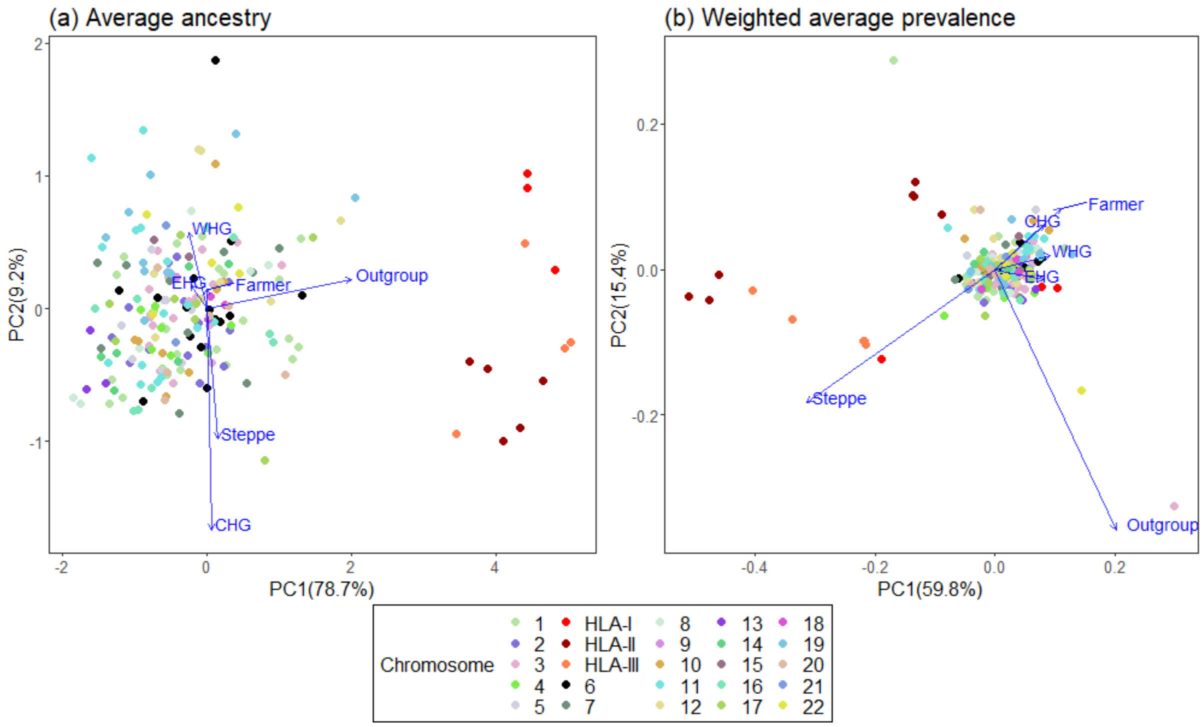


**Figure S3.4: Decomposition of individuals' ancestry at MS risk SNPs in terms of (left) the ancestry of those SNPs alone, or (right) the Weighted average prevalence of MS in each ancestry after "logit" transformation.**

### 4) Ancestral risk scores

William F.S. Barrie[1], Daniel Lawson[2,3]

[1]Zoology Department, University of Cambridge, UK

[2]Department of Statistical Sciences, School of Mathematics, University of Bristol, UK

[3]Integrative Epidemiology Unit, Population Health Sciences, University of Bristol, UK

#### Introduction

Because panels of ancient individuals are small and geographically biased, allele frequency estimates based directly on aDNA genotype calls have low confidence (Dehasque et al., 2020). Equally, selection or drift (e.g. from population bottlenecks) mean that the allele frequency in an ancient population does not necessarily reflect the proportion of effect alleles that that ancestry eventually contributed to a modern population. Therefore, a better estimate of an ancestral contribution is to generate allele frequencies based on local ancestry: if a haplotype is under-sampled in the ancient data or undergoes subsequent positive selection, this will be reflected in an allele frequency that is higher in the estimate based on the painting than one based on the ancient data. We refer to these frequencies as "painting frequencies".

This approach was used to estimate ancestral contributions to a range of phenotypes in Irving-Pease et al. (2022), re-capitulating already known contributions such as height, hair colour and eye colour.

#### Methods

All code for implementing these analyses can be found at https://github.com/will-camb/ms_paper.

##### Imputation of local ancestry

Because not all SNPs in the GWAS data were painted, for each variant in each GWAS dataset we imputed the local ancestry by taking the average of the painting values of the SNPs on either side, weighted by their physical distance (impute_ancestry.py).

##### Ancestral risk score

Following methods developed in Irving-Pease et al. (2022), we calculated the effect allele painting frequency for a given ancestry $f_{\{anc,i\}}$ for SNP $i$ using the formula:

$$f_{\{anc,i\}} = \frac{\sum_{j}^{M_{effect}} Painting\ certainty_{\{j,i,anc\}}}{\sum_{j}^{M_{alt}} Painting\ certainty_{\{j,i,\ anc\}} + \sum_{j}^{M_{effect}} Painting\ certainty_{\{j,i,\ anc\}}},$$

where there are $M_{effect}$ individuals homozygous for the effect allele, $M_{alt}$ individuals homozygous for the other allele, and $\sum_{j}^{M_{effect}} Painting\ certainty_{\{j,i,anc\}}$ is the sum of the painting probabilities for that ancestry $anc$ in individuals homozygous for the effect allele at SNP $i$. This can be interpreted as an estimate of an ancestral contribution to effect allele frequency in a modern population. Per-SNP painting frequencies can be found in ST4, ST5, and ST6.

To calculate the ancestral risk score (ARS) we summed over all $I$ pruned SNPs in an additive model:

$$ARS_{anc} = \sum_{i}^{I} f_{\{anc,i\}} * \ beta_i.$$

We then ran a transformation step as in Berg & Coop (2014). To obtain 95% confidence intervals, we ran an accelerated bootstrap over loci, which accounts for the skew of data to better estimate confidence intervals (Frangos & Schucany, 1990).

### Results

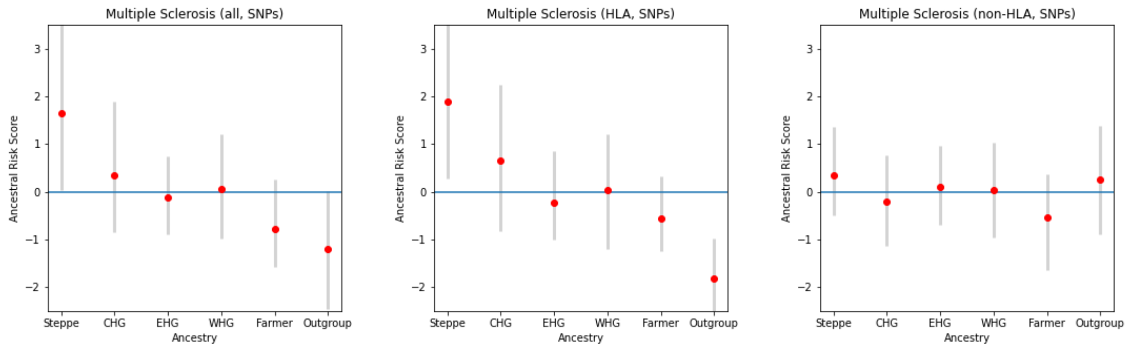


**Figure S4.1.** Ancestral Risk Scores (ARS) for fine-mapped data for MS using all SNPs (left), only SNPs on the HLA(centre), and only SNPs not on the HLA (right). Confidence intervals are estimated by bootstrapping over SNPs, which can be interpreted as testing whether ancestry is associated with MS genome-wide.

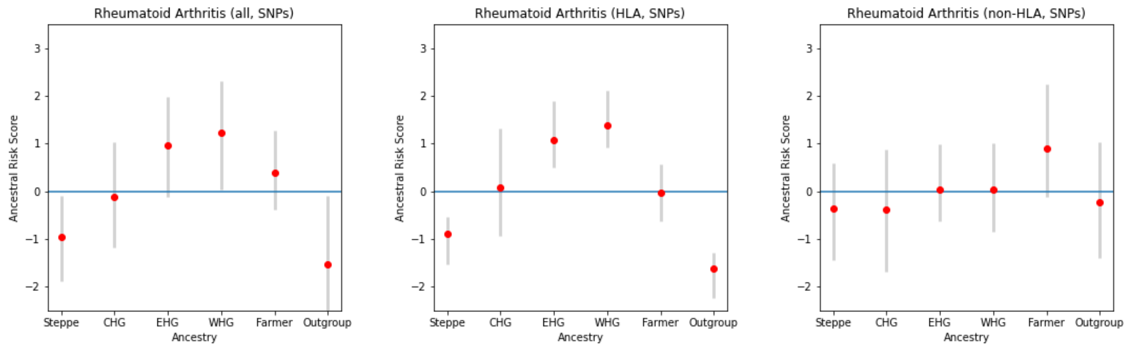


**Figure S4.2.** Ancestral Risk Scores (ARS) for fine-mapped data for RA using all SNPs (left), only SNPs on the HLA(centre), and only SNPs not on the HLA (right). Confidence intervals are estimated by bootstrapping over SNPs, which can be interpreted as testing whether ancestry is associated with RA genome-wide.

### 5) Haplotype Trend Regression with eXtra flexibility (HTRX)

Yaoling Yang[1,2], Daniel Lawson[1,2]

[1]Department of Statistical Sciences, School of Mathematics, University of Bristol, UK
[2]Integrative Epidemiology Unit, Population Health Sciences, University of Bristol, UK

#### Methods

##### Definition

We propose Haplotype Trend Regression with eXtra flexibility (HTRX) which searches for haplotype patterns that include single SNPs and non-contiguous haplotypes. HTRX is an association between a template of $n$ SNPs and a phenotype. A template gives a value for each SNP taking values of '0' or '1', reflecting whether the reference allele of each SNP is present or absent, or an 'X' meaning either value is allowed. For example, haplotype '1X0' corresponds to a 3-SNP haplotype where the first SNP is the alternative allele and the third SNP is the reference allele, while the second SNP can be either the reference or the alternative allele. Therefore, haplotype '1X0' is essentially only a 2-SNP haplotype.

To examine the association between a haplotype and a binary phenotype, we replace the genotype term with a haplotype from the standard GWAS model:

$$Y_i \sim Bin(1, \pi_i);\ log(\frac{\pi_i}{1-\pi_i}) = \beta_j H_{ij} + \sum_{c=1}^{N_c} \gamma_c C_{ic},$$

where $H_{ij}$ denotes the $j$th haplotype probability for the $i$th individual:

$$H_{ij} = \begin{cases} 1 & \text{if } i\text{th individual has haplotype } j \text{ in both genomes,} \\ 1/2 & \text{if } i\text{th individual has haplotype } j \text{ in one of the two genomes,} \\ 0 & \text{otherwise.} \end{cases}$$

HTRX can identify gene-gene interactions, and is superior to HTR not only because it can extract combinations of significant SNPs within a region, leading to improved predictive performance, but the haplotypes are more interpretable as multi-SNP haplotypes are only reported when they lead to increased predictive performance.

#### HTRX Model selection procedure for shorter haplotypes

Fitting HTRX models directly on the whole dataset can lead to significant overfitting, especially when the number of SNPs increases. When overfitting occurs, the models experience poorer predictive accuracy against unseen data. Further, HTRX introduces an enormous model space which much be searched.

To address these problems, we implement a two-step procedure.

Step 1: select **candidate** models. This is to address the model search problem, and is chosen to obtain a set of models more diverse than traditional bootstrap resampling (Efron, 1979[1])

(1) Randomly sample a subset (50%) of data. Specifically, when the outcome is binary, stratified sampling is used to ensure the subset has approximately the same proportion of cases and controls as the whole data;

(2) Start from a model with fixed covariates (18 PCs, sex and age), and perform forward regression on the subset, i.e. iteratively choose a feature (in addition to the fixed covariates) to add whose inclusion enables the model to explain the largest variance, and select $s$ models with the lowest Bayesian Information Criteria (BIC) (Kass 1995) to enter the candidate model pool;

(3) repeat (1)-(2) $B$ times, and select all the different models in the candidate model pool as the candidate models.

Step 2: select the best model using 10-fold cross-validation.

(1) Randomly split the whole data into 10 groups with approximately equal sizes, using stratified sampling when the outcome is binary;

(2) In each of the 10 folds, use a different group as the test dataset, and take the remaining groups as the training dataset. Then, fit all the candidate models on the training dataset, and use these fitted models to compute the additional variance explained by features (out-of-sample $R^2$) in the test dataset. Finally, select the candidate model with the biggest average out-of-sample $R^2$ as the best model.

#### HTRX Model selection procedure for longer haplotypes (Cumulative HTRX)

Longer haplotypes are important for discovering interactions. However, there are $3^k - 1$ haplotypes in HTRX if the region contains $k$ SNPs, making it unrealistic for regions with large numbers of SNPs. To address this issue, we proposed cumulative HTRX to control the number of haplotypes, which is also a two-step procedure.

Step 1: extend haplotypes and select candidate models.

(1) Randomly sample a subset (50%) of data, use stratified sampling when the outcome is binary. This subset is used for all the analysis in (2) and (3);

(2) Start with $L$ randomly chosen SNPs from the entire $k$ SNPs, and keep the top $M$ haplotypes that are chosen from the forward regression. Then add another SNP to the $M$ haplotypes to create $3M + 2$ haplotypes. There are $3M$ haplotypes obtained by adding `0', `1' or `X' to the previous $M$ haplotypes, as well as 2 bases of the added SNP, i.e. `XX...X0' and `XX...X1' (as `X' was implicitly used in the previous step). The top $M$ haplotypes from them are then selected using forward regression. Repeat this process until obtaining $M$ haplotypes which include $k - 1$ SNPs;

(3) Add the last SNP to create $3M + 2$ haplotypes. Afterwards, start from a model with fixed covariates (18 PCs, sex and age), perform forward regression on the training set, and select $s$ models with the lowest BIC to enter the candidate model pool;

(4) repeat (1)-(3) $B$ times, and select all the different models in the candidate model pool as the candidate models.

Step 2: select the best model using 10-fold cross-validation, as described in "**HTRX Model selection procedure for shorter haplotypes**".

We note that because the search procedure in Step 1(2) may miss some highly predictive haplotypes, cumulative HTRX acts as a lower bound on the variance explainable by HTRX.

As a model criticism, only common and highly predictive haplotypes (i.e. those with the greatest adjusted $R^2$) are correctly identified, but the increased complexity of the search space of HTRX leads to haplotype subsets that are not significant on their own but are significant when interacting with other haplotype subsets being missed. This issue would be eased if we increase all the parameters $s$, $l$,

$M$ and $B$ but with higher computational cost, or improve the search by optimizing the order of adding SNPs. This leads to a decreased certainty that the exact haplotypes proposed are `correct', but together reinforces the inference that interaction is extremely important.

#### Simulation for HTRX

To investigate how the total variance explained by HTRX compare to GWAS and HTR, we used a simulation study comparing:

(1) linear models (denoted by "lm") and generalized linear models with a logit link-function (denoted by "glm");

(2) models with or without actual interaction effects;

(3) models with or without rare SNPs (frequency smaller than 5%);

(4) remove or retain rare haplotypes when rare SNPs exist.

We started from creating the genotypes for 4 different SNPs $G_{ijq}$ ($i = 1,..., 100,000$ denotes the index of individuals, $j = 1("1XXX"),\ 2("X1XX"),\ 3("XX1X")\ and\ 4("XXX1")$ represents the index of SNPs, and $q = 1, 2$ for two genomes as individuals are diploid). If no rare SNPs were included, we sampled the frequency $F_j$ of these 4 SNPs from 5% to 95%; otherwise, we sampled the frequency of the first 2 SNPs from 2% to 5% (in practice, we obtained $F_1 = 2.8\%$ and $F_2 = 3.1\%$ under our seed) while the last 2 SNPs from 5% to 95%. For the $i$th individual, we sampled $G_{ijq} \sim Bin(1, F_j)$ for the $q$th genome of the $j$th SNP, and took the average value of two genomes as the genotype for the $j$th SNP of the $i$th individual: $G_{ij} = \frac{G_{ij1}+G_{ij2}}{2}$. Based on the genotype data, we obtained the haplotype data for each individual, and we considered removing haplotypes rarer than 0.1% or not when rare SNPs were generated. In addition, we sampled 20 fixed covariates (including sex, age and 18 PCs) $C_{ic}$ where $c = 1,..., 20$ from UK Biobank for 100000 individuals.

Next, we sampled the effect sizes of SNPs $\beta_{G_j}$ and covariates $\beta_{C_c}$, and standardize them by their standard deviations: $\beta_{G_i} \sim \frac{U(-1,1)}{sd(G_j)}$ and $\beta_{C_c} \sim \frac{U(-1,1)}{sd(C_c)}$ for each fixed $j$ and $c$, respectively. When interaction exists, we created a fixed effect size for haplotype "11XX" as twice the average absolute SNP effects: $\beta_{H_1} = \frac{1}{2}\sum_{j=1}^{4} |\beta_{G_j}|$ where $H_1$ refers to "11XX"; otherwise, $H_1 = 0$. Note that $F_{H_1} = 0.09\%$ when rare SNPs are included.

Finally, we sampled the outcome based on the outcome score (for the $i$th individual)

$$O_i = \sum_{c=1}^{20} \beta_c C_{ic} + \gamma(\sum_{j=1}^{4} \beta_{G_j} G_{ij} + \beta_{H_1} H_1) + e_i + w,$$

where $\gamma$ is the effect scale of SNPs and haplotype "11XX", $e_i \sim N(0, 0.1)$ is the random error and $w$ is a fixed intercept term. For linear models, the outcome $Y_i = O_i$; while for generalized linear models, we sampled the outcome from binomial distribution: $Y_i \sim Bin(1, \pi_i)$, where $\pi_i = \frac{e^{O_i}}{1+e^{O_i}}$ is the probability that the $i$ th individual has the case.

As the simulation is intended to compare the variance explained by HTRX, HTR and SNPs (GWAS) in addition to fixed covariates, we tripled the effect sizes of SNPs and haplotype "11XX" (if interaction exists) by setting $\gamma = 3$. In "glm", to ensure a reasonable case prevalence (e.g. below 5%), we set $w =- 7$, which was also applied in "lm".

We applied the procedure described in " **HTRX Model selection procedure for shorter haplotypes**" for HTRX, HTR and GWAS, and visualized the distribution of the out-of-sample $R^2$ for each of the best models selected by each method in Figure S5.1. In both "lm" and "glm", HTRX has equal predictive performance as the true model. It performs as well as GWAS when the interaction effects is absent, explains more variance when an interaction is present, and is significantly more explanatory than HTR. When rare SNPs are included, the only effective interaction term is rare. In this case the difference between GWAS and HTRX becomes smaller as expected, and removing the rare haplotypes hardly reduces the performance of HTRX.

In conclusion, we demonstrate through simulation that our HTRX implementation a) searches haplotype space effectively, and b) protects against overfitting. This makes it a superior approach compared to HTR and GWAS to integration SNP effects with gene-gene interaction. Its robustness also retains when there are rare effective SNPs and haplotypes.

### Results

#### HTRX simulation

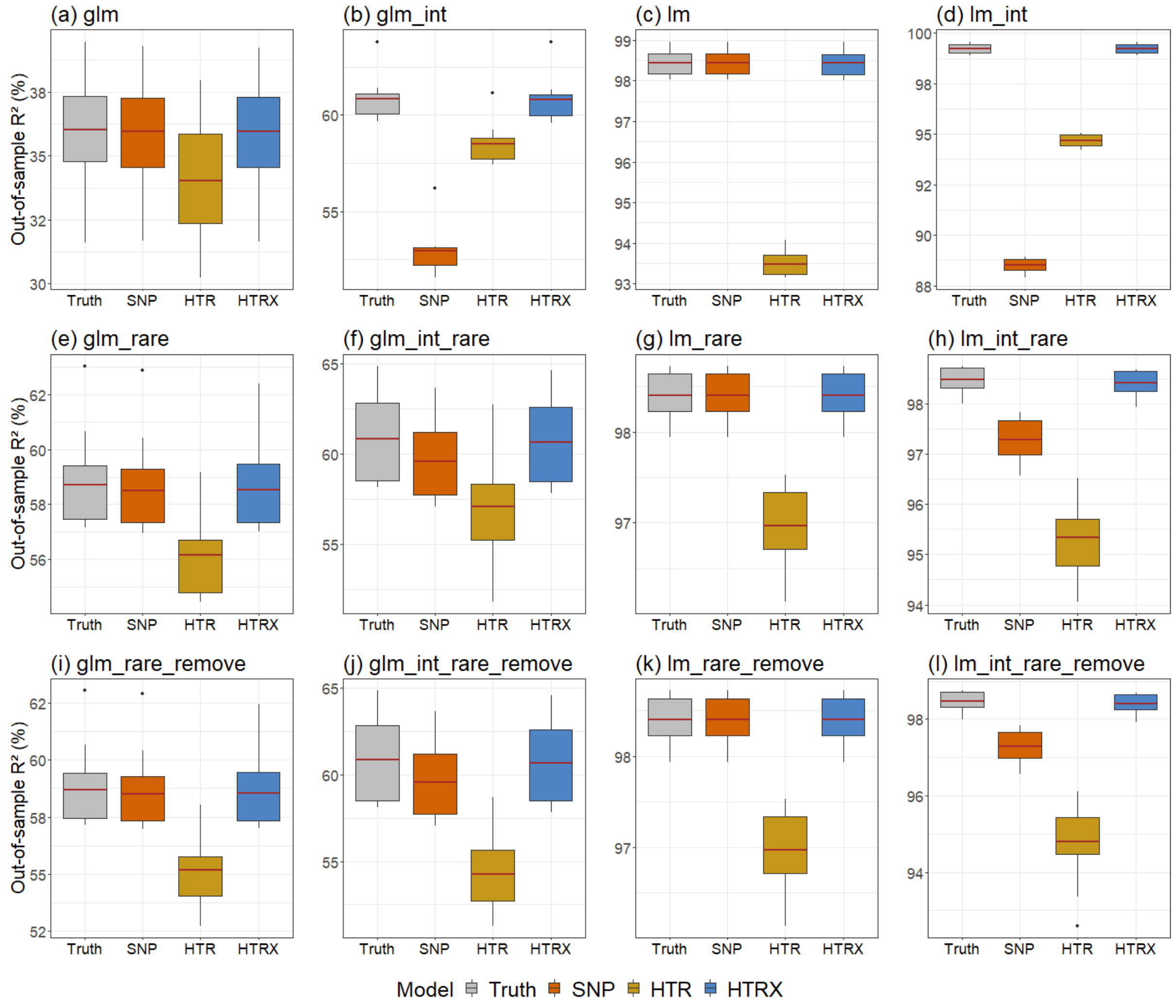


**Figure S5.1: Simulation study with four SNPs showing the boxplots of out-of-sample variance (with the red line representing the average) explained by HTRX compared to GWAS, HTR and the true model.**

The total variance explained by HTRX is the same as SNP and bigger than HTR when there are no interactions. When interaction (with subtitle "int") exists, HTRX significantly outperforms GWAS and HTR. In all the situations, HTRX works similarly as the truth.

### 6) Polygenic selection analysis for auto-immune disease risk

Evan K. Irving-Pease[1], Aaron J. Stern[2], Rasmus Nielsen[1,2]

[1] Lundbeck Foundation GeoGenetics Centre, GLOBE Institute, University of Copenhagen, Copenhagen, Denmark

[2] Department of Integrative Biology, University of California, Berkeley

#### Introduction

The rates and prevalence of auto-immune diseases in present-day populations varies considerably across West Eurasia. The drivers of this variation are complex, and include both environmental and genetic components. Understanding the genetic component of auto-immune disease is important for predicting patient risk and for developing novel therapeutics. Auto-immune disease susceptibility, like other complex traits, has been subject to evolutionary processes that have shaped the patterns of genetic variation seen in present-day populations. To understand how natural selection has influenced the genetic component of auto-immune disease risk, we sought to model the allele frequency trajectories of risk associated variants through time, in a large panel of ancient genomes, and to test for evidence of polygenic selection acting on two auto-immune diseases: (i) multiple sclerosis (MS); and (ii) rheumatoid arthritis (RA).

#### Methods

The computational pipeline to perform these analyses was written in the snakemake workflow management system[1]. For a full list of all the software and versions used, see Table S6.1. All pipeline code, custom scripts and a conda environment to replicate the analyses are available in the GitHub repository (https://github.com/ekirving/ms_paper).

##### Sample data

Our analyses are undertaken on a large sample of shotgun-sequenced ancient genomes, presented in the accompanying study 'Population Genomics of Stone Age Eurasia'[2]. This dataset comprises 1,664 imputed diploid ancient genomes and more than 8.5 million SNPs. Here, we used a subset of 1,015 imputed ancient genomes from West Eurasia, which passed all quality control filters. To account for population structure in our samples, we applied a novel chromosome painting technique based on inference of a sample's nearest neighbours in the marginal trees of an ARG that contains labelled individuals. Details of the chromosome painting methods are presented in the accompanying study 'The Selection Landscape and Genetic Legacy of Ancient Eurasians'[3] and further described in reference [4].

#### GWAS ascertainment

We obtained genome-wide association study (GWAS) summary statistics for two auto-immune diseases: (i) multiple sclerosis[5]; and (ii) rheumatoid arthritis[6–8]. To ascertain statistically independent and genome-wide significant markers, we filtered the GWAS summary statistics to retain only biallelic SNPs that passed quality control filters in our imputed dataset, then performed LD-based clumping using the software PLINK[9], with the parameters `--clump-p1 5e-8 --clump-r2 0.05 --clump-kb 250`, using the 1000 Genomes Project (1000G) Phase 3 populations FIN, GBR and TSI[10] as the combined LD reference panel. We also obtained fine-mapped SNPs for each of the two traits, and inferred proxy SNPs in high LD with variants which were not present in our imputed ancient callset (see methods).

#### Selection analysis

We inferred allele frequency trajectories and selection coefficients for all genome-wide significant trait associated variants using a modified version of the software CLUES[11], following the methods described in our companion paper[3]. In brief, we jointly inferred genome-wide genealogies and a population size history for the 1000G populations FIN, GBR and TSI using the software Relate. We then used CLUES to infer allele frequency trajectories and selection coefficients using a time-series of imputed ancient DNA (aDNA) genotype probabilities, and using the population size history from Relate. We produced four additional models for each trait associated variant, by conditioning the analysis on one of the four ancestral path labels from our chromosome painting model: either Western hunter-gatherers (WHG), Eastern hunter-gatherers (EHG), Caucasus hunter-gatherers (CHG), or Anatolian farmers (ANA). We determined statistical significance of each CLUES model by applying a Bonferroni correction for the number of tests performed for each trait.

#### Polygenic selection analysis

We inferred polygenic selection gradients ($\omega$) and p-values for MS and RA using the software PALM[12]. We used the modified version of CLUES to extract the posterior likelihood surface for each statistically independent marker, using the `--lik` argument, and restructured the outputs into the PALM folder structure. We then transformed the GWAS summary statistics into PALM format (w/ beta and SE scores) and standardised allele polarisation (ALT as the effect allele), then ran the polygenic selection analysis in single trait mode.

#### Pleiotropic trait analysis

To explore the extent to which selected variants share pleiotropic associations with other traits, we obtained GWAS summary statistics for 4,359 traits from the UK Biobank (UKB)[13,14] and 2,202 traits from the FinnGen study[15]. We filtered the summary statistics to retain only genome-wide significant trait associations (p < 5e-8) for SNPs which were found to be statistically significant (using a Bonferroni corrected significance threshold) in our CLUES analysis for MS and RA. We then produced plots for each UK Biobank trait with more than one significant SNP, showing the trajectories of pleiotropic SNPs polarised by their effect direction in the marginal trait.

#### Joint polygenic selection analysis

To determine if the observed signal of polygenic selection in MS could be better explained by selection acting on a genetically correlated trait, we performed a systematic analysis of traits in UKB and FinnGen. To identify traits with a shared genetic architecture to MS, we took the list of SNPs with significant CLUES p-values from the MS analysis and queried both UKB and FinnGen to retrieve a list of all traits in which the selected MS SNPs also had a genome-wide association with another trait. We then used a cut off of 20% overlap to narrow down the list of 6,561 possible traits (i.e. for a trait to be considered, at least 20% of the MS selected SNPs must also be associated with that trait). This resulted in 115 traits with a shared architecture to MS, including 49 in UKB and 66 in FinnGen.

To ensure that all SNPs used in the joint analysis were callable in both the focal GWAS (i.e. MS) and the marginal GWAS (i.e. UKB or FinnGen) we took the intersection of the SNPs available in both GWAS and in our imputed ancient callset. We then performed SNP clumping in the marginal trait, following the same procedure as we used for MS, to ascertain a list of independent markers. We then ran the J-PALM joint analysis, comparing MS to each of the 115 marginal traits, and repeated this for each of the 5 ancestry paths. Full results are available in table ST14.

**Table S6.1.** Software and versions used in the polygenic selection pipeline

| Software | Version | URL | Reference |
|---|---|---|---|
| bcftools | 1.15 | https://github.com/samtools/bcftools | [16] |
| biopython | 1.79 | https://github.com/biopython/biopython | [17] |
| clues | d8ec8b4 | https://github.com/standard-aaron/clues | [11] |
| conda | 4.14.0 | https://github.com/conda/conda | [18] |
| numpy | 1.22.2 | https://github.com/numpy/numpy | [19] |
| palm | 1d3b7fc | https://github.com/standard-aaron/palm | [12] |
| pandas | 1.3.4 | https://github.com/pandas-dev/pandas | [20] |
| plink | 1.90b6.21 | https://www.cog-genomics.org/plink/ | [9] |
| pysam | 0.18.0 | https://github.com/pysam-developers/pysam | [21] |
| python | 3.9.7 | https://www.python.org | [22] |
| r-base | 4.0.5 | https://www.r-project.org/ | [23] |
| r-dplyr | 0.8.0.1 | https://github.com/tidyverse/dplyr | [24] |
| r-ggplot2 | 3.1.1 | https://github.com/tidyverse/ggplot2 | [25] |
| r-ggrepel | 0.9.1 | https://github.com/slowkow/ggrepel | [26] |
| r-ggridges | 0.5.1 | https://github.com/wilkelab/ggridges | [27] |
| r-stringr | 1.4.0 | https://github.com/tidyverse/stringr | [28] |
| relate | 1.1.3 | https://myersgroup.github.io/relate | [29] |
| scipy | 1.8.0 | https://github.com/scipy/scipy | [30] |
| snakemake | 6.12.3 | https://github.com/snakemake/snakemake | [1] |

### Results

#### Multiple sclerosis

The CLUES results for all genome-wide significant MS associations are available in ST8, and the results for the subset of statistically independent markers used in the PALM analysis are available in ST7.

##### Pan-ancestry analysis

The PALM results for the pan-ancestry analysis of MS, using 62 LD-pruned markers, found statistically significant evidence for directional polygenic selection (p = 1.02e-5; ω = 0.017).

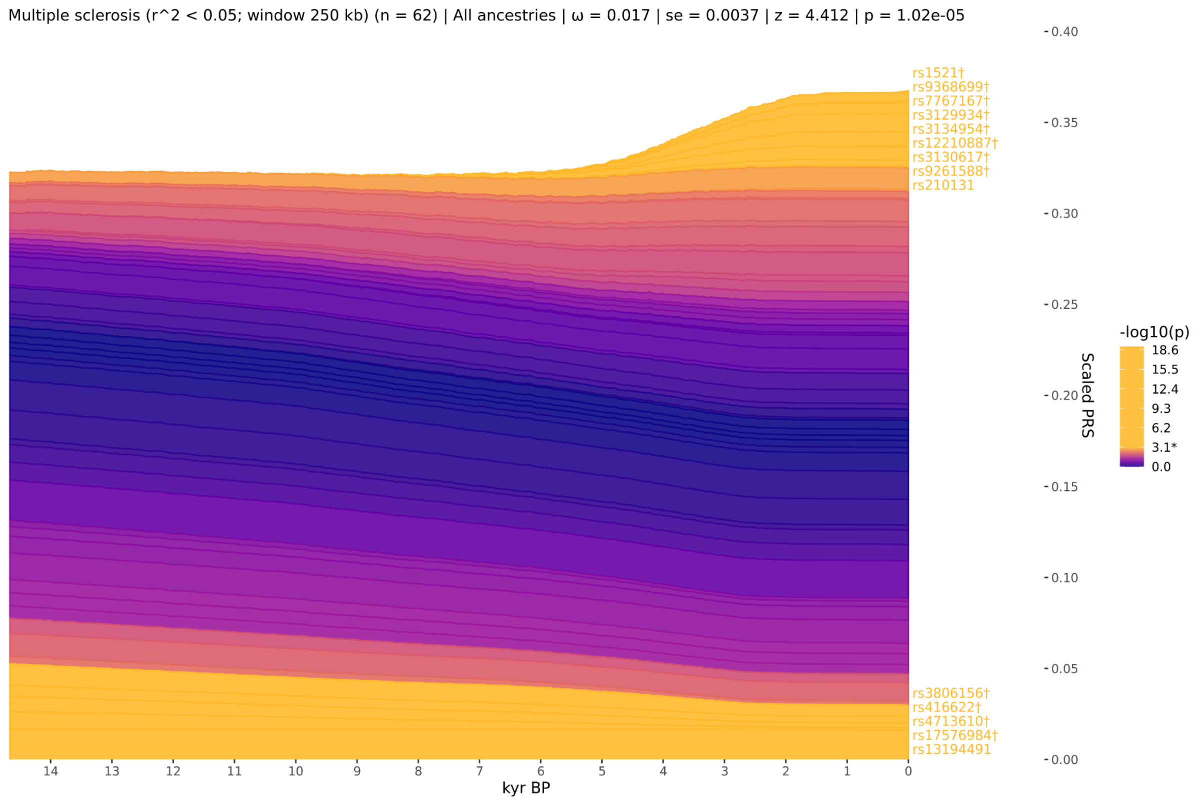


**Figure S6.1.** Stacked line plot of the pan-ancestry *PALM* analysis for Multiple sclerosis, showing the *CLUES* inferred allele frequency trajectories of each statistically independent SNP. Individual trajectories have been polarised to show the frequency of the positive risk allele, weighted by their scaled effect size. The y-axis shows the scaled polygenic risk score (PRS), which ranges from 0 to 1, representing the maximum possible additive genetic risk in a population. SNP trajectories are sorted by their *CLUES* p-values and direction of effect, with selected SNPs that increase risk plotted on top. SNPs are coloured by their marginal p-values, and significant SNPs are shown in yellow.

#### Western hunter-gatherer ancestral path

The PALM results for the WHG ancestral path analysis of MS, using 62 LD-pruned markers, found no significant evidence for directional polygenic selection (p = 7.22e-5; ω = 0.021).

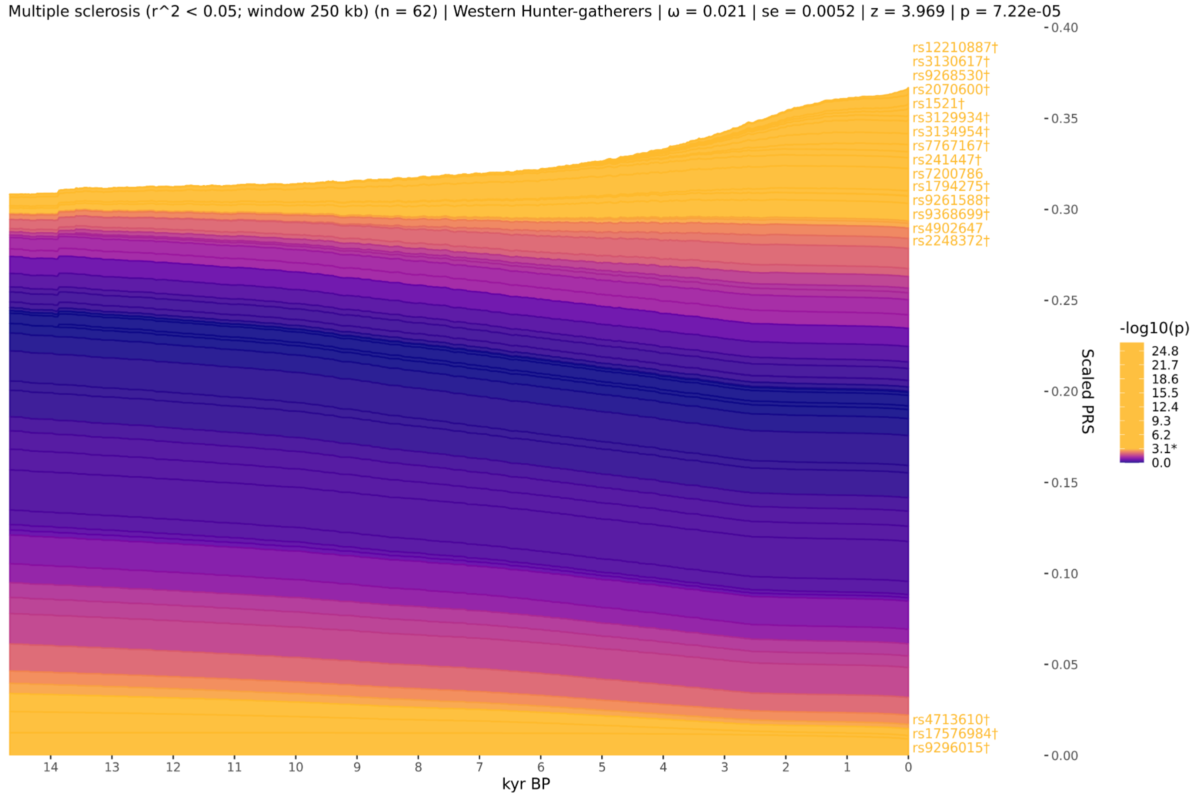


**Figure S6.2.** Stacked line plot of the WHG ancestry *PALM* analysis for Multiple sclerosis, showing the *CLUES* inferred allele frequency trajectories of each statistically independent SNP. Individual trajectories have been polarised to show the frequency of the positive risk allele, weighted by their scaled effect size. The y-axis shows the scaled polygenic risk score (PRS), which ranges from 0 to 1, representing the maximum possible additive genetic risk in a population. SNP trajectories are sorted by their *CLUES* p-values and direction of effect, with selected SNPs that increase risk plotted on top. SNPs are coloured by their marginal p-values, and significant SNPs are shown in yellow.

#### Eastern hunter-gatherer ancestral path

The PALM results for the EHG ancestral path analysis of MS, using 62 LD-pruned markers, found no significant evidence for directional polygenic selection (p = 2.60e-3; ω = 0.016).

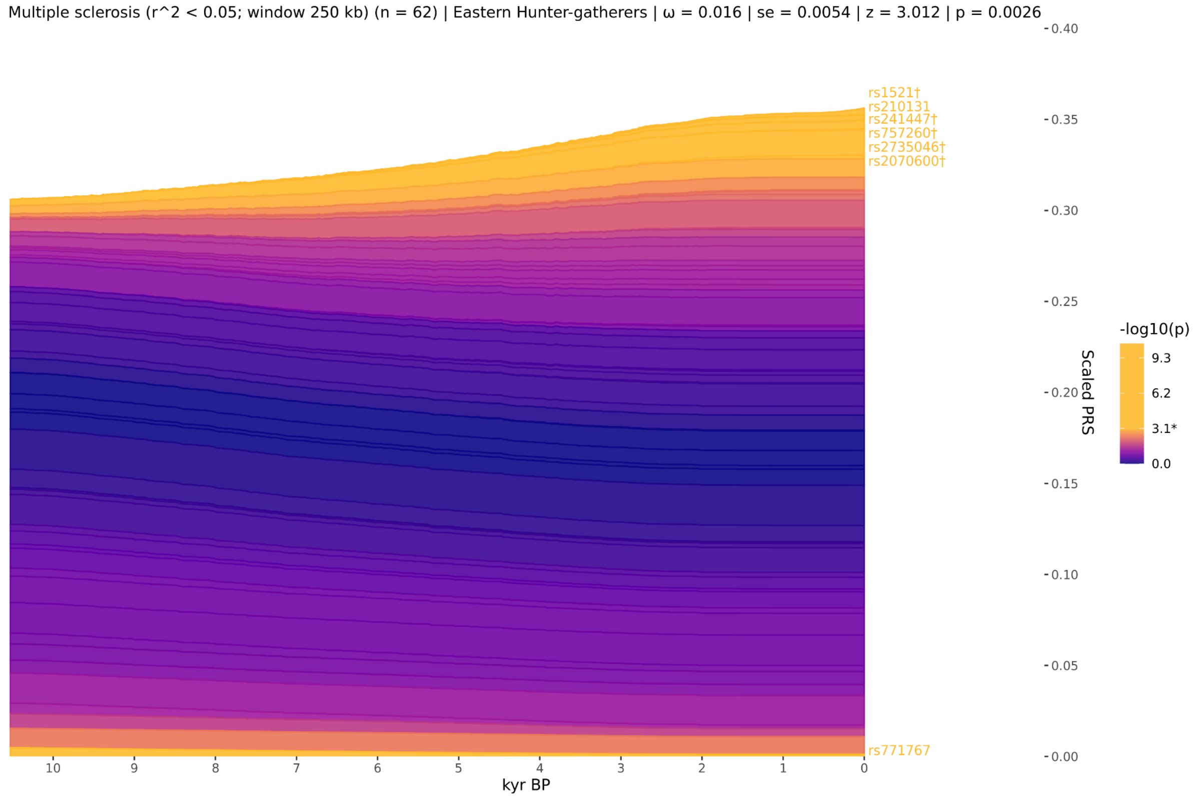


**Figure S6.3.** Stacked line plot of the EHG ancestry *PALM* analysis for Multiple sclerosis, showing the *CLUES* inferred allele frequency trajectories of each statistically independent SNP. Individual trajectories have been polarised to show the frequency of the positive risk allele, weighted by their scaled effect size. The y-axis shows the scaled polygenic risk score (PRS), which ranges from 0 to 1, representing the maximum possible additive genetic risk in a population. SNP trajectories are sorted by their *CLUES* p-values and direction of effect, with selected SNPs that increase risk plotted on top. SNPs are coloured by their marginal p-values, and significant SNPs are shown in yellow.

#### Caucasus hunter-gatherer ancestral path

The PALM results for the CHG ancestral path analysis of MS, using 62 LD-pruned markers, found statistically significant evidence for directional polygenic selection (p = 3.06e-2; ω = 0.009).

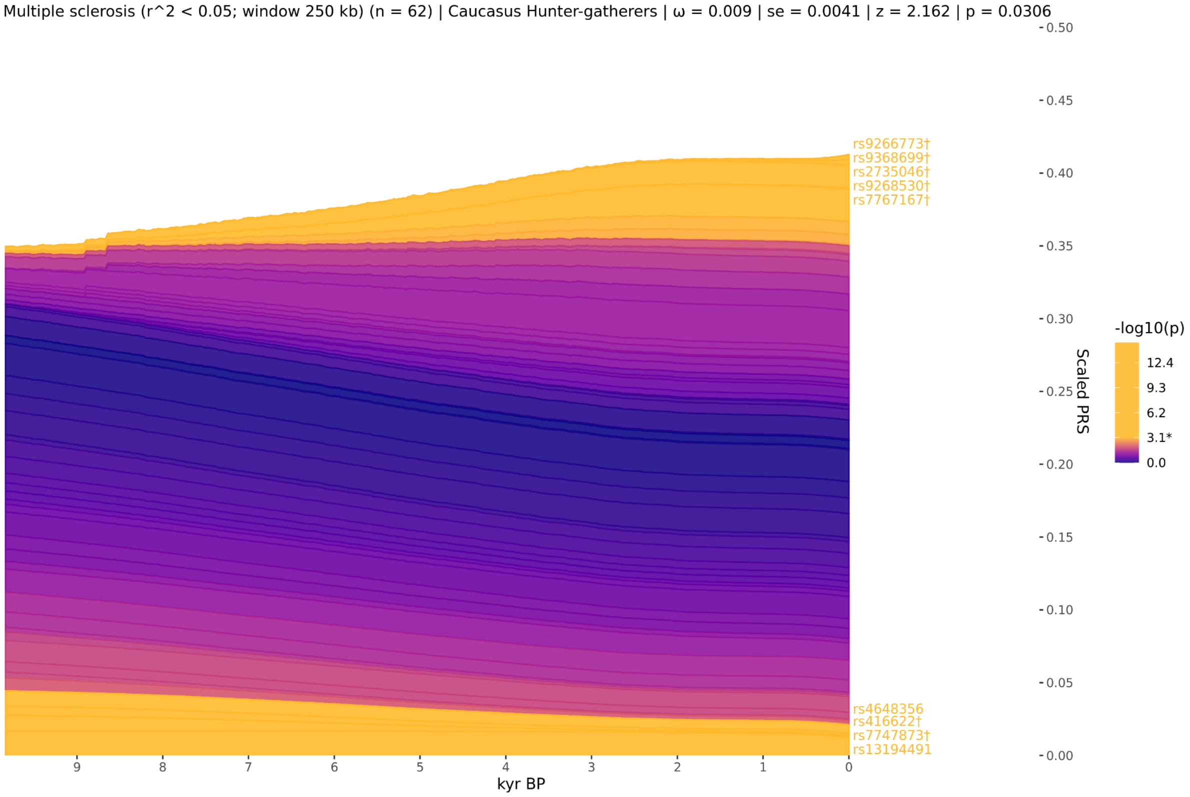


**Figure S6.4.** Stacked line plot of the CHG ancestry *PALM* analysis for Multiple sclerosis, showing the *CLUES* inferred allele frequency trajectories of each statistically independent SNP. Individual trajectories have been polarised to show the frequency of the positive risk allele, weighted by their scaled effect size. The y-axis shows the scaled polygenic risk score (PRS), which ranges from 0 to 1, representing the maximum possible additive genetic risk in a population. SNP trajectories are sorted by their *CLUES* p-values and direction of effect, with selected SNPs that increase risk plotted on top. SNPs are coloured by their marginal p-values, and significant SNPs are shown in yellow.

#### Anatolian farmer ancestral path

The PALM results for the ANA ancestral path analysis of MS, using 62 LD-pruned markers, found no significant evidence for directional polygenic selection (p = 6.43e-1; ω = 0.004).

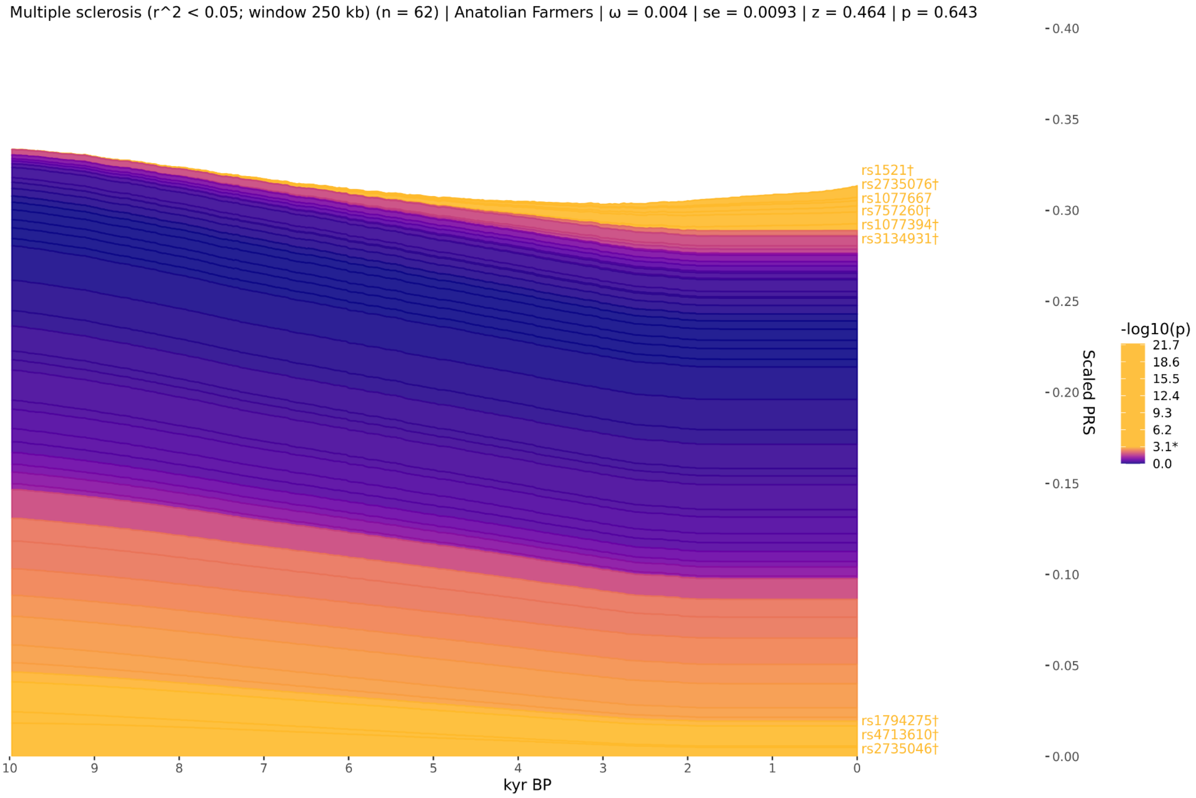


**Figure S6.5.** Stacked line plot of the ANA ancestry *PALM* analysis for Multiple sclerosis, showing the *CLUES* inferred allele frequency trajectories of each statistically independent SNP. Individual trajectories have been polarised to show the frequency of the positive risk allele, weighted by their scaled effect size. The y-axis shows the scaled polygenic risk score (PRS), which ranges from 0 to 1, representing the maximum possible additive genetic risk in a population. SNP trajectories are sorted by their *CLUES* p-values and direction of effect, with selected SNPs that increase risk plotted on top. SNPs are coloured by their marginal p-values, and significant SNPs are shown in yellow.

### Cross ancestry comparisons

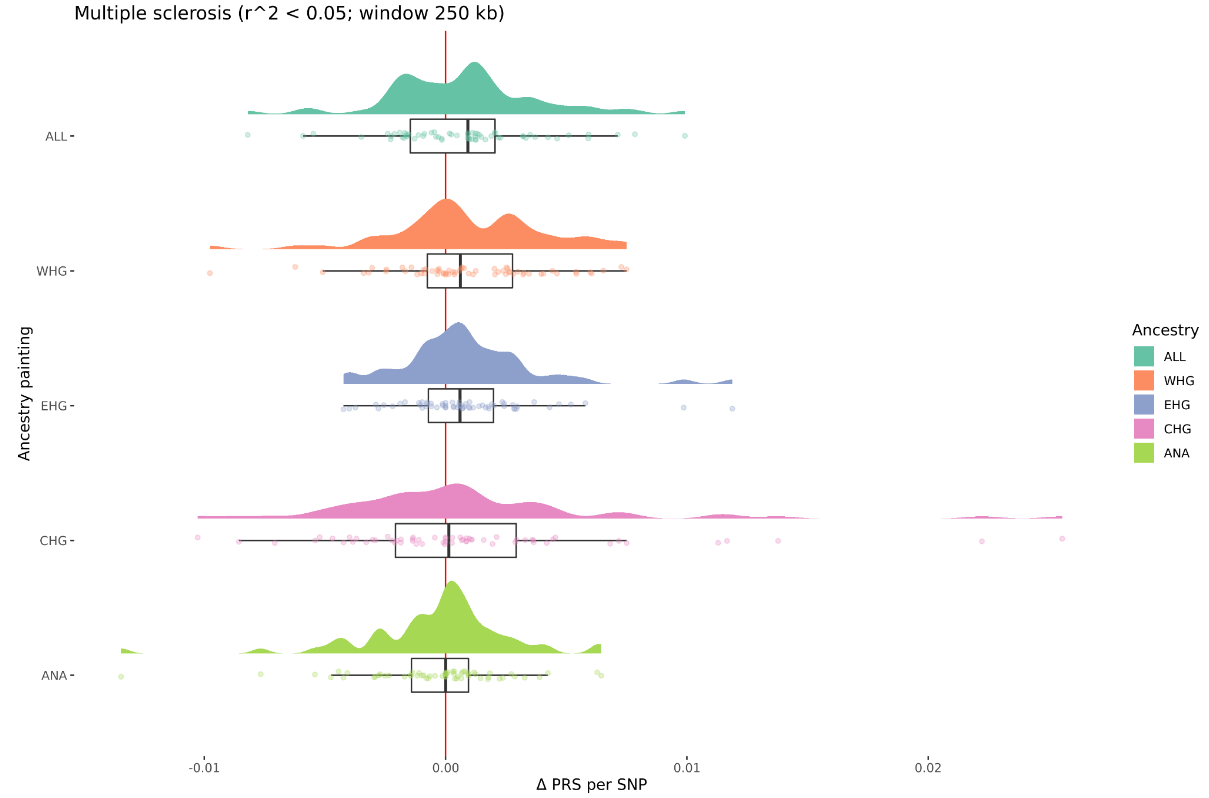


**Figure S6.6.** Density plots of the change over time in scaled PRS for each SNP in each marginal ancestry for Multiple sclerosis. Delta PRS per SNP is calculated from the *CLUES* models by taking the difference between the maximum likelihood estimates of the frequency of each SNP in the most recent and most ancient time points, weighted by the scaled effect size of the SNP in the focal trait.

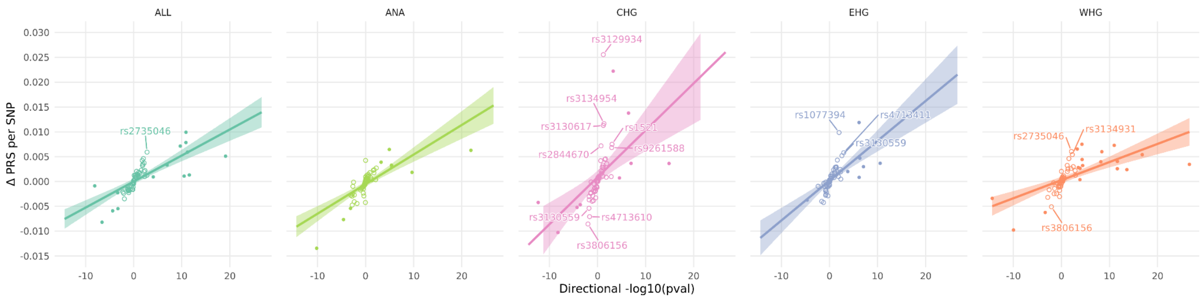


**Figure S6.7.** Scatter plots showing the delta PRS per SNP and the directional -log10(p-value) for each SNP in each marginal ancestry for Multiple sclerosis. Solid lines with shading show the linear regressions and standard errors. SNPs that do not achieve statistical significance in the marginal *CLUES* test but which have a large delta PRS are labelled as outliers.

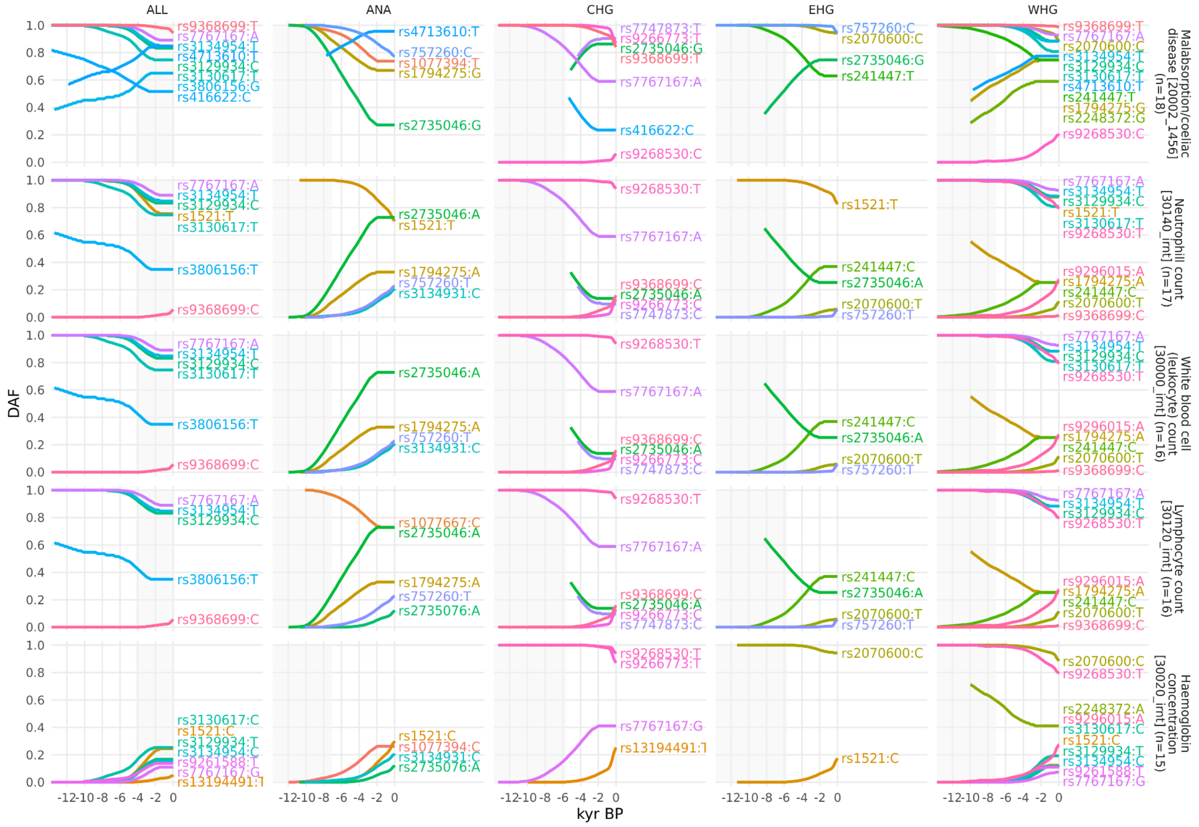


**Figure S6.8.** Allele frequency plots for positively selected MS-associated SNPs that are also associated with other phenotypes in the UK Biobank. Traits 1-5. SNPs are shown with their maximum likelihood trajectories, and polarised by the direction of their effect on the marginal UK Biobank trait (i.e. showing the 'risk' allele). Phenotypes are ordered according to the number of common SNPs, portions of the trajectory with low posterior density are cropped off, and the background is shaded for the approximate time period in which the ancestry existed as an actual population.

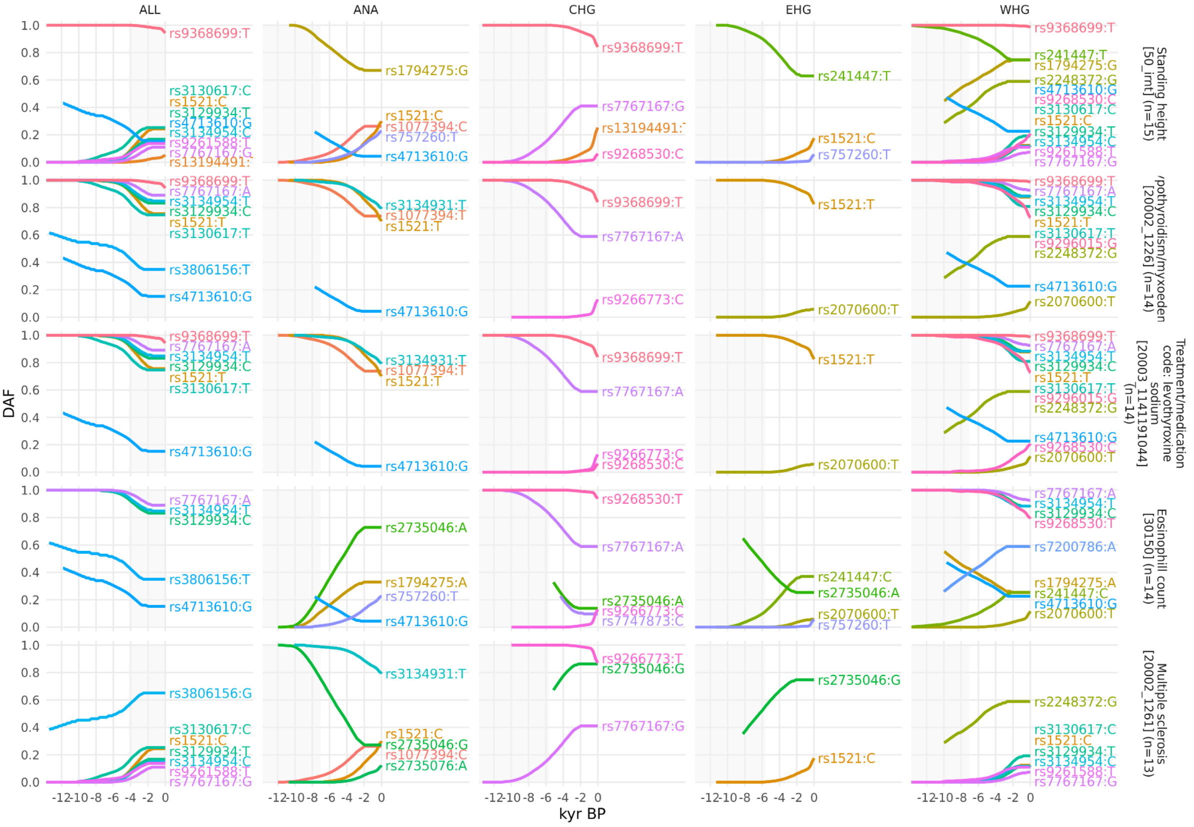


**Figure S6.9.** Allele frequency plots for positively selected MS-associated SNPs that are also associated with other phenotypes in the UK Biobank. Traits 6-10. SNPs are shown with their maximum likelihood trajectories, and polarised by the direction of their effect on the marginal UK Biobank trait (i.e. showing the 'risk' allele). Phenotypes are ordered according to the number of common SNPs, portions of the trajectory with low posterior density are cropped off, and the background is shaded for the approximate time period in which the ancestry existed as an actual population.

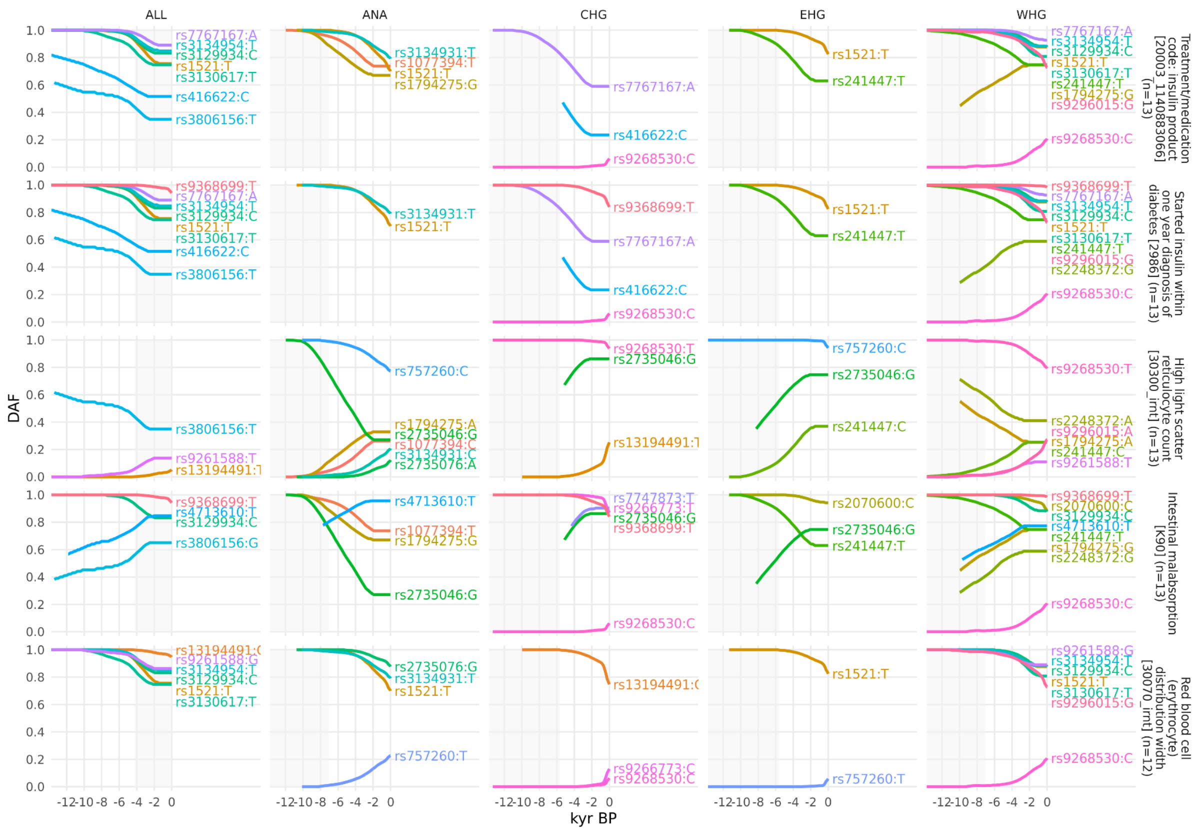


**Figure S6.10.** Allele frequency plots for positively selected MS-associated SNPs that are also associated with other phenotypes in the UK Biobank. Traits 11-15. SNPs are shown with their maximum likelihood trajectories, and polarised by the direction of their effect on the marginal UK Biobank trait (i.e. showing the 'risk' allele). Phenotypes are ordered according to the number of common SNPs, portions of the trajectory with low posterior density are cropped off, and the background is shaded for the approximate time period in which the ancestry existed as an actual population.

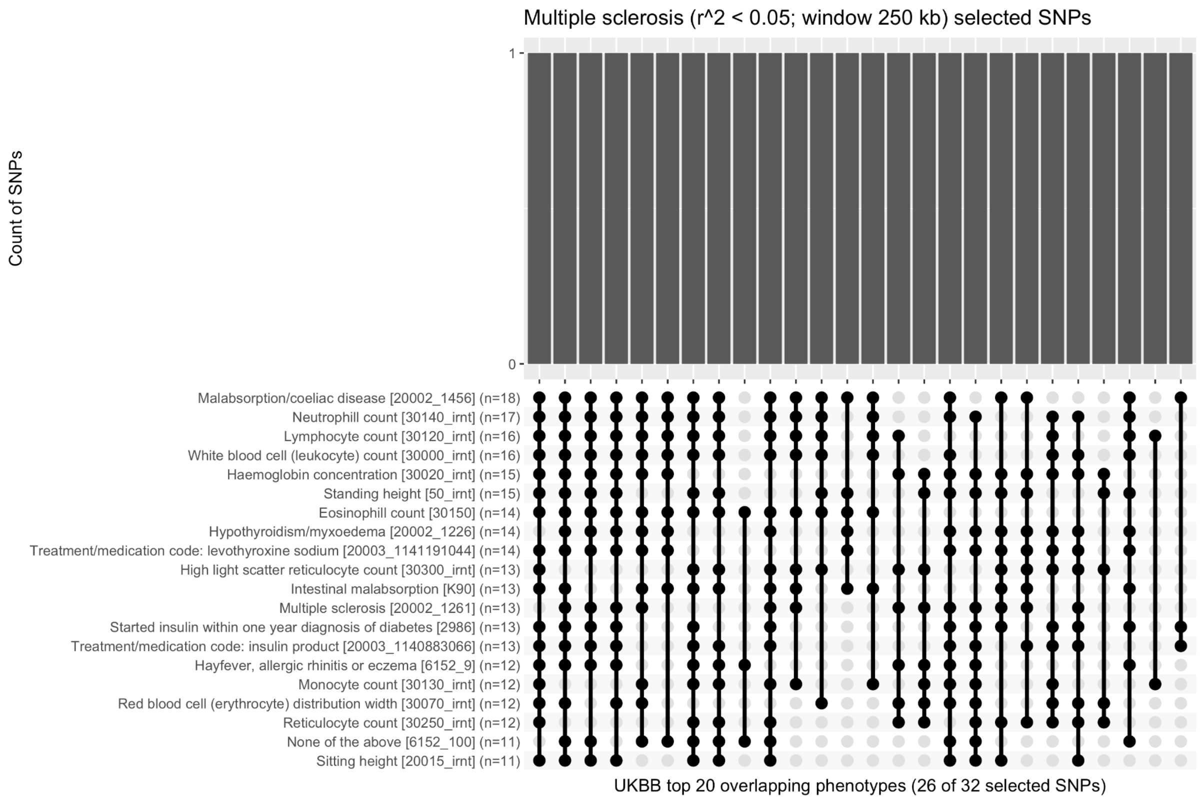


**Figure S6.11.** Upset plot showing a histogram of selected MS-associated SNPs which share a pleiotropic association with one or more marginal phenotypes in the UK Biobank. Top 20 traits shown. Of the 32 selected MS-associated SNPs, 26 (81%) are also associated with one or more of the top 20 genetically correlated phenotypes.

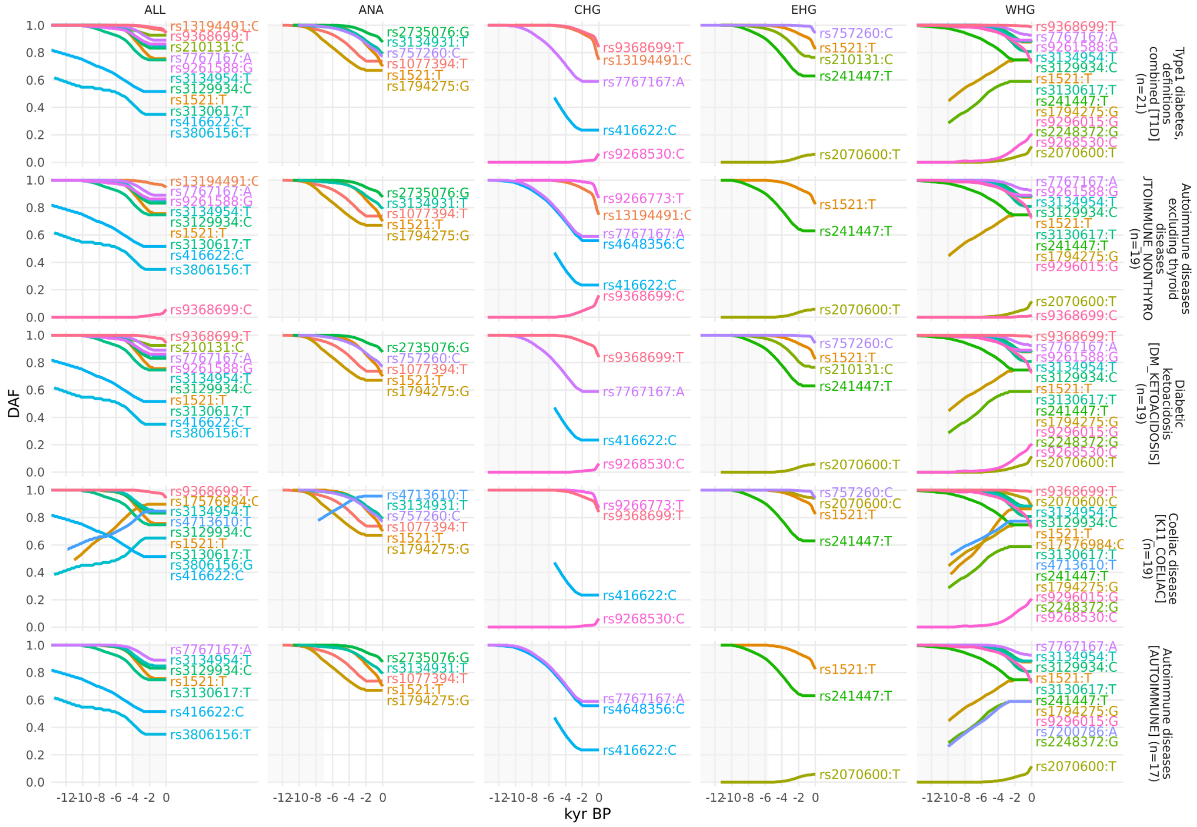


**Figure S6.12.** Allele frequency plots for positively selected MS-associated SNPs that are also associated with other phenotypes in the FinnGen study. Traits 1-5. SNPs are shown with their maximum likelihood trajectories, and polarised by the direction of their effect on the marginal FinnGen trait (i.e. showing the ‘risk’ allele). Phenotypes are ordered according to the number of common SNPs, portions of the trajectory with low posterior density are cropped off, and the background is shaded for the approximate time period in which the ancestry existed as an actual population.

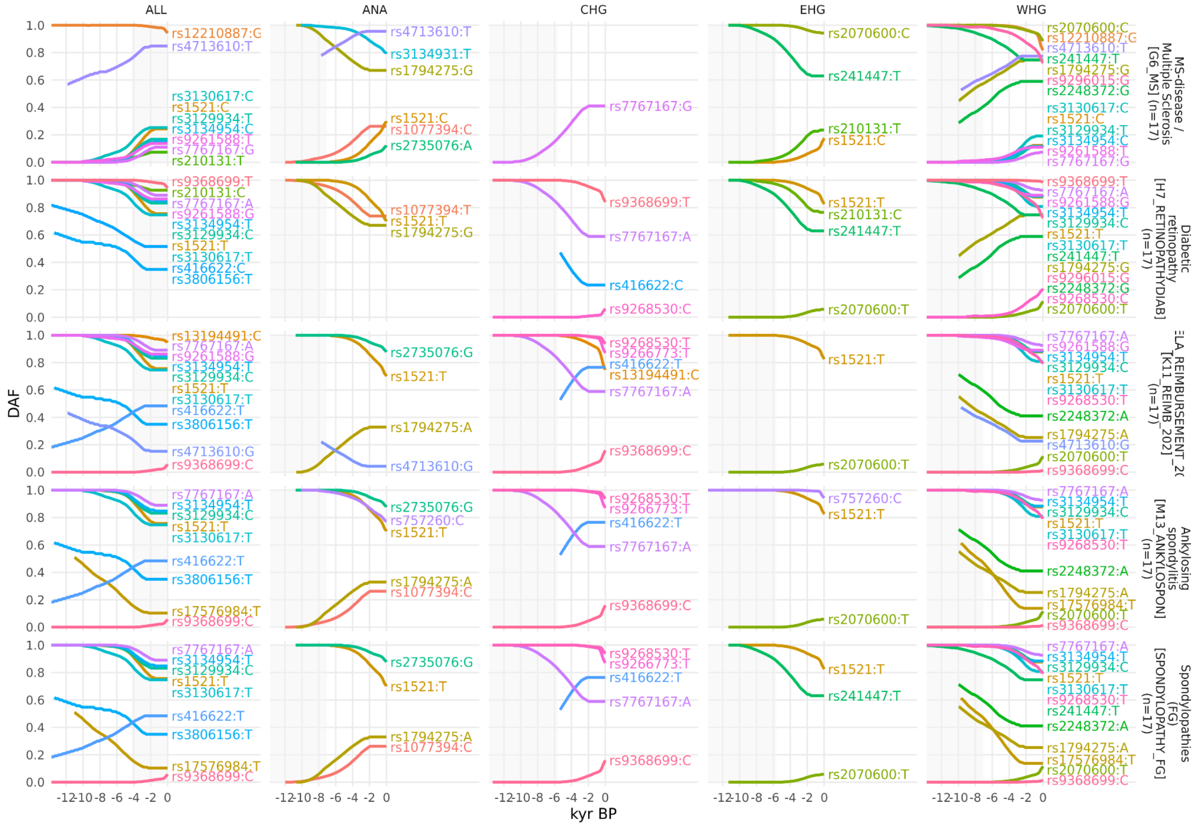

**Figure S6.13.** Allele frequency plots for positively selected MS-associated SNPs that are also associated with other phenotypes in the FinnGen study. Traits 6-10. SNPs are shown with their maximum likelihood trajectories, and polarised by the direction of their effect on the marginal FinnGen trait (i.e. showing the 'risk' allele). Phenotypes are ordered according to the number of common SNPs, portions of the trajectory with low posterior density are cropped off, and the background is shaded for the approximate time period in which the ancestry existed as an actual population.

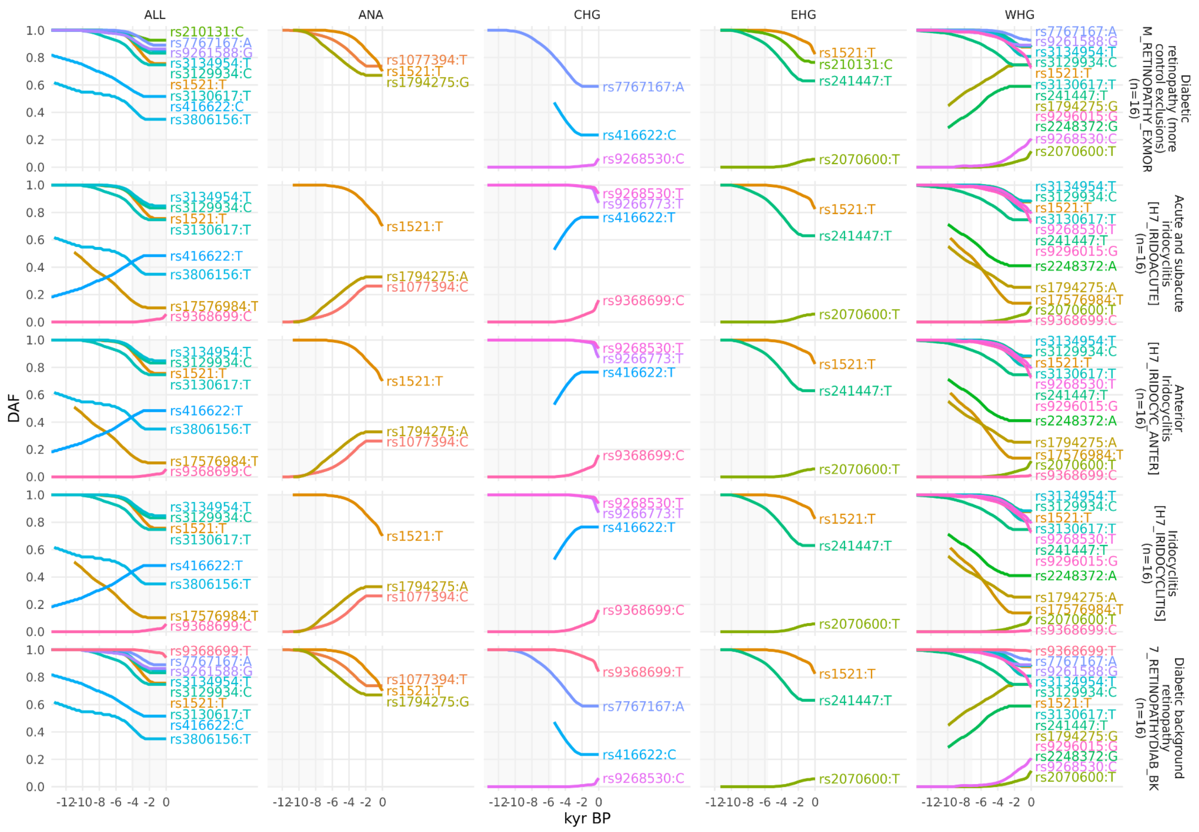


**Figure S6.14.** Allele frequency plots for positively selected MS-associated SNPs that are also associated with other phenotypes in the FinnGen study. Traits 11-15. SNPs are shown with their maximum likelihood trajectories, and polarised by the direction of their effect on the marginal FinnGen trait (i.e. showing the 'risk' allele). Phenotypes are ordered according to the number of common SNPs, portions of the trajectory with low posterior density are cropped off, and the background is shaded for the approximate time period in which the ancestry existed as an actual population.

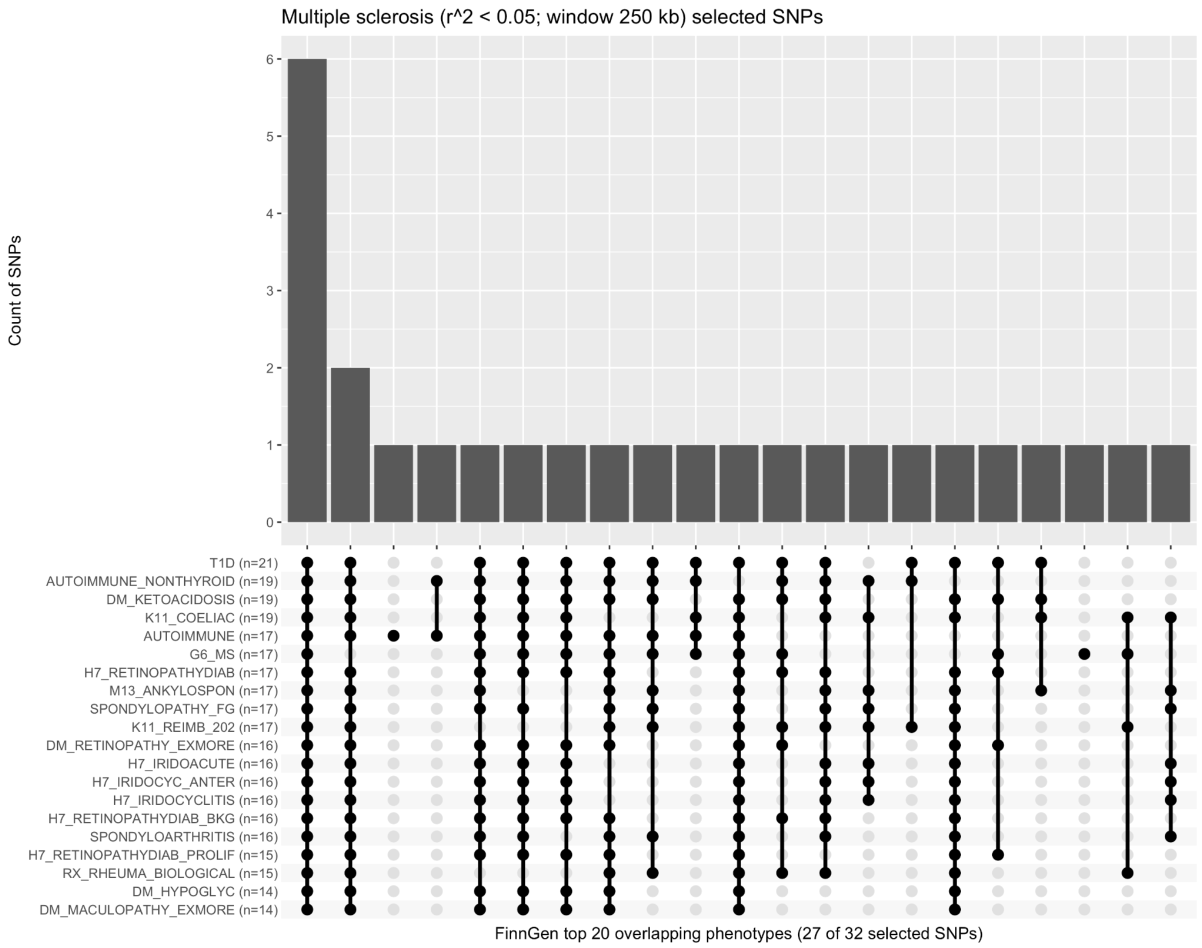


**Figure S6.15.** Upset plot showing a histogram of selected MS-associated SNPs which share a pleiotropic association with one or more marginal phenotypes in the FinnGen study. Top 20 traits shown. Of the 32 selected MS-associated SNPs, 27 (84%) are also associated with one or more of the top 20 genetically correlated phenotypes.

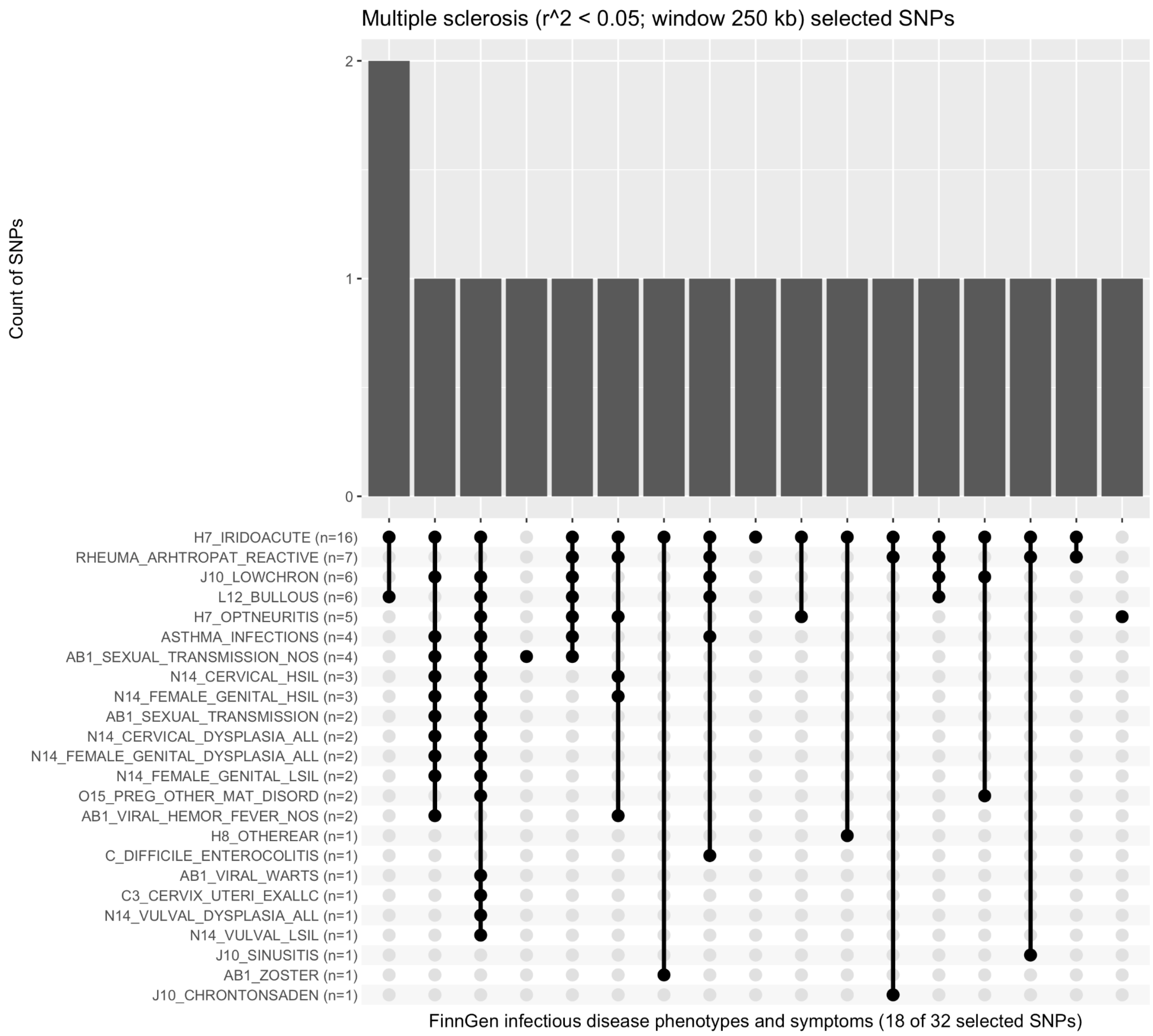


**Figure S6.16.** Upset plot showing a histogram of selected MS-associated SNPs which share a pleiotropic association with one or more infectious disease phenotypes and symptoms in the FinnGen study. Of the 32 selected MS-associated SNPs, 18 (56%) are also associated with one or more infectious disease phenotypes.

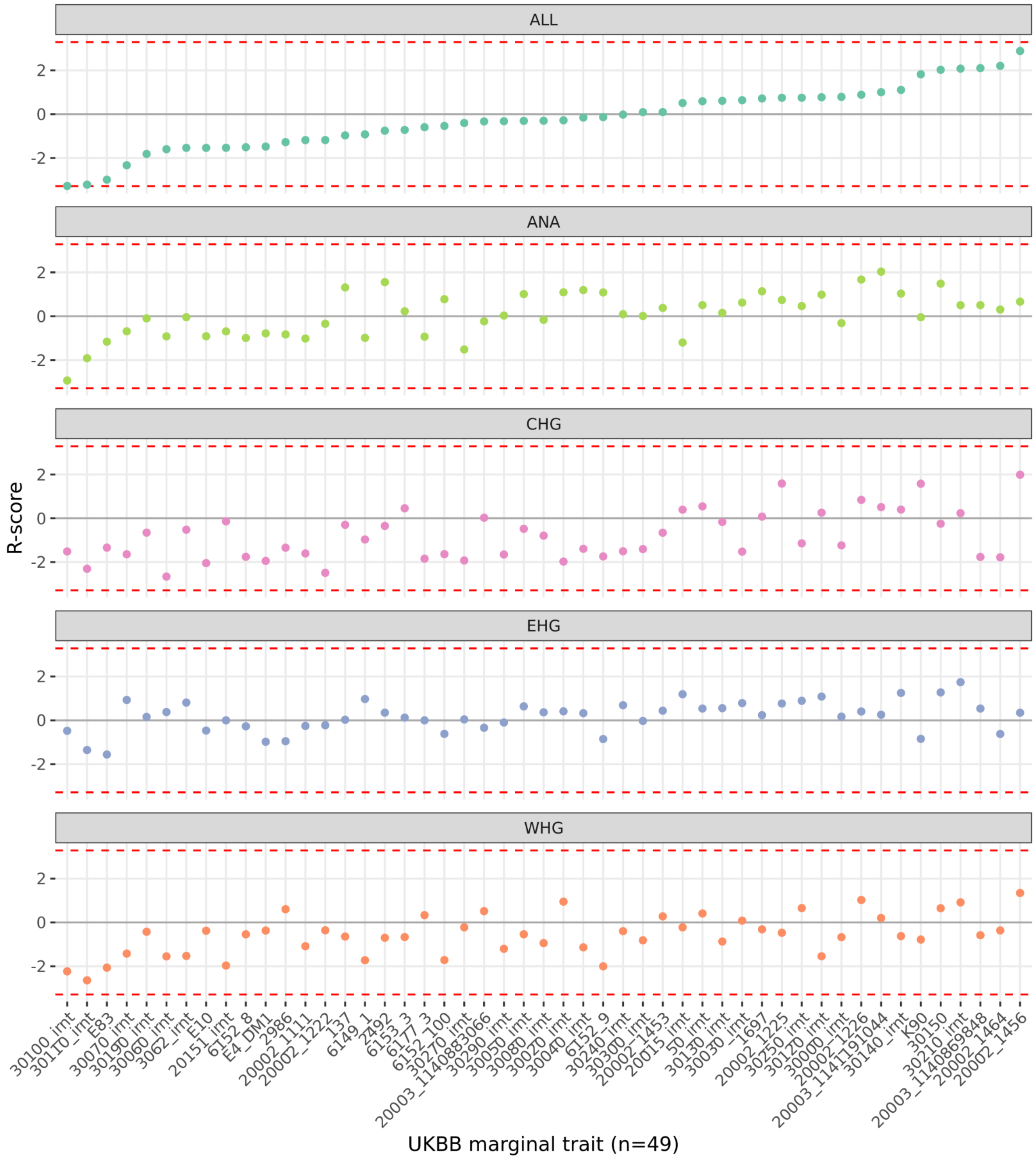


**Figure S6.17.** Dot plot showing the R-scores from the J-PALM tests of MS against each of 49 overlapping traits in UK Biobank. The dotted red lines show the Bonferroni corrected significance threshold. No marginal trait significantly attenuates the signal of selection seen in MS.

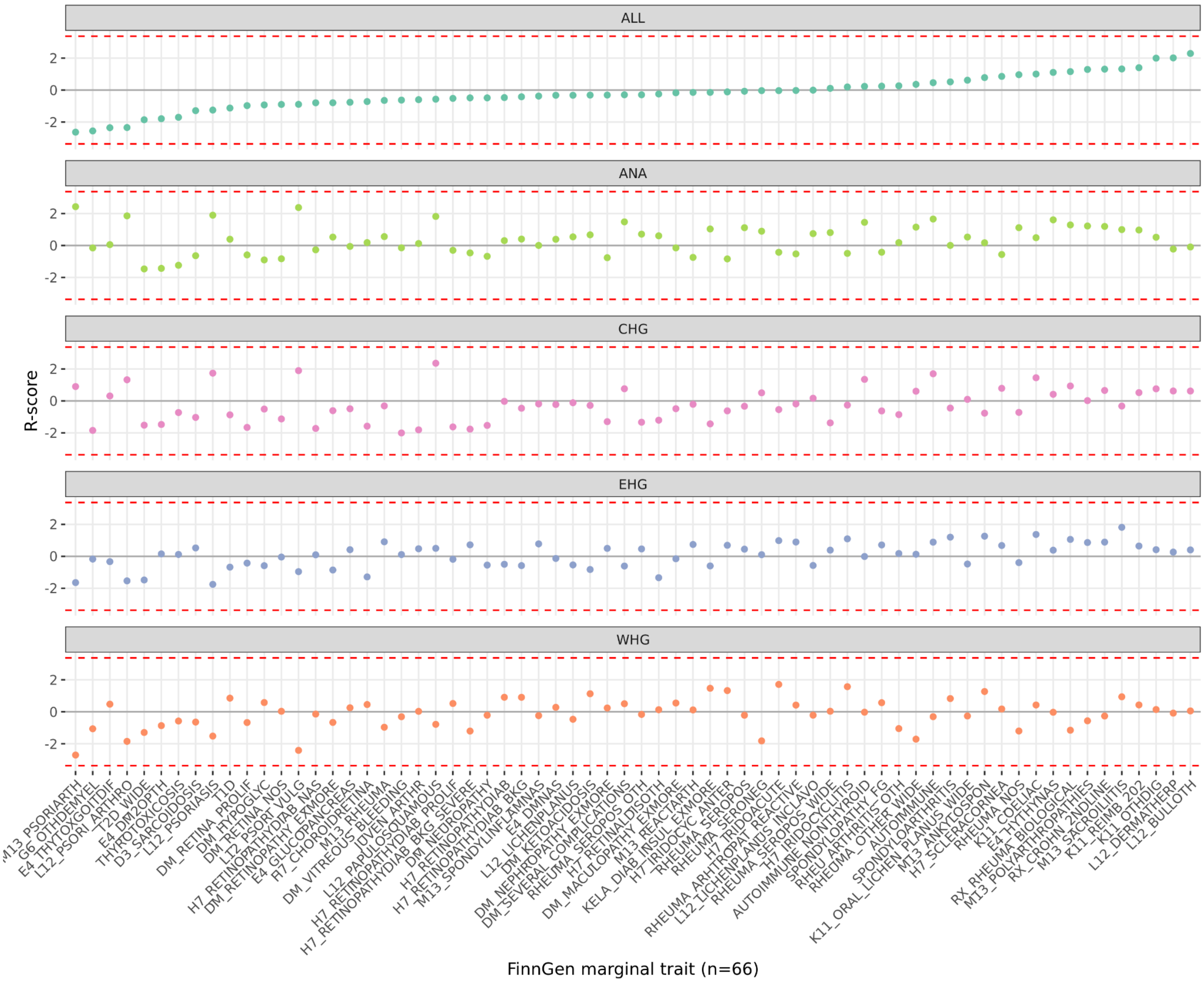


**Figure S6.18.** Dot plot showing the R-scores from the J-PALM tests of MS against each of 66 overlapping traits in FinnGen. The dotted red lines show the Bonferroni corrected significance threshold. No marginal trait significantly attenuates the signal of selection seen in MS.

#### Rheumatoid arthritis

The CLUES results for all genome-wide significant RA associations are available in ST10, and the results for the subset of statistically independent markers used in the PALM analysis are available in ST9.

##### Pan-ancestry analysis

The PALM results for the pan-ancestry analysis of RA, using 153 LD-pruned markers, found statistically significant evidence for directional polygenic selection (p = 3.26e-3; ω = -0.007).

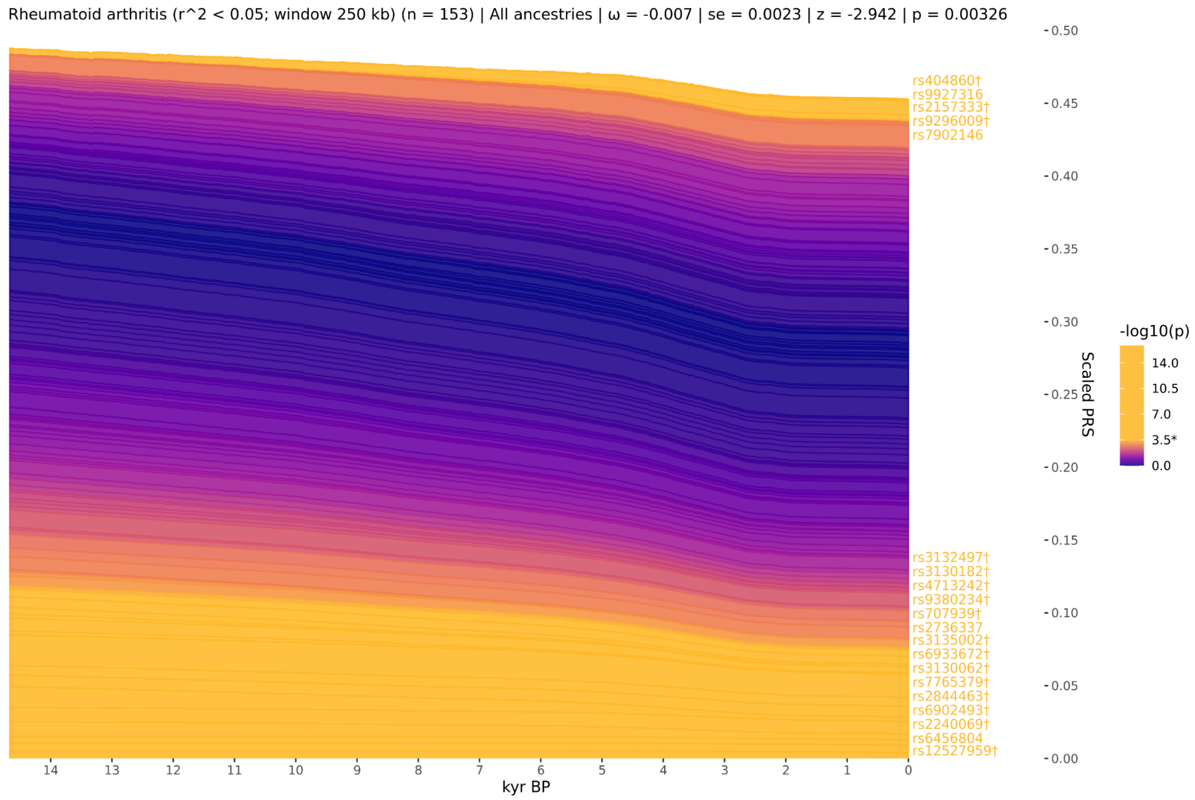


**Figure S6.19.** Stacked line plot of the pan-ancestry *PALM* analysis for Rheumatoid arthritis, showing the *CLUES* inferred allele frequency trajectories of each statistically independent SNP. Individual trajectories have been polarised to show the frequency of the positive risk allele, weighted by their scaled effect size. The y-axis shows the scaled polygenic risk score (PRS), which ranges from 0 to 1, representing the maximum possible additive genetic risk in a population. SNP trajectories are sorted by their *CLUES* p-values and direction of effect, with selected SNPs that increase risk plotted on top. SNPs are coloured by their marginal p-values, and significant SNPs are shown in yellow.

#### Western hunter-gatherer ancestral path

The PALM results for the WHG ancestral path analysis of RA, using 153 LD-pruned markers, found no significant evidence for directional polygenic selection (p = 8.63e-1; ω = -0.001).

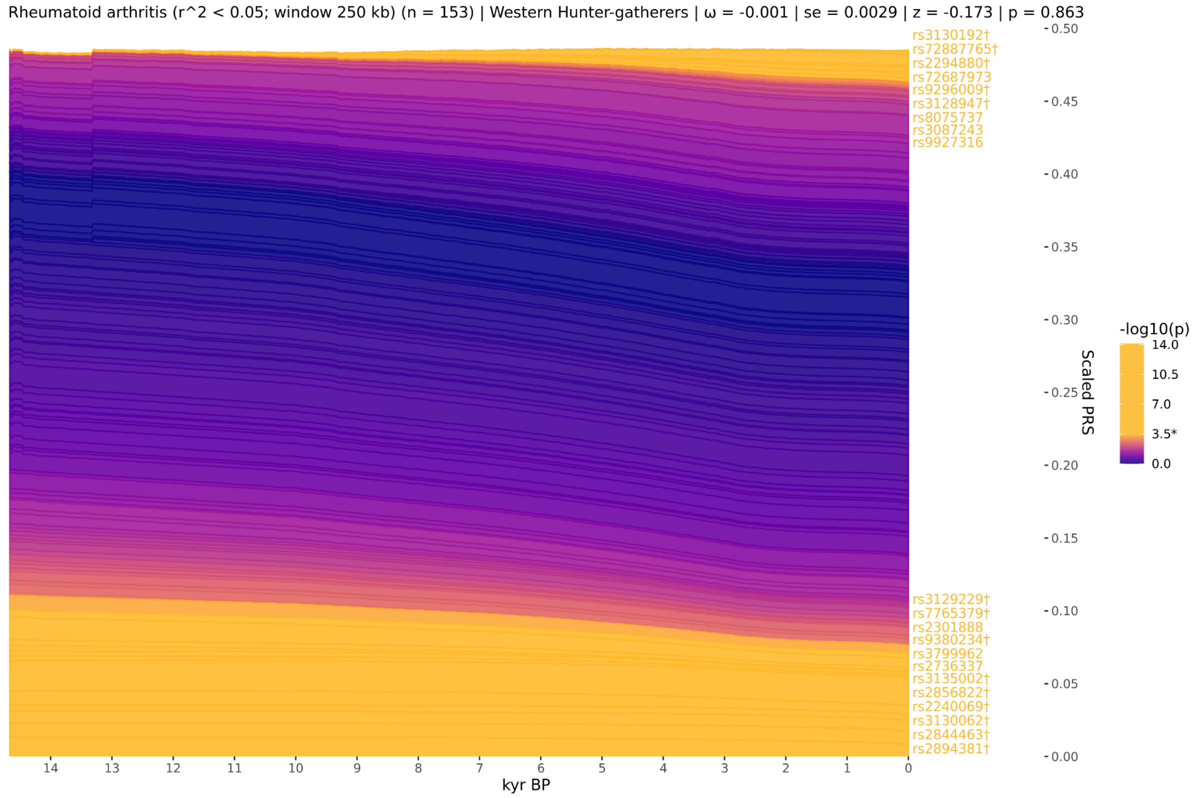


**Figure S6.20.** Stacked line plot of the WHG ancestry *PALM* analysis for Rheumatoid arthritis, showing the *CLUES* inferred allele frequency trajectories of each statistically independent SNP. Individual trajectories have been polarised to show the frequency of the positive risk allele, weighted by their scaled effect size. The y-axis shows the scaled polygenic risk score (PRS), which ranges from 0 to 1, representing the maximum possible additive genetic risk in a population. SNP trajectories are sorted by their *CLUES* p-values and direction of effect, with selected SNPs that increase risk plotted on top. SNPs are coloured by their marginal p-values, and significant SNPs are shown in yellow.

#### Eastern hunter-gatherer ancestral path

The PALM results for the EHG ancestral path analysis of RA, using 153 LD-pruned markers, found no significant evidence for directional polygenic selection ($p = 6.24e\text{-}1$; $\omega = -0.003$).

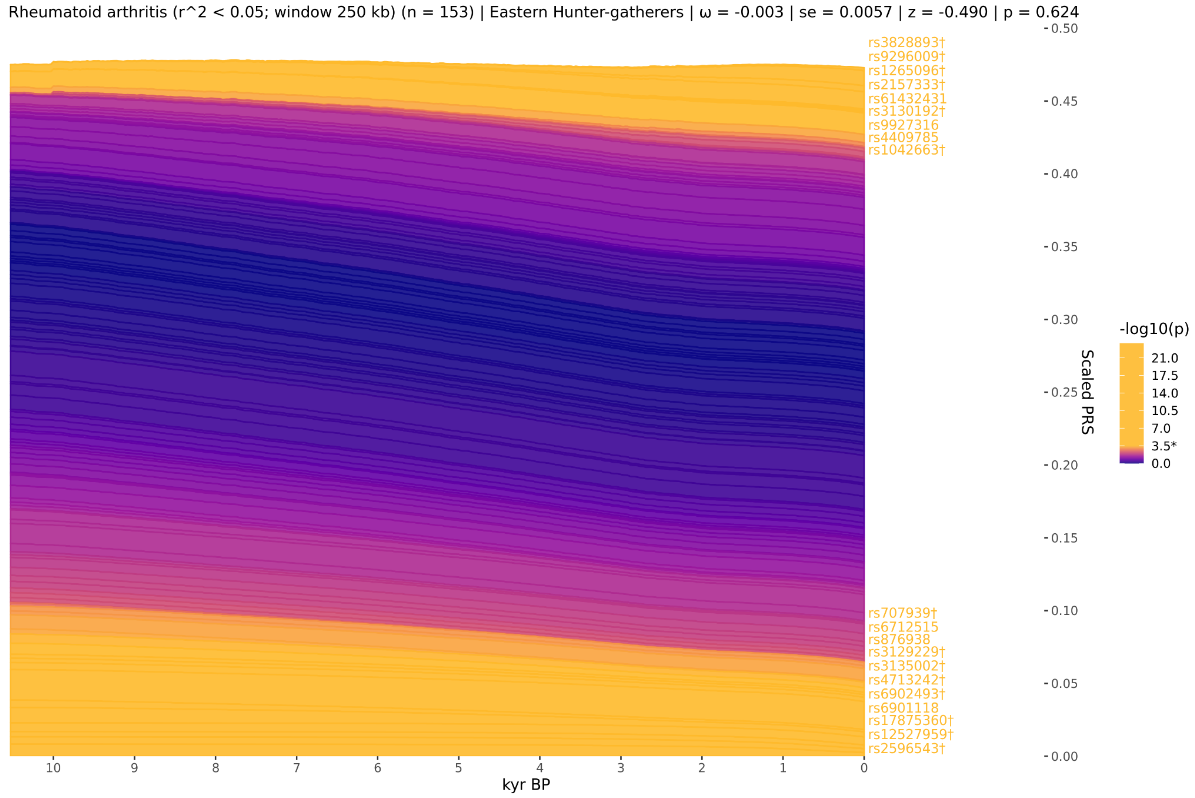


**Figure S6.21.** Stacked line plot of the EHG ancestry *PALM* analysis for Rheumatoid arthritis, showing the *CLUES* inferred allele frequency trajectories of each statistically independent SNP. Individual trajectories have been polarised to show the frequency of the positive risk allele, weighted by their scaled effect size. The y-axis shows the scaled polygenic risk score (PRS), which ranges from 0 to 1, representing the maximum possible additive genetic risk in a population. SNP trajectories are sorted by their *CLUES* p-values and direction of effect, with selected SNPs that increase risk plotted on top. SNPs are coloured by their marginal p-values, and significant SNPs are shown in yellow.

#### Caucasus hunter-gatherer ancestral path

The PALM results for the CHG ancestral path analysis of RA, using 153 LD-pruned markers, found statistically significant evidence for directional polygenic selection (p = 6.33e-2; ω = -0.014).

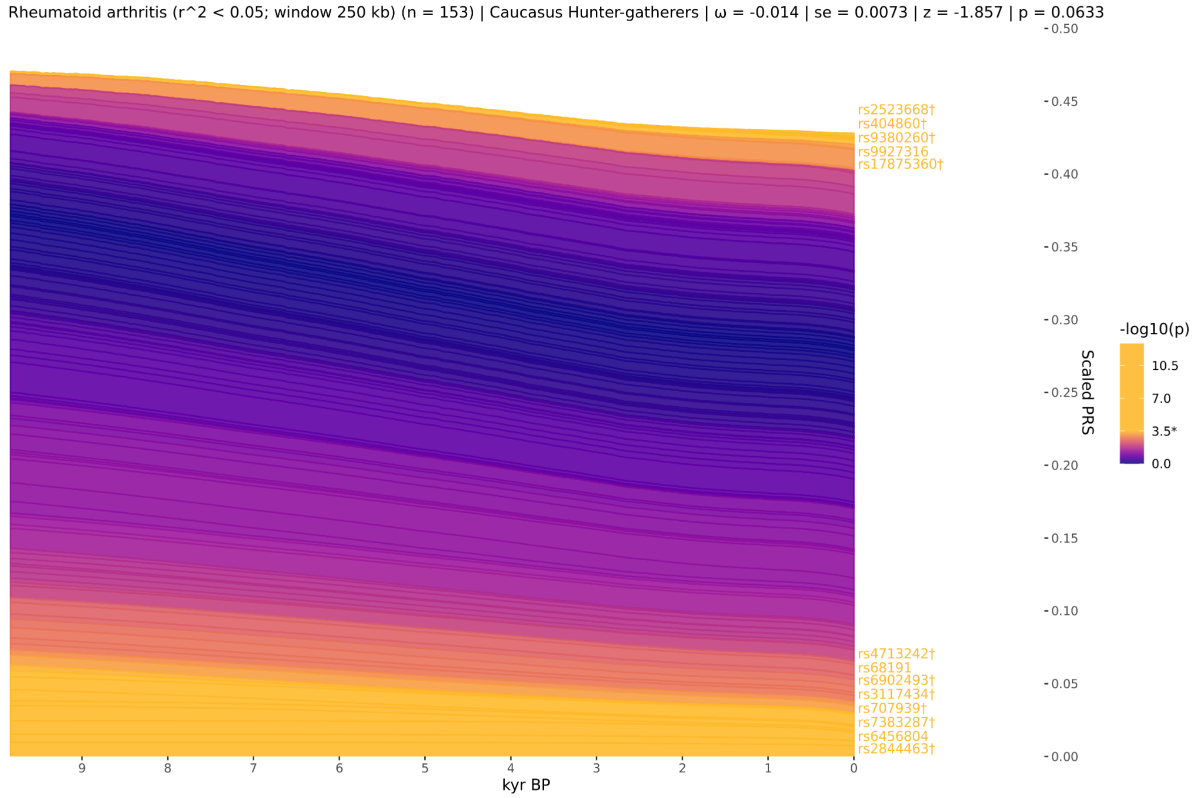


**Figure S6.22.** Stacked line plot of the CHG ancestry *PALM* analysis for Rheumatoid arthritis, showing the *CLUES* inferred allele frequency trajectories of each statistically independent SNP. Individual trajectories have been polarised to show the frequency of the positive risk allele, weighted by their scaled effect size. The y-axis shows the scaled polygenic risk score (PRS), which ranges from 0 to 1, representing the maximum possible additive genetic risk in a population. SNP trajectories are sorted by their *CLUES* p-values and direction of effect, with selected SNPs that increase risk plotted on top. SNPs are coloured by their marginal p-values, and significant SNPs are shown in yellow.

#### Anatolian farmer ancestral path

The PALM results for the ANA ancestral path analysis of RA, using 153 LD-pruned markers, found statistically significant evidence for directional polygenic selection (p = 1.49e-1; ω = -0.006).

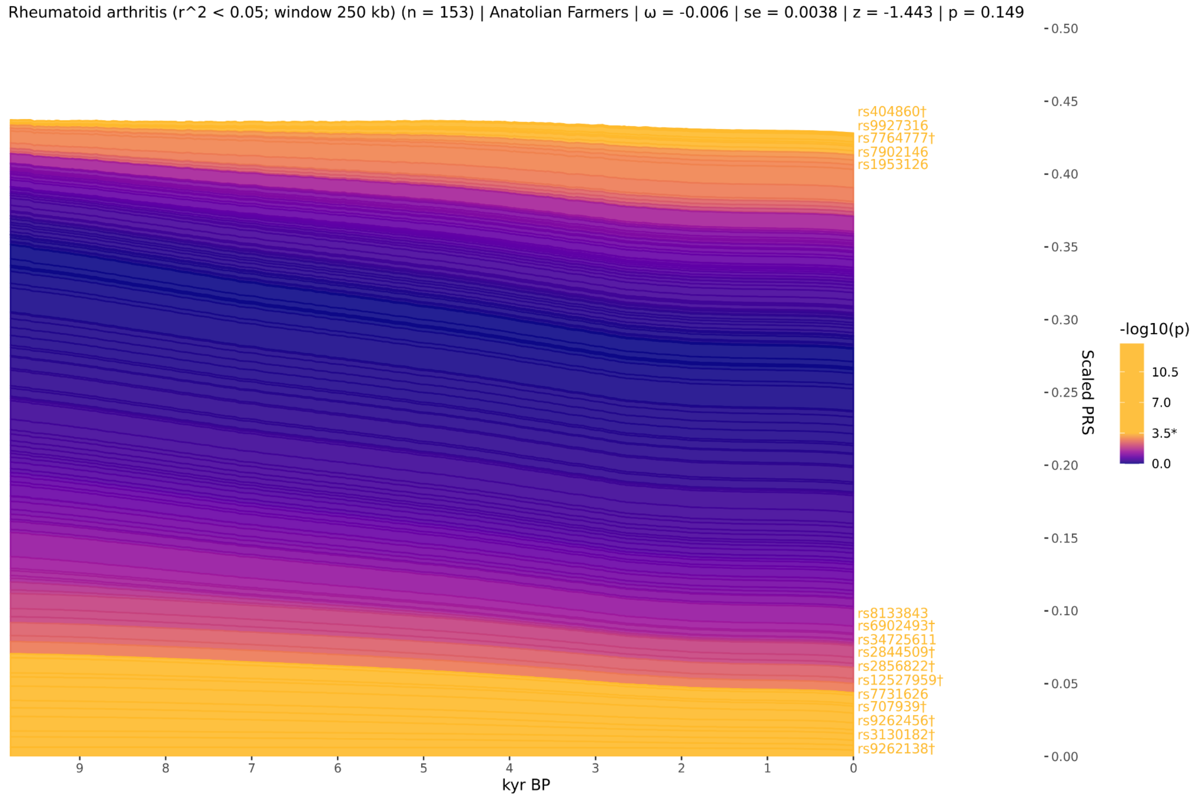


**Figure S6.23.** Stacked line plot of the ANA ancestry *PALM* analysis for Rheumatoid arthritis, showing the *CLUES* inferred allele frequency trajectories of each statistically independent SNP. Individual trajectories have been polarised to show the frequency of the positive risk allele, weighted by their scaled effect size. The y-axis shows the scaled polygenic risk score (PRS), which ranges from 0 to 1, representing the maximum possible additive genetic risk in a population. SNP trajectories are sorted by their *CLUES* p-values and direction of effect, with selected SNPs that increase risk plotted on top. SNPs are coloured by their marginal p-values, and significant SNPs are shown in yellow.

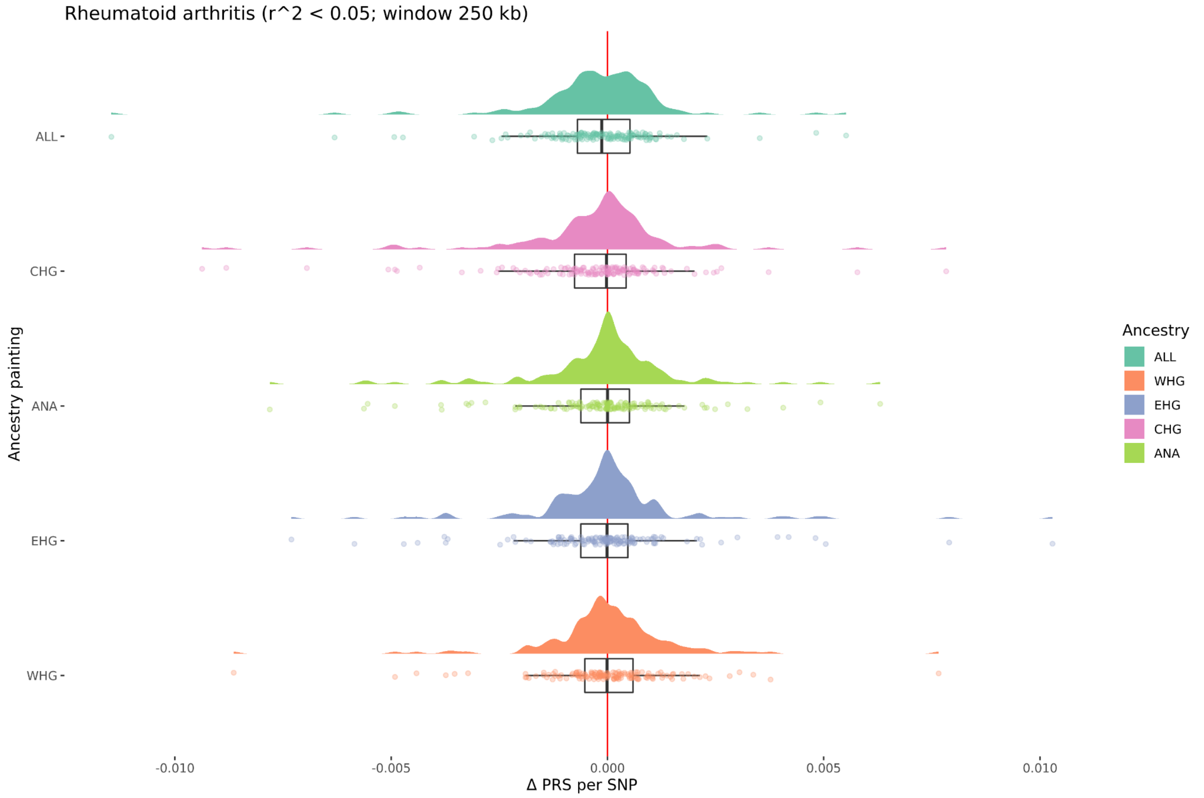


**Figure S6.24.** Density plots of the change over time in scaled PRS for each SNP in each marginal ancestry for Rheumatoid arthritis. Delta PRS per SNP is calculated from the *CLUES* models by taking the difference between the maximum likelihood estimates of the frequency of each SNP in the most recent and most ancient time points, weighted by the scaled effect size of the SNP in the focal trait.

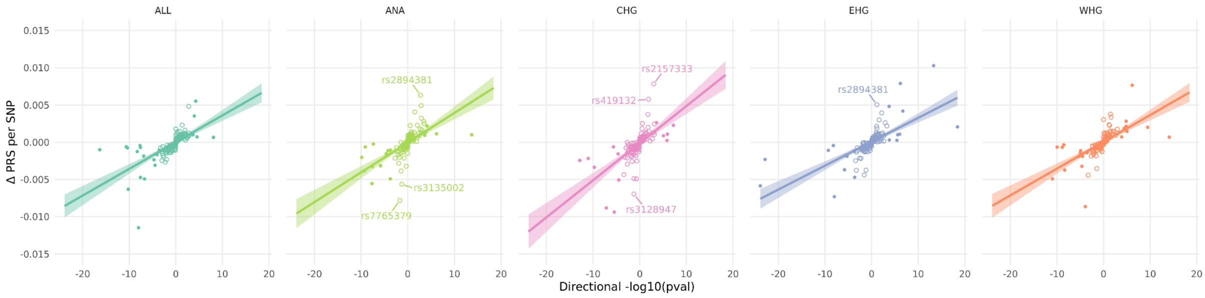


**Figure S6.25.** Scatter plots showing the delta PRS per SNP and the directional -log10(p-value) for each SNP in each marginal ancestry for Rheumatoid arthritis. Solid lines with shading show the linear regressions and standard errors. SNPs that do not achieve statistical significance in the marginal *CLUES* test but which have a large delta PRS are labelled as outliers.

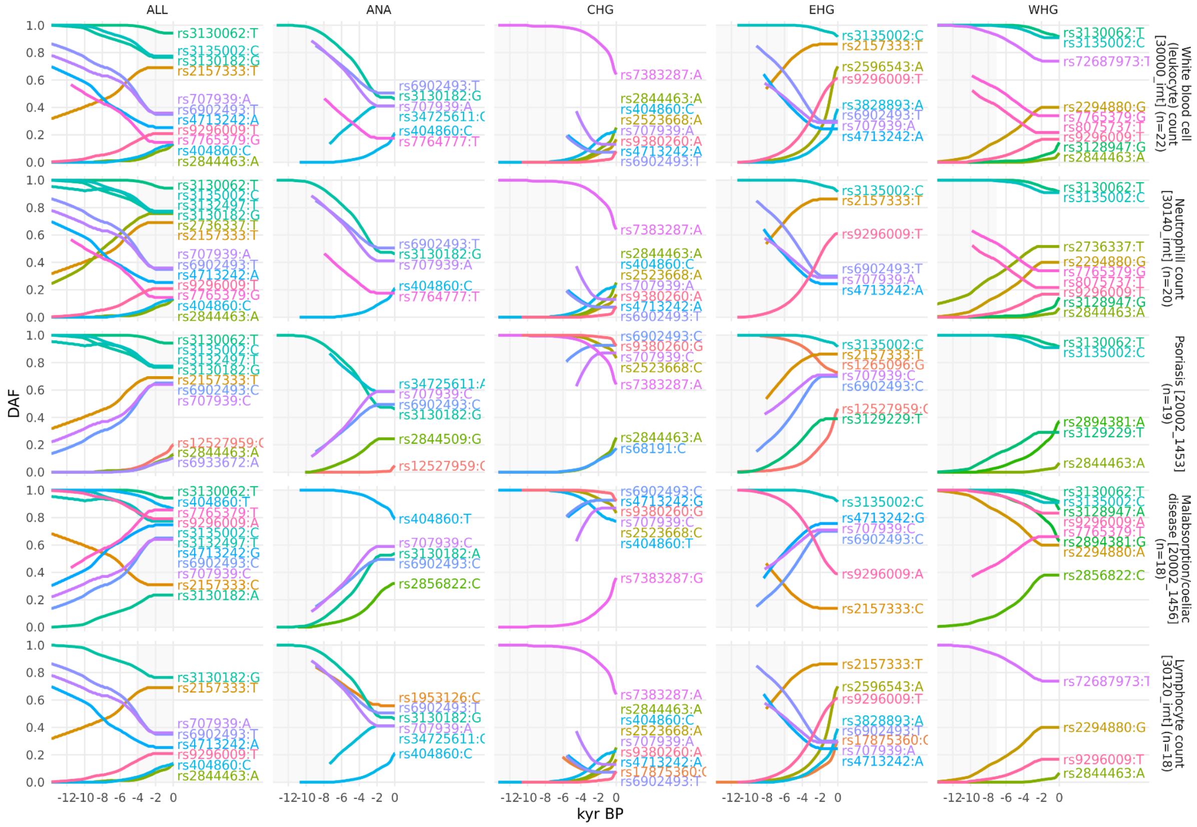


**Figure S6.26.** Allele frequency plots for positively selected RA-associated SNPs that are also associated with other phenotypes in the UK Biobank. Traits 1-5. SNPs are shown with their maximum likelihood trajectories, and polarised by the direction of their effect on the marginal UK Biobank trait (i.e. showing the 'risk' allele). Phenotypes are ordered according to the number of common SNPs, portions of the trajectory with low posterior density are cropped off, and the background is shaded for the approximate time period in which the ancestry existed as an actual population.

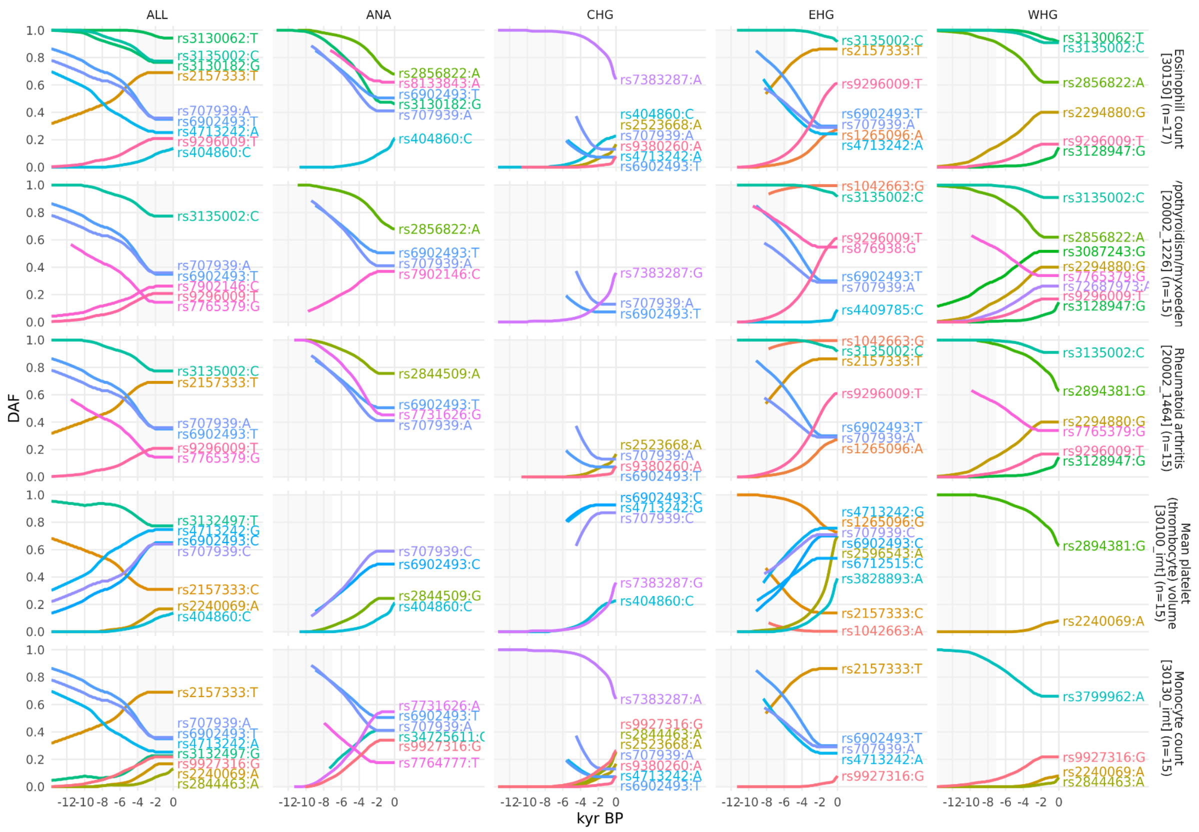


**Figure S6.27.** Allele frequency plots for positively selected RA-associated SNPs that are also associated with other phenotypes in the UK Biobank. Traits 6-10. SNPs are shown with their maximum likelihood trajectories, and polarised by the direction of their effect on the marginal UK Biobank trait (i.e. showing the 'risk' allele). Phenotypes are ordered according to the number of common SNPs, portions of the trajectory with low posterior density are cropped off, and the background is shaded for the approximate time period in which the ancestry existed as an actual population.

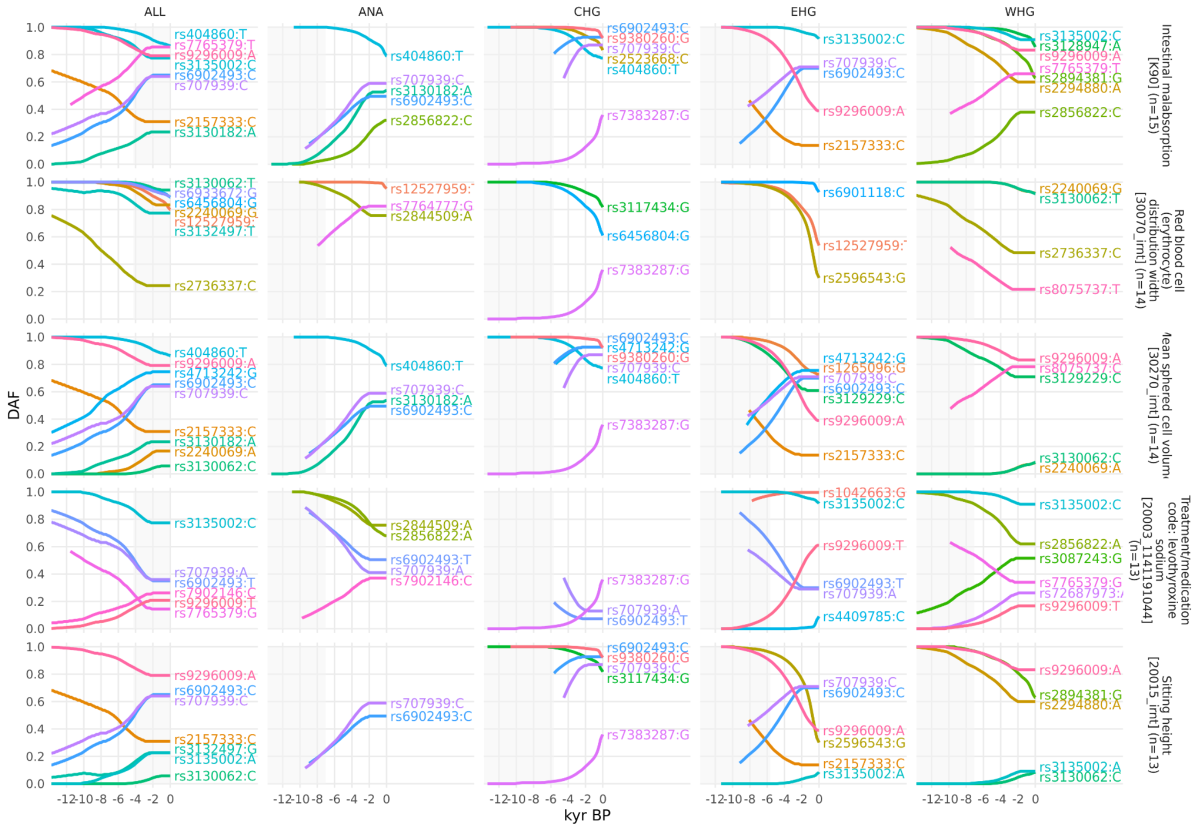

**Figure S6.28.** Allele frequency plots for positively selected RA-associated SNPs that are also associated with other phenotypes in the UK Biobank. Traits 11-15. SNPs are shown with their maximum likelihood trajectories, and polarised by the direction of their effect on the marginal UK Biobank trait (i.e. showing the 'risk' allele). Phenotypes are ordered according to the number of common SNPs, portions of the trajectory with low posterior density are cropped off, and the background is shaded for the approximate time period in which the ancestry existed as an actual population.

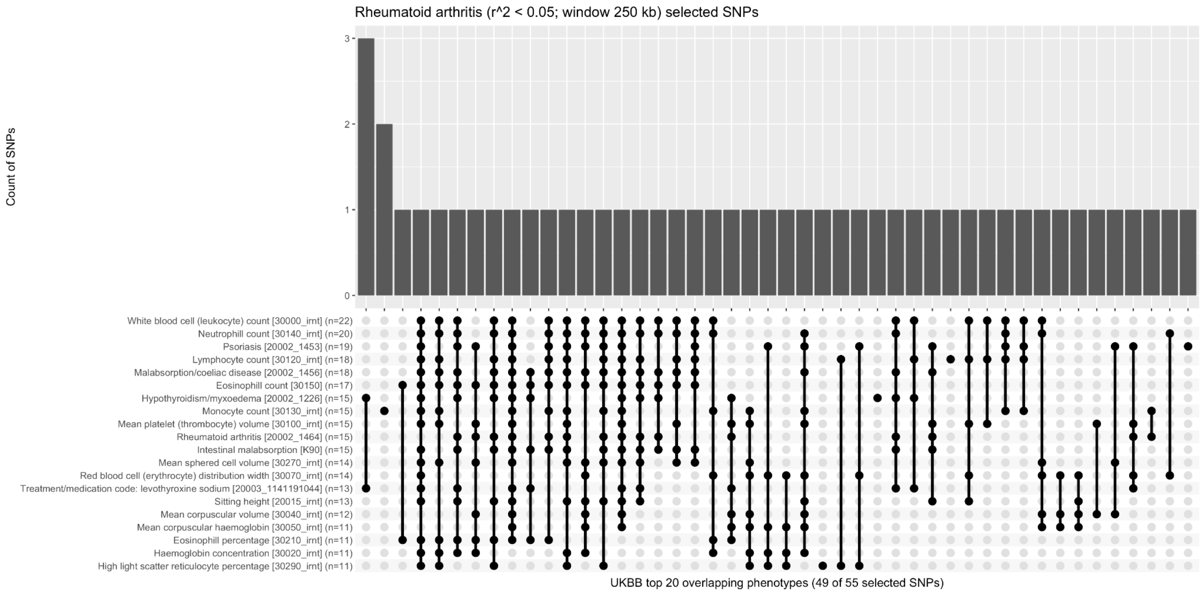


**Figure S6.29.** Upset plot showing a histogram of selected RA-associated SNPs which share a pleiotropic association with one or more marginal phenotypes in the UK Biobank. Top 20 traits shown. Of the 55 selected RA-associated SNPs, 49 (89%) are also associated with one or more of the top 20 genetically correlated phenotypes.

#### Pleiotropic FinnGen traits

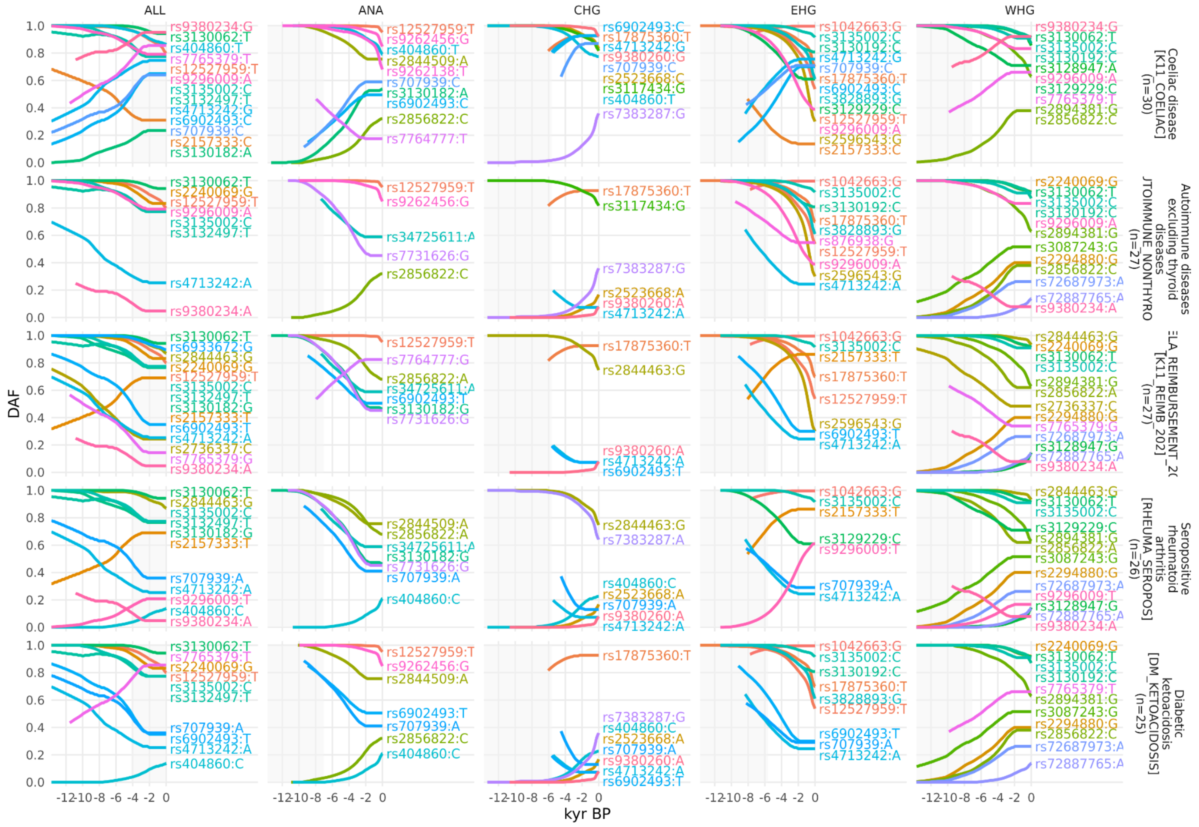

**Figure S6.30.** Allele frequency plots for positively selected RA-associated SNPs that are also associated with other phenotypes in the FinnGen study. Traits 1-5. SNPs are shown with their maximum likelihood trajectories, and polarised by the direction of their effect on the marginal FinnGen trait (i.e. showing the 'risk' allele). Phenotypes are ordered according to the number of common SNPs, portions of the trajectory with low posterior density are cropped off, and the background is shaded for the approximate time period in which the ancestry existed as an actual population.

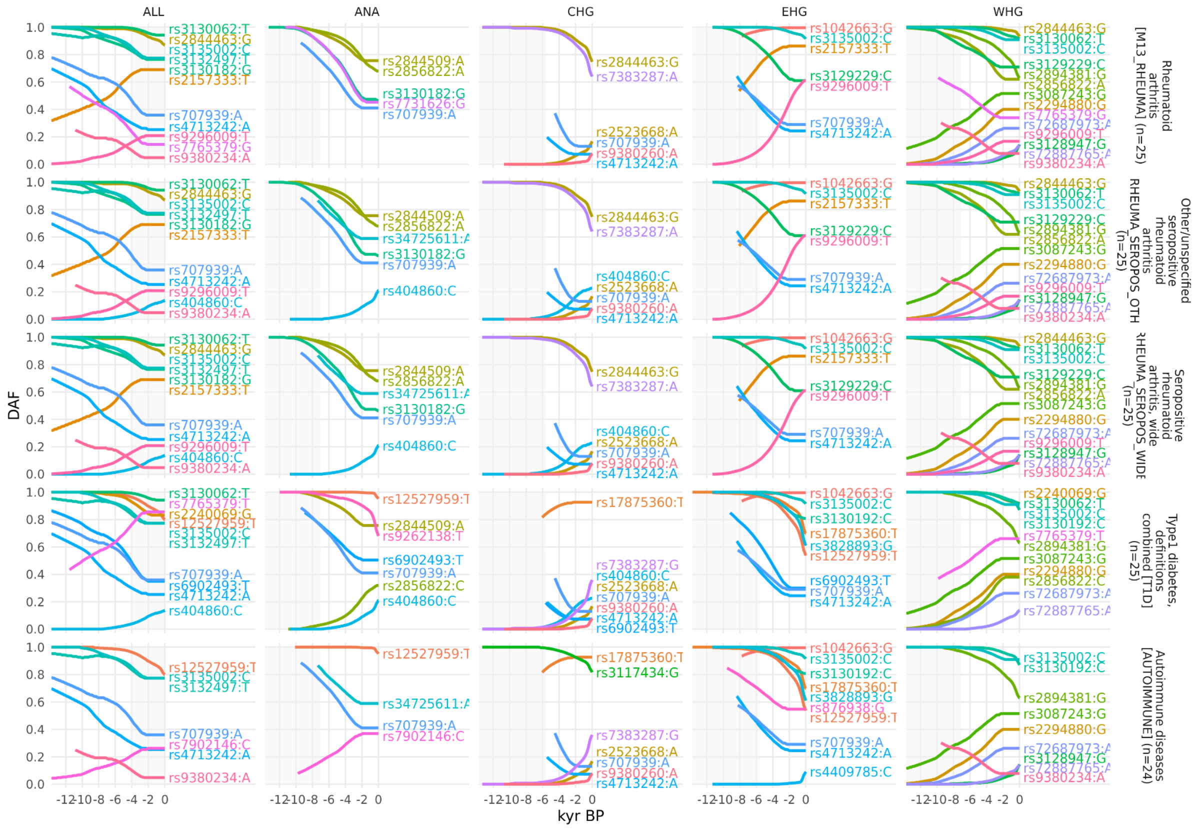


**Figure S6.31.** Allele frequency plots for positively selected RA-associated SNPs that are also associated with other phenotypes in the FinnGen study. Traits 6-10. SNPs are shown with their maximum likelihood trajectories, and polarised by the direction of their effect on the marginal FinnGen trait (i.e. showing the 'risk' allele). Phenotypes are ordered according to the number of common SNPs, portions of the trajectory with low posterior density are cropped off, and the background is shaded for the approximate time period in which the ancestry existed as an actual population.

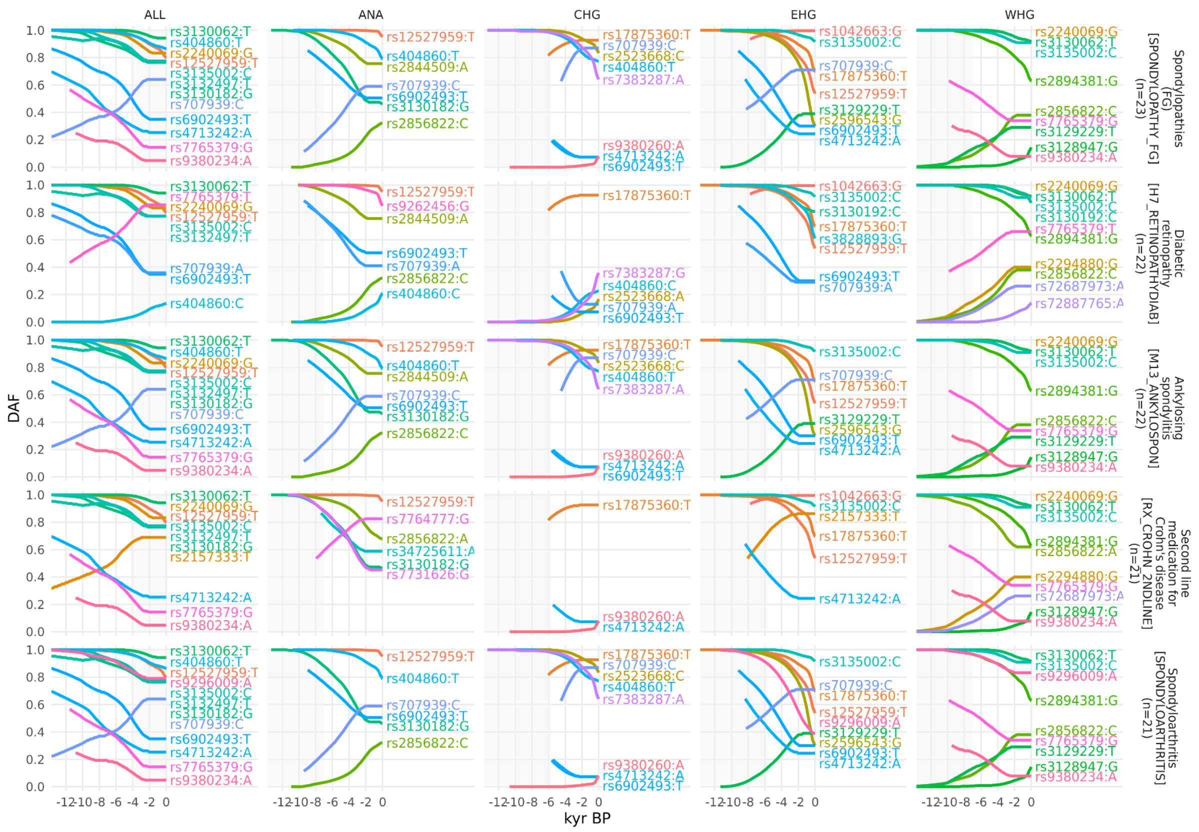


**Figure S6.32.** Allele frequency plots for positively selected RA-associated SNPs that are also associated with other phenotypes in the FinnGen study. Traits 11-15. SNPs are shown with their maximum likelihood trajectories, and polarised by the direction of their effect on the marginal FinnGen trait (i.e. showing the 'risk' allele). Phenotypes are ordered according to the number of common SNPs, portions of the trajectory with low posterior density are cropped off, and the background is shaded for the approximate time period in which the ancestry existed as an actual population.

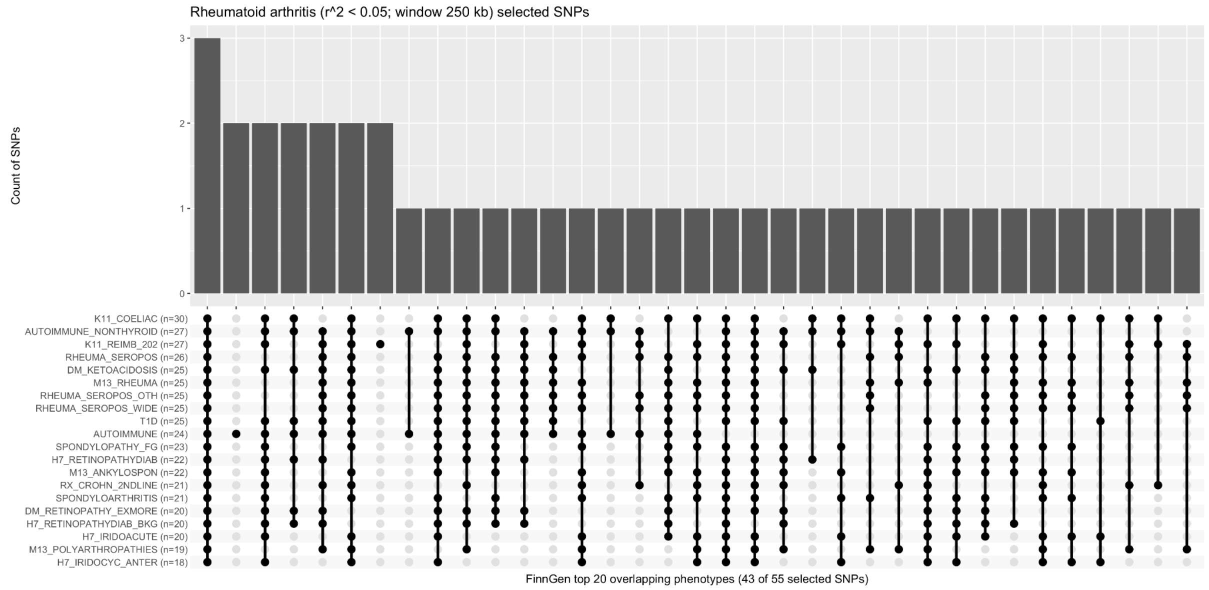


**Figure S6.33.** Upset plot showing a histogram of selected RA-associated SNPs which share a pleiotropic association with one or more marginal phenotypes in the FinnGen study. Top 20 traits shown. Of the 55 selected RA-associated SNPs, 43 (78%) are also associated with one or more of the top 20 genetically correlated phenotypes.

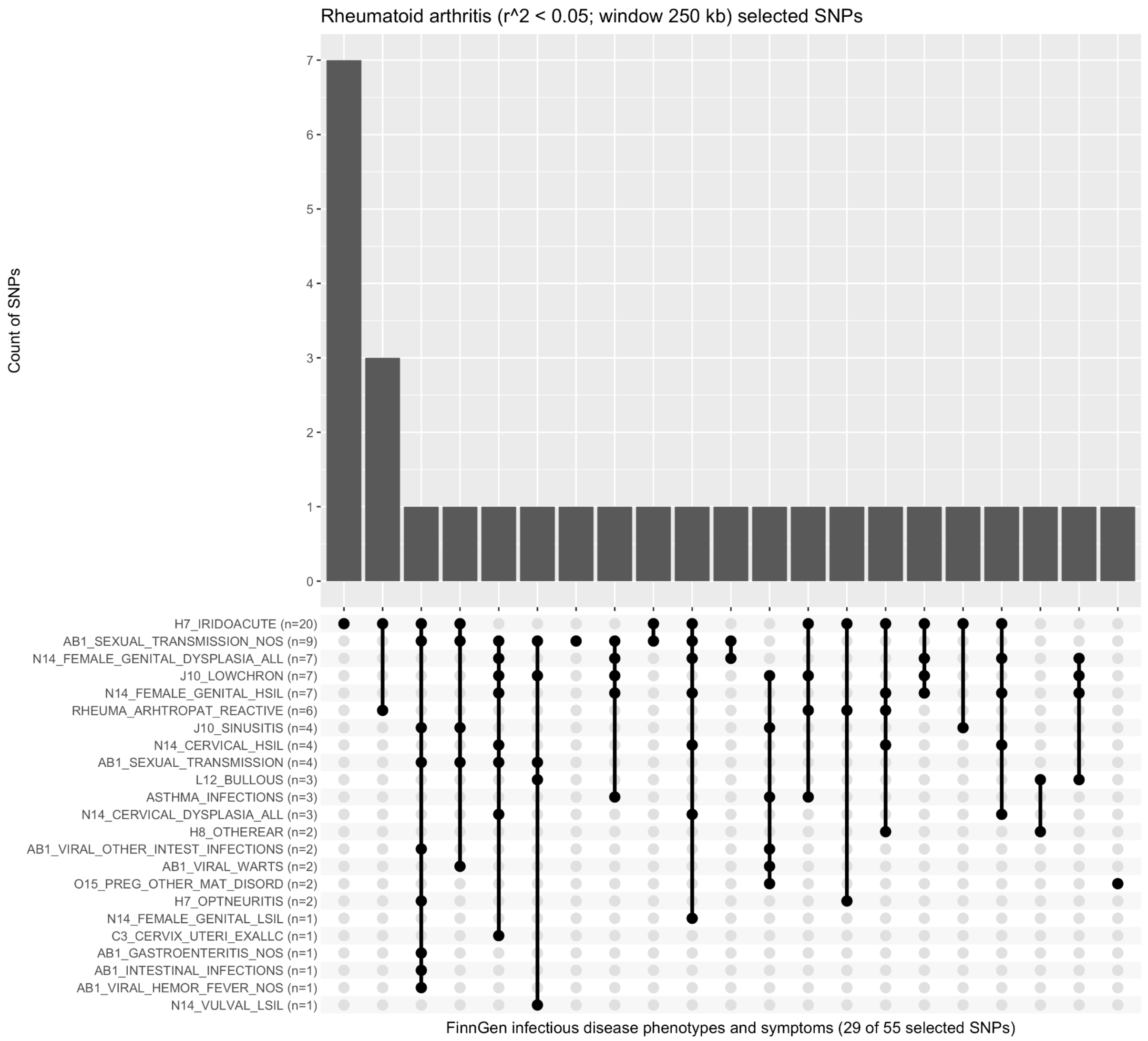


**Figure S6.34.** Upset plot showing a histogram of selected RA-associated SNPs which share a pleiotropic association with one or more infectious disease phenotypes and symptoms in the FinnGen study. Of the 55 selected RA-associated SNPs, 29 (53%) are also associated with one or more infectious disease phenotypes and symptoms.

### Discussion

To understand how natural selection has influenced the genetic component of auto-immune disease risk, we sought to model the allele frequency trajectories of risk associated variants through time, in a large panel of ancient genomes, and to test for evidence of polygenic selection acting on two auto-immune diseases: (i) multiple sclerosis (MS); and (ii) rheumatoid arthritis (RA).

We used two different callsets of trait associated variants for our two auto-immune diseases. The first callset contained fine-mapped variants; however, a large fraction of these variants were not callable in our imputed ancient dataset, due to quality control filtering and the difficulty of accurately inferring HLA alleles in ancient samples[31]. The resulting loss of so many high-effect HLA variants meant that our polygenic selection tests were statistically underpowered, and we did not identify any evidence of selection using this callset. To address this loss of statistical power, we ascertained a second callset of statistically independent and genome-wide significant markers, by LD-pruning the full set of genome-wide summary statistics for each auto-immune disease.

For multiple sclerosis, our our LD-pruned analyses identified statistically significant evidence of positive polygenic selection for trait associated variants in the ALL analysis ($p = 1.02e-5$; $\omega = 0.017$), and in the WHG ($p = 7.22e-5$; $\omega = 0.021$), EHG ($p = 2.60e-3$; $\omega = 0.016$) and CHG ($p = 3.06e-2$; $\omega = 0.009$) ancestral paths, but not in the ANA ($p = 6.43e-1$; $\omega = 0.004$) path. For rheumatoid arthritis, our analyses identified statistically significant evidence of negative polygenic selection for trait associated variants in the ALL analysis ($p = 3.26e-3$; $\omega = -0.007$) only, although the CHG ($p = 6.33e-2$; $\omega = -0.014$) path came close to reaching nominal significance. The WHG ($p = 8.63e-1$; $\omega = -0.001$), EHG ($p = 6.24e-1$; $\omega = -0.003$) and ANA ($p = 1.49e-1$; $\omega = -0.006$) ancestral paths showed no significant evidence of selection, although they all exhibited a negative selection gradient.

Our results also demonstrate that the strength of evidence for selection in a marginal selection test (i.e., the p-value for an individual SNP) is only partially correlated with the longitudinal effect of that variant on the trait itself (Figues S6.7. and S6.25). Our CLUES selection test is best powered to identify selection under a model in which the change in frequency is rapid. When changes in frequency occur gradually over a long period of time, it is harder to reject a neutral model of drift. Consequently, small but rapid changes in allele frequencies achieve low p-values, whilst large but slow changes achieve higher p-values. These small rapid changes may achieve statistical significance in a marginal test, whilst having very little longitudinal effect on the trait of interest. Conversely, large gradual changes with substantial longitudinal effect may not achieve statistical significance in a marginal test.

We caution that our results do not indicate that either MS or RA were directly under selection; and we are not suggesting that MS was evolutionary adaptive, or that ancient people suffered from either of these diseases. Rather, we observe that there is a very high degree of pleiotropy in auto-immune disease associated variants, consistent with the hypothesis that selection has favoured a strong immune response to exposure to environmental pathogens as a result of the transition to a pastoralist lifestyle, which may have driven the observed positive signals of polygenic selection for MS. We also note that RA has many overlapping SNPs with MS, and that the direction of effect for these SNPs is often opposing between the two traits.

Because MS would not have conferred a fitness advantage on ancient individuals, it is likely that this selection was driven by traits with shared genetic architecture, of which increased risk for MS in the present is a pleiotropic by-product. We therefore looked at LD-pruned MS-associated SNPs that showed statistically significant evidence for selection using CLUES (n=32) and which also had a genome-wide significant trait association ($p < 5e-8$) in any of the 4,359 traits from the UK Biobank[13,14] and 2,202 traits from the FinnGen study[15]. We observed that many selected SNPs were also associated with a variety of other traits, including type 1 diabetes (FinnGen, n=21), celiac disease (FinnGen, n=19; UKB, n=18), ankylosing spondylitis (FinnGen, n=17), white blood cell count (UKB, n=16), and lymphocyte count (n=16 UKB) (Figure S6.8 - S6.15). To determine if the observed signal of polygenic selection favouring MS risk could be better explained by selection acting on a genetically correlated trait, we performed a systematic analysis of traits in UK Biobank and FinnGen with at least 20% overlap among the MS-associated selected SNPs (n=115 traits). Using a joint test in PALM, specifically designed for disentangling polygenic selection on correlated traits, we did not identify any UK Biobank or FinnGen traits where the selection signal favouring MS risk was significantly attenuated by selection acting on a genetically correlated trait, when accounting for the number of tests (Figures S6.17 - S6.18). In the UK Biobank the trait which showed the strongest attenuating effect in the ALL ancestry path was mean platelet (thrombocyte) volume (30100_irnt) ($p$ = 1.06e-3), although it did not achieve nominal significance. In the FinnGen study, the trait which showed the strongest attenuating effect in the ALL ancestry path was psoriatic arthropathies (M13_PSORIARTH) ($p$ = 8.46e-3). These results show that the signal of polygenic selection favouring MS risk is at least partially independent of all other tested traits, consistent with our hypothesis that genetic risk for MS is the result of a complex evolutionary response to changes in exposure to multiple pathogens during the Bronze Age.

### 7) Ancestry linkage disequilibrium (LDA) and Ancestry linkage disequilibrium score (LDAS)

Yaoling Yang[1,2], Daniel Lawson[1,2]

[1]Department of Statistical Sciences, School of Mathematics, University of Bristol, UK
[2]Integrative Epidemiology Unit, Population Health Sciences, University of Bristol, UK

#### Methods

##### Definition

In population genetics, linkage disequilibrium (LD) is defined as the non-random association of alleles at different loci in a given population (Slatkin, 2008[1]). We propose an ancestry linkage disequilibrium (LDA) approach to measure the association of ancestries between SNPs.

Let $A(i, j, k)$ denote the probability of the $k$th ancestry ($k = 1,..., K$) at the $j$th SNP ($j = 1,..., J$) of a chromosome for the $i$th individual ($i = 1,..., N$).

We define the distance between SNP $l$ and $m$ as the average $L_2$ norm between ancestries at those SNPs. Specifically we compute the $L_2$ norm for the $i$th genome as

$$D_i(l, m) = ||A(i, l, \cdot) - A(i, m, \cdot)||_2 = \sqrt{\frac{1}{K}\sum_{k=1}^{K} (A(i, l, k) - A(i, m, k))^2}.$$

Then we compute the distance between SNP $l$ and $m$ by averaging $D_i(l, m)$:

$$D(l, m) = \frac{1}{N}\sum_{i=1}^{N} D_i(l, m).$$

We define $D^*(l, m)$ as the theoretical distance between SNP $l$ and $m$ if there were no linkage disequilibrium of ancestry (LDA) between them. $D^*(l, m)$ is estimated by

$$D^*(l, m) \approx \frac{1}{N}\sum_{i=1}^{N} ||A(i^*, l, \cdot) - A(i, m, \cdot)||_2,$$

where $i^* \in \{1,..., N\}$ are re-sampled without replacement at SNP $l$. Using the empirical distribution of ancestry probabilities accounts for variability in both the average ancestry and its distribution across

SNPs. Ancestry assignment can be very precise in regions of the genome where our reference panel matches our data, and uncertain in others where we only have distant relatives of the underlying populations.

The LDA between SNP $l$ and $m$ is a similarity, defined in terms of the negative distance $-\ D(l, m)$ normalized by the expected value $D^*(l, m)$ under no LD, as:

$$LDA(l, m)\ = \frac{D^*(l,m)-D(l,m)}{D^*(l,m)}.$$

LDA therefore takes an expected value 0 when haplotypes are randomly assigned at different SNPs, and positive values when the ancestries of haplotypes are correlated.

LDA is a pairwise quantity. To arrive at a per-SNP property, we define the LDA score (LDAS) of SNP $j$ as the total LDA of this SNP with the rest of the genome, i.e. the integral of the LDA for that SNP. Because this quantity decreases to zero as we move away from the target SNP, this is in practice computed within an $X$cM-window (we use $X\ =\ 5$ as LDA is approximately zero outside this region in our data) on both sides of the SNP. Note that we measure this quantity in terms of the genetic distance, and therefore LDAS is measuring the length of ancestry-specific haplotypes compared to individual-level recombination rates.

As a technical note, when the SNPs approach either end of the chromosome, they no longer have a complete window, which results in a smaller LDAS. This would be appropriate for measuring total ancestry correlations, but to make LDAS useful for detecting anomalous SNPs, we use the LDAS of the symmetric side of the SNP to estimate the LDAS within the non-existent window.

$$LDAS(j;X) = \begin{cases} \int_{gd(j)-X}^{gd(j)+X} LDA(j,l)\,dgd & \text{if } X \le gd(j) \le tg - X, \\ \int_{0}^{gd(j)+X} LDA(j,l)\,dgd + \int_{2gd(j)}^{gd(j)+X} LDA(j,l)\,dgd & \text{if } gd(j) < X, \\ \int_{gd(j)-X}^{tg} LDA(j,l)\,dgd + \int_{gd(j)-X}^{2gd(j)-tg} LDA(j,l)\,dgd & \text{if } gd(j) > tg - X. \end{cases}$$

where $gd(l)$ is the genetic distance (i.e. position in cM) of SNP $l$, and $tg$ is the total genetic distance of a chromosome. We also assume the LDA on either end of the chromosome equals the LDA of the SNP closest to the end: $LDA(j, gd\ =\ 0)\ =\ LDA(j, l_{min(gd)})$ and $LDA(j, gd\ =\ td)\ =\ LDA(j, l_{max(gd)})$ , where $gd$ is the genetic distance, $l_{min(gd)}$ and $l_{max(gd)}$ are the indexes of the SNP with the smallest and largest genetic distance, respectively.

The integral $\int_{gd(j)-X}^{gd(j)+X} LDA(j, l)dgd$ is computed assuming linear interpolation of the LDA score between adjacent SNPs.

LDA thus quantifies the correlations between the ancestry of two SNPs, measuring the proportion of individuals who have experienced a recombination leading to a change in ancestry, relative to the genome-wide baseline. The LDA score is the total amount of genome in LDA with each SNP (measured in recombination map distance).

#### Simulation for selection: LDA

An ancient population $P_0$ evolved for 2200 generations before splitting into two sub-populations $P_1$ (Steppe) and $P_2$ (Farmer). After evolving 400 generations, we added mutation "$m_1$" and "$m_2$" at the different locus in $P_1$ and $P_2$. Both added mutations were then positively selected in the following 300 generations, after which they merged to $P_3$, where both added mutations experienced strong positive selection for 20 generations. Finally, we sampled 1000 individuals from $P_3$ to compute their ancestry proportions of $P_1$ and $P_2$ using the "chromosome painting" technique, and calculated the LDA score of the simulated chromosome positions.

The above describes the simulation in Figure S7.1.

We investigated balancing selection at 2 loci as well. The balancing selection in $P_1$ and $P_2$ ensured the mutated allele reaches around 50% frequency, while positive selection made the mutated allele become almost the only allele. In $P_3$, if $m_1$ or $m_2$ was positively selected, its frequency reached more than 80% regardless of whether the allele experienced balancing or positive selection in $P_1$ or $P_2$, because we set a strong positive selection. If $m_1$ or $m_2$ was balancing selected in $P_3$, its frequency slightly increased, e.g. if $m_1$ underwent balancing selection in $P_1$, it had 25% frequency when $P_3$ was created, and the frequency reached around 37.5% after 20 generations of balancing selection in $P_3$.

The results (Figure S7.2) show that positive selection in $P_3$ resulted in low LDA scores around the selected locus, if this allele was not uncommon (i.e. had 50% (balancing selection) or 100% frequency (positive selection) in subpopulation $P_1$ or $P_2$). Note that the balancing selection in $P_1$ or $P_2$ worked the same as “weak positive selection”, because $m_1$ and $m_2$ were rare when they first occured, which were positively selected until 50% frequency.

We also performed simulations for selection at a single locus (Figure S7.2 & Figure S7.3).

Stage 1: We added a mutation $m_1$ in the 1600 generation in $P_0$, which then underwent balancing selection until generation 2200, when $P_0$ split into $P_1$ and $P_2$, where the frequency of $m_1$ was around 50%.

Stage 2: Then we explored different combinations of positive, balancing and negative selection of $m_1$ in $P_1$ and $P_2$. the frequency of $m_1$ reached 80%, 50% and 20% when it was positively, balancing or negatively selected, respectively, until generation 2899. Here we sampled 20 individuals each in $P_1$ and $P_2$ as the ancient samples.

Stage 3: Then $P_1$ and $P_2$ merged into $P_3$ in generation 2900. In $P_3$, for each combination of selection in Stage 2, we simulated positive, balancing and negative selection for $m_1$. The selection lasted for 20 generations, and then we sampled 4000 individuals from $P_3$ as the modern population.

Results: when $m_1$ was positively selected in only one of $P_1$ and $P_2$, and it experienced negative selection in $P_3$, the LDA scores around the locus of $m_1$ were low. Otherwise, no abnormal LDA scores were found at $m_1$.

### Results

#### Simulation for LDA scores with selection in one or two loci

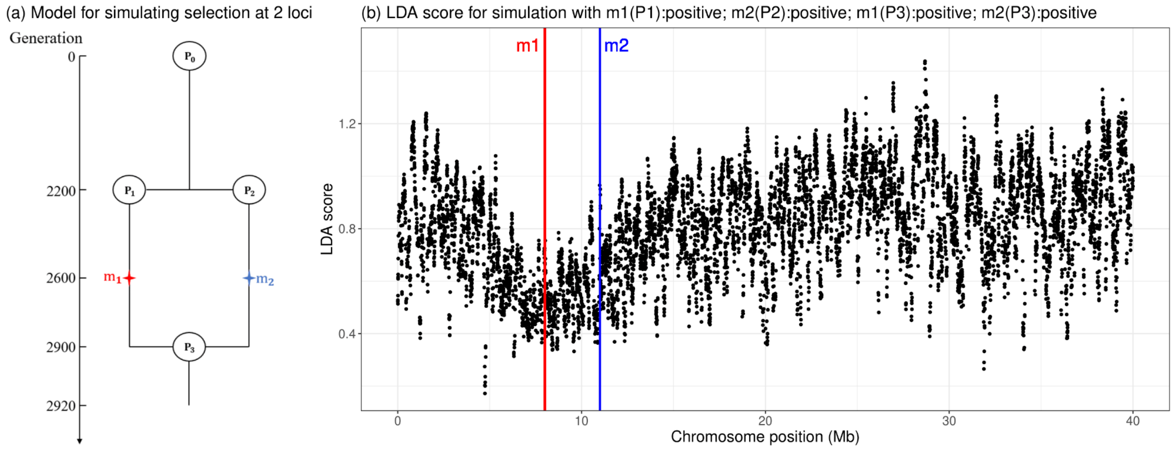


**Figure S7.1: Simulating Low LDA score**.

Left: A simulated history in which a single population splits into two ("Steppe" and "Farmer") after 2200 generations and experiences positive selection on different loci ($m_1$ in $P_1$ and $m_2$ in $P_2$). After 2900 generations the populations merge ("Europeans") but selection continues on *both* loci.

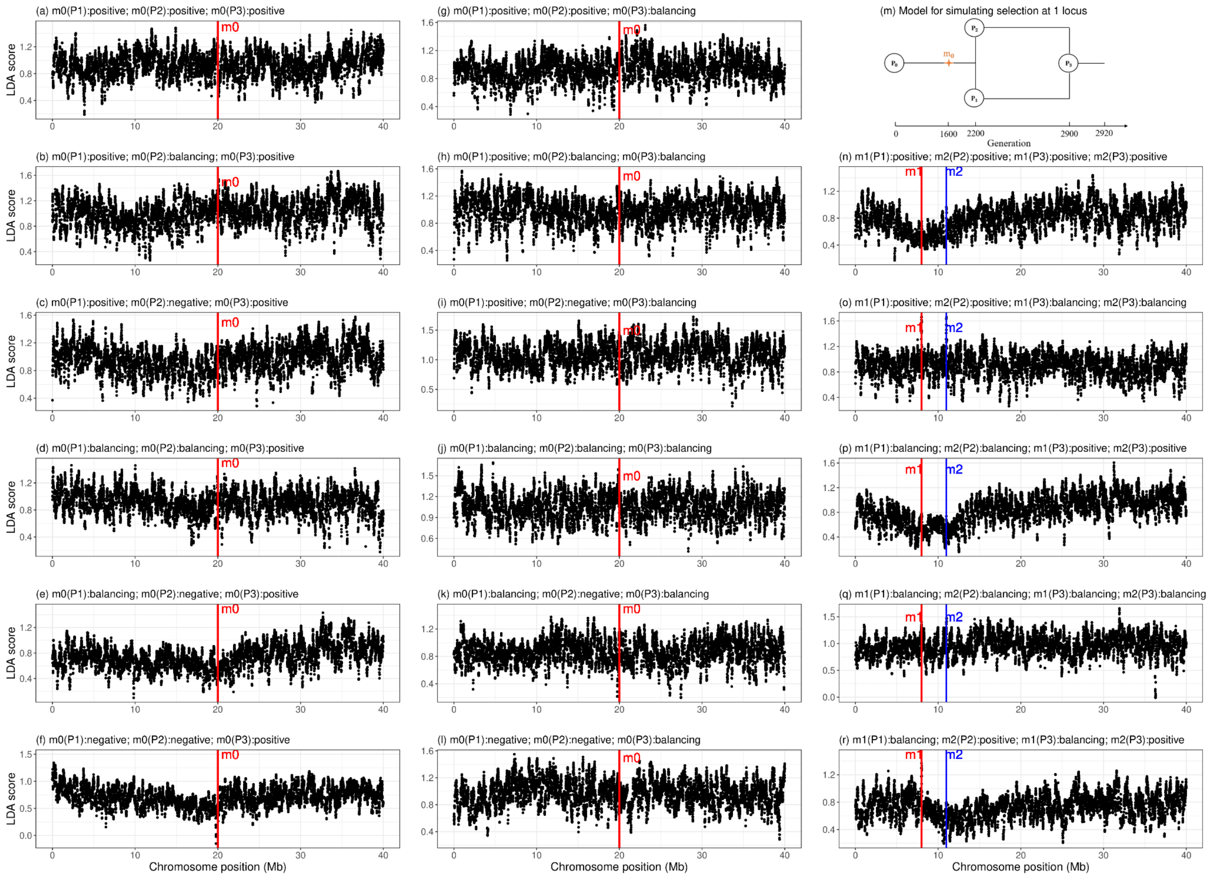


**Figure S7.2: LDAS simulation with positive or balancing selection in the modern population.** The left two columns show simulations with a single variant satisfying the observed constraint that modern-day frequencies are not decreasing (i.e. not negative selection). The right column shows simulations with two variants, also obeying this constraint. The model for simulating 2 loci is the same as in Figure S7.1, and that for 1 locus is in the top right of this plot (which differs only in the location of the selected variant in the separated populations).

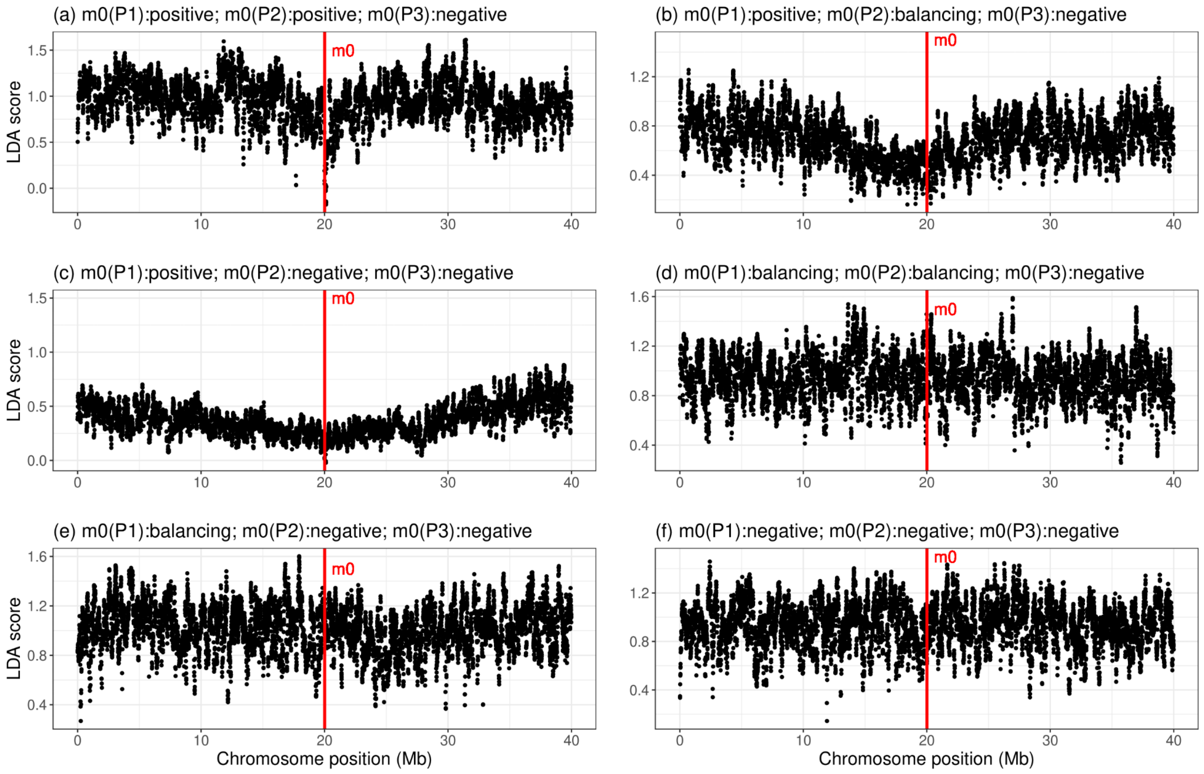


**Figure S7.3: LDAS simulation with single locus negatively selected in the modern population.** In two cases this generates a low LDAS score, which requires recent negative selection (which is ruled out for HLA by the observed frequency trend). The model used is in the top right of Figure S7.2.

#### LDA score for chromosome 2 and 6

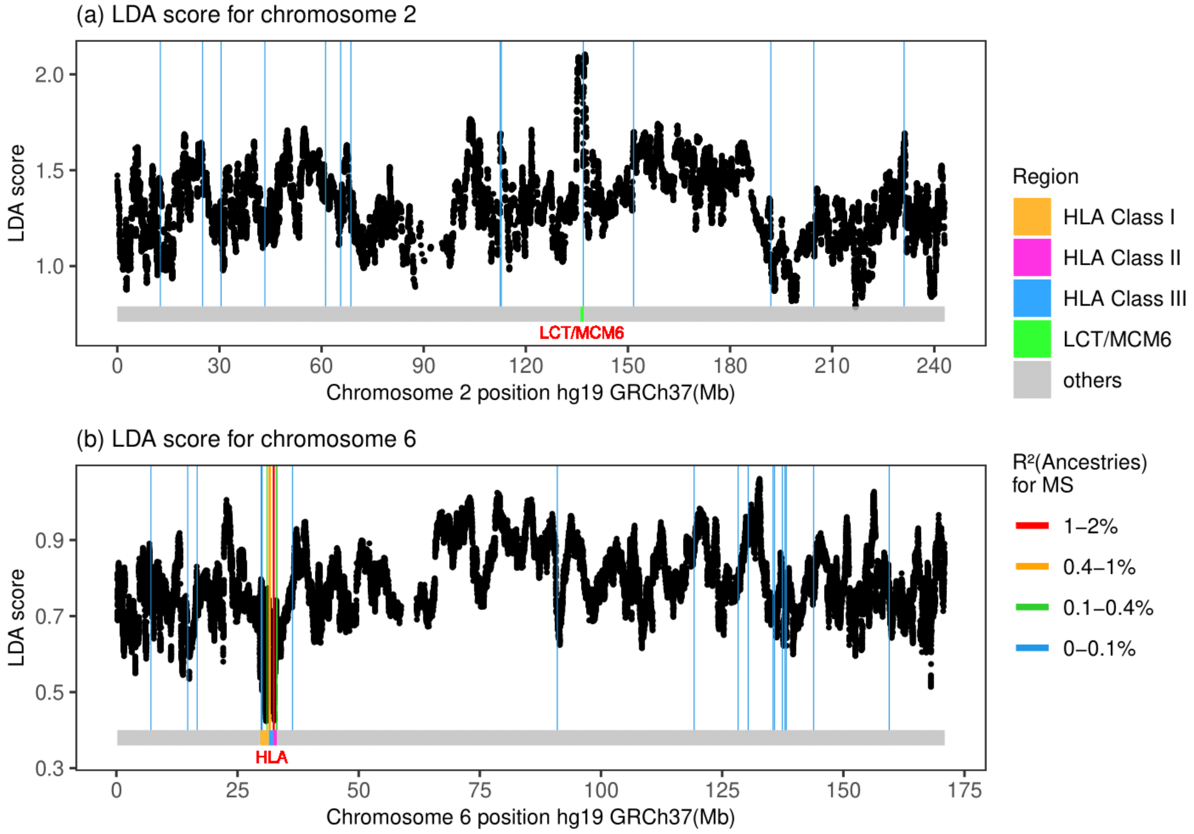


**Figure S7.4: LDAS on chromosome 6 and 2.** LDA score is a) high in the LCT/MCM6 region while is b) low in the HLA region.

#### LD plot for chromosome 6 MS-associated SNPs

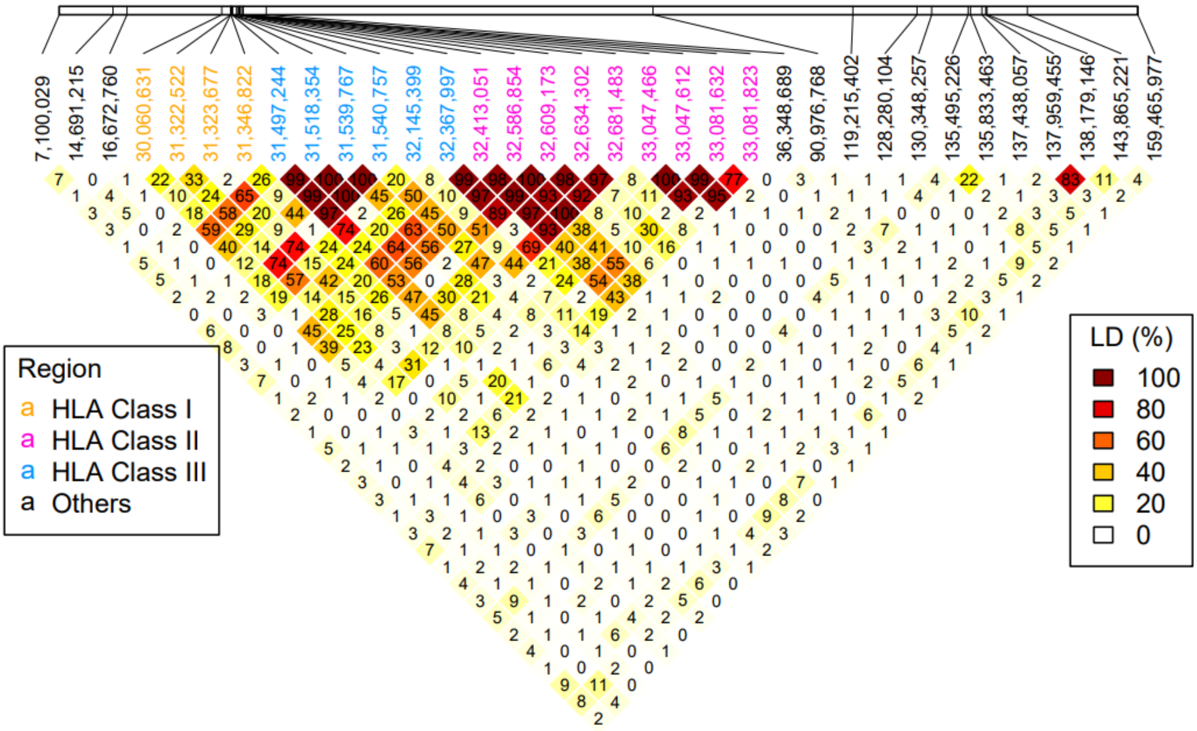


**Figure S7.5: Pairwise Linkage Disequilibrium (LD) plot (measured by D') for all the MS-associated SNPs on chromosome 6.**

### 8) Pleiotropic effects of selected SNPs associated with multiple sclerosis or rheumatoid arthritis

Astrid K.N. Iversen[1], William F.S. Barrie[2], Evan K. Irving-Pease[3]

[1]Nuffield Department of Clinical Neurosciences, Division of Clinical Neurology, Weatherall Institute of Molecular Medicine, University of Oxford, John Radcliffe Hospital, Oxford, United Kingdom

[2]Zoology Department, University of Cambridge, UK.

#### Introduction

Autoimmune disease susceptibility is unlikely to be selected due to any beneficial effect on survival. We propose that an increase in the risk of autoimmune disease is more likely a byproduct of selection for gene variants associated with, e.g., advantageous immune responses to the pathogenic challenges ancient populations faced.

The current incidence of RA (Figure S8.1) is approximately 100 times higher than that of MS. Although people commonly develop RA between the age of 35 to 60, it can occur at any age. The average age of early-onset RA is 14. MS can develop at any age, but onset usually occurs between 20-40. Women are more frequently affected by both RA and MS. Studies of ancient skeletons suggest that RA has affected people for thousands of years but the prevalence over time is unknown (Entezami et al., 2011). MS leaves no unique marks on the skeleton, so we cannot know how frequently MS developed in ancient times; the first MS patient was described in 1859.

RA and MS are autoimmune diseases with genetic and environmental risk factors. While several pathogens are thought to play a part in MS disease development, EBV has been identified as a major risk factor for MS, especially if infection occurs after 18 years of age (Bjornevik et al., 2022). In contrast, although several bacteria and respiratory viruses have been implicated as potential RA disease triggers, no single pathogen stands out (Joo et al., 2019). Multiple RA studies suggest that mucosal surfaces, particularly the periodontal region, lung and gut, may be sites of inflammation and potential generation of RA-related autoimmunity in the preclinical period of the disease (Deane et al., 2017). Whether the pathogens that trigger MS and RA today were present to the same extent in ancient times is unknown.

HLA-DRB1*15:01 has been associated with some protection against tuberculosis (Tervi et al., 2022), while HLA-DRB1*04:01 has been suggested to be associated with better outcomes in, e.g., hepatitis C virus (HCV) (Hydes et al., 2015), Hepatitis B virus (HBV) (Yan et al., 2012), and SARS-CoV-2 infections (Langton et al., 2021) in people of European descent, and with strong immune responses to influenza peptides (Danke & Kwok, 2003).

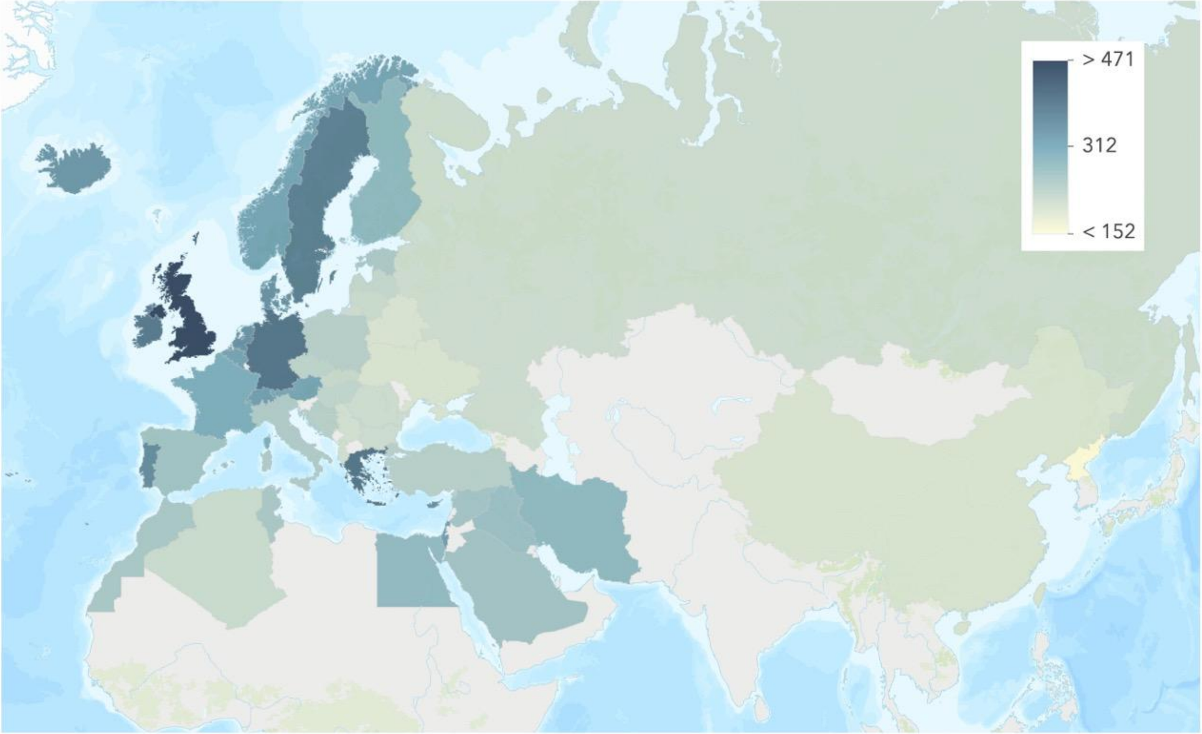


**Figure S8.1. Modern prevalences of RA.**
Modern-day geographical distribution of RA prevalence in Europe. Prevalence data for RA (cases per 100,000) was obtained from Safiri et al. (2019).

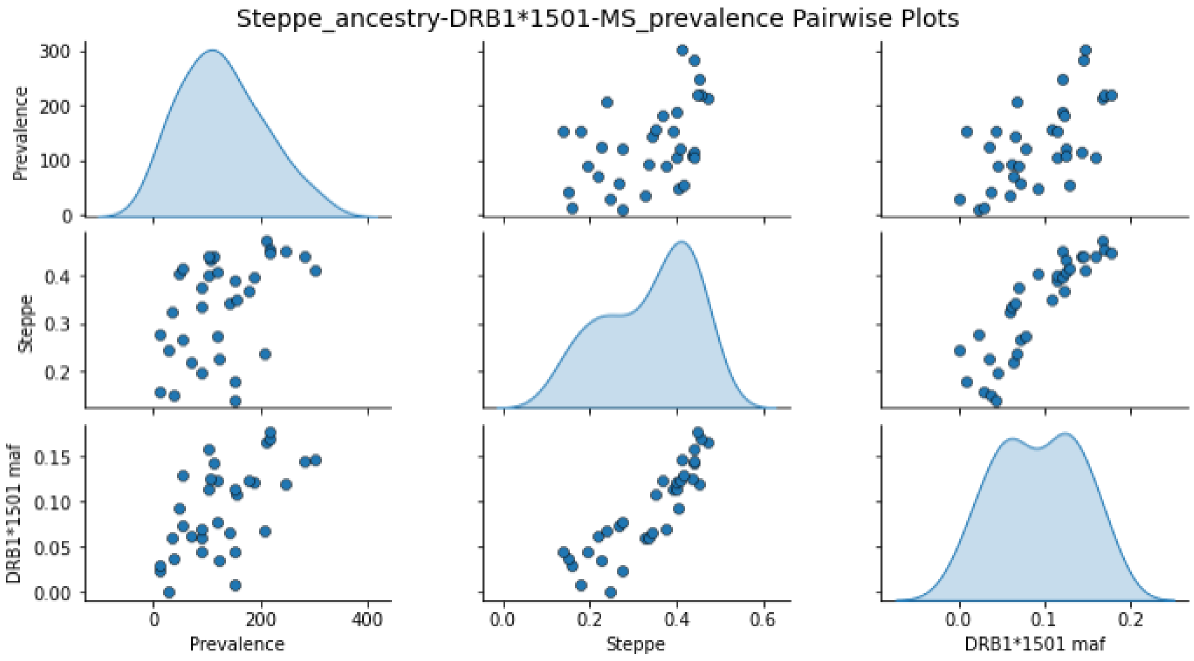


**Figure S8.2. Association between genome-wide Steppe ancestry, MS prevalence and DRB1*15:01 frequency in modern populations in the UK Biobank.**

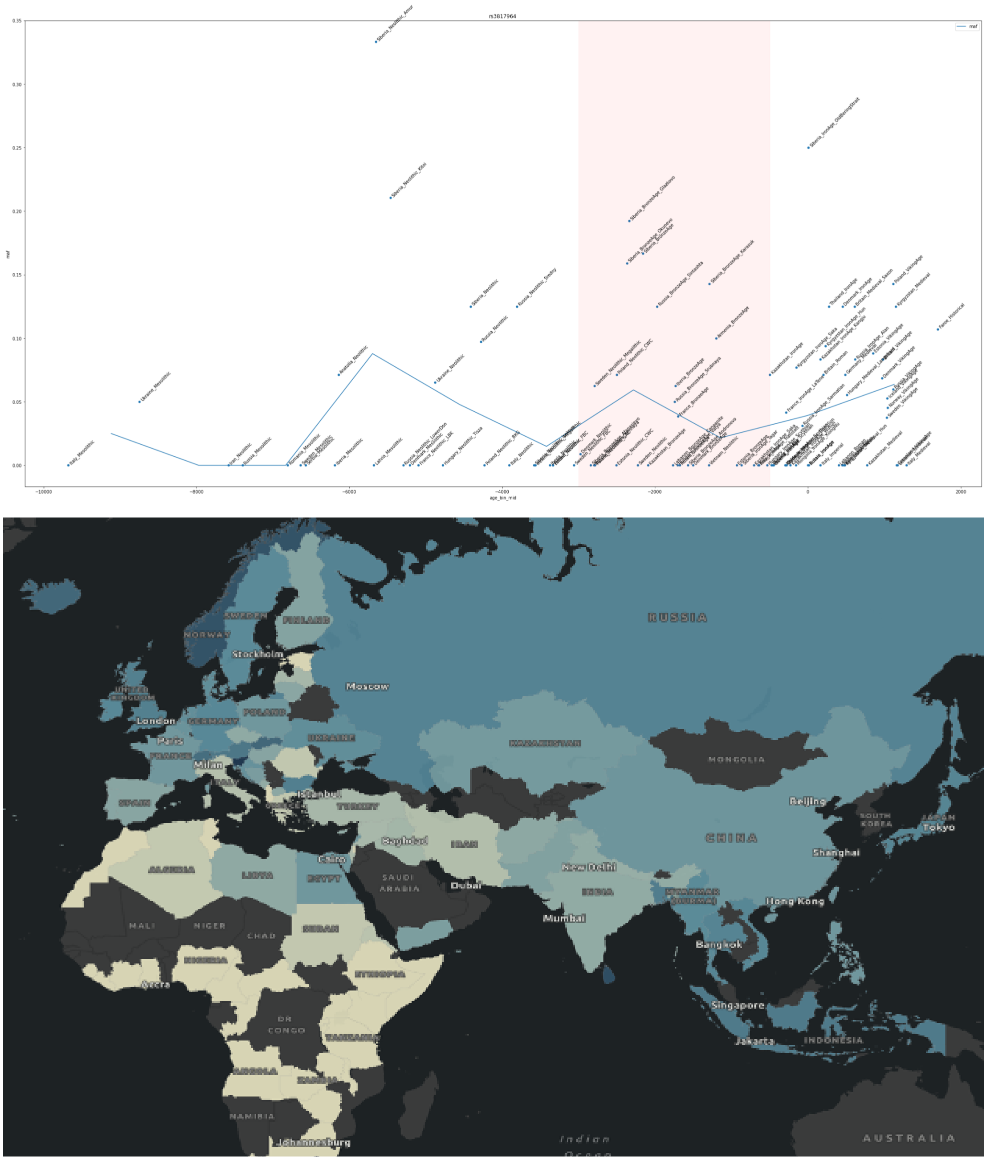


**Figure S8.3. Ancient and modern prevalences of HLA-DRB1*04:01 (rs3817964).**

Top: Ancient distributions of HLA-DRB1*04:01, the largest genetic risk factor in RA. Average frequency across all populations is shown (blue line, 10 time bins) as well as the Bronze Age (red shading).

Bottom: Modern distribution of HLA-DRB1*04:01 in populations in the UK Biobank. NB the tag SNPs may be less effective at tagging these types in non-European populations, so we urge caution in interpretation - especially in African populations.

### Methods

#### Pleiotropic trait analysis

To investigate possible selective pressures driving these signals, we analysed the pleiotropic pathogen- and/or infectious disease-associated effects of alleles found to have been under positive selection (Supplementary Note 6) in each of the four ancestry paths and the pan-ancestry path by systematically searching for associations in existing literature in PubMed, the UK Biobank, and the Finngen Biobank.

We observed three main types of protective links: (i) to distinct pathogens, e.g., herpes simplex virus (HSV), or their associated diseases, e.g., chickenpox, which is caused by varicella zoster virus (VZV), (ii) to diseases affecting specific systems, e.g., respiratory infections, or, (iii) to distinct diseases caused by nonspecific parasites, bacteria, fungi or viruses in, e.g., the skin and subcutis. These SNP associations are shown in ST11 and ST12.

Our analyses had several limitations. We cannot identify alleles associated with protection against pathogens known to have exerted strong selective pressures on human populations throughout history for which no GWAS data exists, e.g., plague and smallpox. Neither is robust GWAS data available that identify alleles associated with protection against pathogens which we currently vaccinate most of the populations against, e.g., mumps, rubella, measles, H. influenza and polio. Moreover, some studies are small, and some SNP associations might differ between ethnicities (for example due to different LD patterns between populations). As a result, many of these associations are suggestive rather than conclusive and may be limited to European populations.

### Results

#### Pleiotropic trait analysis of SNPs associated with multiple sclerosis

We identified 32 selected MS SNPs (ST8, ST11); of these, six (6/32, 19%) were not associated with protection against any specific pathogen or infectious disease. Two of these seven selected SNPs were associated with positive selection of the risk allele for MS. In many of these cases, SNPs were associated with, e.g., decreased risk of diabetes type I and coeliac disease (rs3134931; OR 0.750 and 0.818, respectively), coeliac disease and malabsorption (rs3806156; OR 0.658 and 0.672, respectively), diabetes and coeliac disease (rs9368699; OR 0.339 and 0.308, respectively), which suggests the presence of potentially strong, non-infectious selective forces.

We observed shared protective alleles when considering Epstein-Barr Virus (EBV). 1/32 positively selected SNPs (~3%) were associated with a lower risk of EBV-associated disease: this conferred protection to both EBV and HSV in ANA (rs2735076; HLA-A).

Overall, 4/32 SNPs (~13%) were linked to protection against cellulitis and/or abscesses in different body parts, usually encompassing arms, hands, legs, feet and toes (ST11) (rs2735076, HLA-A, ANA path; rs9261588, TRIM26, WHG and pan-ancestral paths; rs1077667, TNFSF14, ANA path; rs1794275, ANA and WHG but in opposite directions). For each SNP there were additional links to protection against different pathogens and/or diseases such as herpes simplex virus (HSV), keratitis and keratoconjunctivitis (rs1794275),  HSV and EBV (rs2735076), and retrovirus infection (rs9261588).

This analysis revealed two protective alleles against the mumps virus (MuV) (rs2070600, AGER, WHG and EHG paths; rs241447, HLA-DOB, WHG and EHG paths). Both SNPs were also associated with protection against a range of other pathogens.

Four of the 32 SNPs (~13%) were associated with protection against chickenpox or shingles (caused by varicella zoster virus (VZV)) and each were found in unique genes. One SNP was selected exclusively in the CHG path (rs7747873, HLA-E), one in the EHG, WHG and CHG paths (2735046, HLA-F), one in the WHG, and pan-ancestral paths (rs17576984, PGBD1), and one in the CHG path (rs9266773, MICA). Whereas no other protective effects were associated with rs17576984, the other four SNPs were associated with protection against other non-specific or specific pathogens or diseases, e.g., coronavirus, viral infections, pyogenic arthritis, sexually transmitted diseases (rs9266773, MICA).

One SNP was associated with protection against influenza or influenza and pneumonia: rs210131 was exclusively selected in the EHG and pan-ancestral paths (rs210131, BAK1).

Two SNPs were associated with protection against streptococcus pneumonia: one in WHG (rs9296015, NOTCH4), and one in WHG and the pan-ancestry paths (rs3129934, TSBP1). Several SNPs were associated with protection against non-specific viral or bacterial pneumonia (ST11).

Three SNPs were associated with protection against non-specific or specific parasites, and some had additional protective effects against viruses and/or bacteria or broadly against infection regardless of associated pathogen. One parasite-associated protective SNP was observed in the WHG and EHG paths (rs241447, HLA-DOB), one in the WHG path (rs7200786, CLEC16A; specifically protective against helminths), and one SNP was shared between the ANA and EHG paths (rs757260, TRIM40).

We found four selected SNPs were linked to protection against gastrointestinal infections; two of these were specifically linked to protection against clostridium difficile in CHG and the pan-ancestry paths (rs416622, HLA-DOA) and in the ANA path (rs1077394, APOM). Moreover, one SNP was protective against bacterial enteritis in CHG and WHG (rs9268530, BTNL2) and one against intestinal infections in the WHG and EHG paths (rs241447, HLA-DOB). One SNP was protective against spirochetes in WHG (rs2248372, HCP5, MICB); no protective SNPs were found against spirochetes in the RA analysis.

One MS-risk SNP was associated with protection against cytomegalovirus (CMV) infection (rs12210887, NEU1). This protective allele was selected in the WHG and pan-ancestry paths.

We found several SNPs linked to protection against pneumonia caused by streptococcus (n=2, rs9296015, NOTCH4, WHG and rs3129934, TSBP1, WHG and pan-ancestral paths). One SNP was associated with protection against infection with mycobacteria, including Mycobacterium tuberculosis in the WHG path (rs4902647, ZFP36L1). Overall, most of the associated SNPs allowed for protection against pathogens/diseases affecting the respiratory system, the gastrointestinal tract, and genitalia. A complete list of SNP associations is available in ST11 and are outlined in Extended Data Figure 9.1.

#### Pleiotropic trait analysis of SNPs associated with rheumatoid arthritis

We identified 55 selected RA SNPs (ST9, ST12); 17 (31%) were not associated with protection against any specific pathogen or infectious disease (ST12). Seven of these 17 selected SNPs were associated with positive selection of the risk allele for RA in EHG, WHG, ANA, or CHG. SNPs were associated with e.g., a reduced risk of thyrotoxicosis (OR 0.889), intestinal malabsorption (OR 0.722), and celiac disease (OR 0.691) (rs2294880, BTNL2, WHG), or a lower risk of malabsorption (OR 0.608, rs9380260, MICB, CHG) and celiac disease (OR 0.628, rs9380260, MICB, CHG). In the other 12 cases, the selected variant alleles decreased the risk of RA. As RA could also develop in ancient times (Entezami et al., 2011), the disease itself and the impaired ability to combat other infections caused by RA-associated inflammation (Listing et al., 2013), would likely impose a negative selective pressure on the risk alleles as it would negatively impact the affected person's chances of survival.

In the RA analysis, two protective alleles against HSV were selected in the WHG path (rs72887765, RGL2), and the ANA, CHG, EHG, WHG and the pan-ancestral paths (rs9927316, LINC02132).

One RA-associated SNP was linked to a lower risk of EBV infection (1/55 SNPs, 2%) resulting in infectious mononucleosis (rs3130062, ATP6V1G2), and positive selection was only identified in the WHG and pan-ancestry paths.

The RA selection analysis included one protective allele against MuV (rs404860, NOTCH4) in the ANA, CHG and the pan-ancestry paths. This SNP was different from the protective SNPs found in the WHG and EHG paths in the MS selection analysis.

Six RA-risk SNPs were associated with protection against cellulitis and abscesses and were also linked to a reduction of the risk of several other pathogens and diseases. One SNP was linked to broad protection against bacterial and parasitic diseases and was found in the EHG and WHG paths (rs3130192, HLA-DPA1). One SNP was linked to a decreased risk of dermatophytosis and erysipelas and was selected in the WHG path (rs3087243, CTLA4). Postdysenteric arthropathy typically manifests after a bacterial gastrointestinal infection. Two SNPs were selected in either the WHG and pan-ancestry paths (rs2240069, TRIM31), or only in ANA (rs1953126, PHF19); these also decreased the risk of viral hepatitis in pregnancy and during the puerperium (rs2240069, TRIM31) or bronchopneumonia and lung abscesses (rs1953126, PHF19). Of the two SNPs selected in the WHG path, one also protected against meningitis and other infections caused by Neisseria meningitidis (rs8075737, NEUROD2). One SNP was selected in the ANA and pan-ancestry paths and protected against sexually transmitted diseases and optic neuritis (rs3130182; HLA-DPB1) and one SNP selected in WHG decreased the risk of puerperal infections, i.e infections during childbirth and during the subsequent 6 weeks thereafter (rs2301888, PADI6).

We observed one selected SNPs associated with protection against measles. This was selected in EHG, CHG, ANA and the pan-ancestry analysis (rs707939, MSH5).

Five SNPs were associated with protection against VZV-associated disease (9%). Of these SNPs, one was only associated with protection against VZV and was observed in the CHG path (rs2523668, MICB), while the rest also have been associated with protection against other pathogens/diseases. One was selected in the CHG, EHG and pan-ancestry paths (rs4713242, HLA-F), and was broadly protective against several viruses (e.g., viral haemorrhagic fevers and coronavirus). One SNP was found in the EHG path (rs1265096, PSORS1C2), and one in the CHG, EHG, ANA and pan-ancestry paths (rs6902493, CYP21A2). None of the VZV-protective SNPs in the MS and the RA analyses overlapped.

Two SNPs were protective against tuberculosis; one was only selected in the ANA path (rs7764777, HLA-G) and one was selected in the EHG, CHG, ANA and the pan-ancestry paths (rs2596543, MICA). This latter SNP is also associated with protection against candida infections. One SNP was associated with protection against pneumococcal pneumonia and was shared between the ANA and WHG paths (rs2856822, HLA-DPA1). One SNP was associated with protection against asthma-related pneumonia, and was selected in the ANA path (rs9262138, DHX16). One SNP was associated with protection against asthma-related acute respiratory infections in ANA, CHG and the pan-ancestry paths (rs404860, NOTCH4).

Five SNPs were associated with protection against parasites; all of these had additional protective effects against bacteria and possibly viruses and fungi (ST12). Two were also associated with protection against some sexually transmitted diseases, including one also protecting against influenza. All but one of the parasite-protective alleles were exclusively selected in one population path: EHG (rs876938, PRXL2B and rs1042663, CF8), and WHG (rs3128947, HLA-DOA and rs2894381, HLA-DQ2). The one shared SNP was selected in the EHG and WHG paths (rs3130192, HLA-DPA1). Two SNPs were linked to protection against viral hepatitis; the first SNP was associated with protection against hepatitis A, B and C virus and was selected in CHG (rs7383287, HLA-DOB). This was also associated with protection against clostridium difficile and the latter against a vast range of pathogens. The second was associated with protection against other forms of viral hepatitis (rs2240069, TRIM31) and was selected in the WHG and pan-ancestry paths. One SNP was associated with protection against arthropod-borne and other viral haemorrhagic fevers and was selected in the EHG, CHG and pan-ancestry paths (rs4713242, HLA-F).

We found one selected SNP was linked to protection against gastrointestinal infections; this was specifically linked to protection against clostridium difficile in CHG (rs7383287, HLA-DOB), and also offered protection against several other pathogens.

Overall, we found that 38 of the 55 (69%) RA-risk SNPs were associated with specific pathogens and/or diseases. Of these 38 SNPs, many were associated with pathogens or diseases linked to mucosal surfaces in the mouth, respiratory tract, or with pathogens which first replicate in the respiratory tract (e.g., measles (Lin et al., 2021), mumps (Katoh et al., 2015), EBV (Egan et al., 1995), VZV (Zerboni et al., 2014)), to gastrointestinal infections caused by parasites, bacteria or viruses, to urinary tract infections, or to infections in the genital tract. A complete list of SNP associations is available in ST12 and are outlined in Extended Data Figure 9.1.

#### Discussion

The SNPs associated with the risk of MS or RA under selection were often linked to protection against the same pathogens or diseases, e.g. EBV, HSV, VZV, mumps, acute respiratory infections, tuberculosis, viral hepatitis, cellulitis and abscesses and sexually transmitted diseases/genital infections. Some protective alleles were linked to a single pathogen or disease while others were associated with several pathogens and/or diseases. The observation that the MS and RA risk SNPs in most cases were linked to the same pathogens and/or diseases suggests that we have identified some of the significant infectious challenges encountered by ancient Bronze Age populations (Extended Data Figure 9.1).

Within the MS and RA analyses, we often identified protective SNPs in different genes selected for by the same pathogen(s)/diseases within the same or different ancestry paths. We also observed some shared SNPs between ancestral paths that were selected by the same pathogen(s)/disease. However, the number of MS- and RA-associated SNPs linked to protection against a given pathogen or disease could vary significantly.

In the MS analysis, one EBV-protective SNP was observed (~3% of all SNPs), in the ANA path (rs2735076; HLA-A). In the RA analysis, one SNP (~2% of all SNPs) was linked to a lower risk of EBV infection (rs3130062, ATP6V1G2) and positive selection was identified in the WHG and pan-ancestry paths. This result suggests that adaptation to decrease the risk of infection by a given pathogen - or the risk associated with a given disease once infected - can occur through modifications of genes affecting multiple different immune response pathways which might haphazardly increase or decrease the risk associated with a given autoimmune disease. In some populations, this adaptation increases or decreases the risk of MS or RA through one or several immune pathways; in other populations, the risk is increased or decreased through changes in different immune pathways or combinations of pathways. The population structure of EBV correlates with the genetic ancestry of infected populations and EBV diversification has been found to be shaped by host immune responses (Wegner et al., 2019).

It is perhaps surprising that MS-associated risk SNPs can be associated with protection against EBV, a significant risk factor for MS, especially if people are infected after puberty (Bjornevik et al., 2022). It is tempting to speculate that the protective effects of some of these allele variants might be age-dependent, but no data exists to prove or disprove the suggestion.

Distinct protective mumps alleles were found in the MS and RA analyses in the pan-ancestry path and in the CHG or ANA paths, respectively. Accurate estimates of the origin of mumps are not available at present, but Hippocrates described an outbreak of mumps on the Greek island of Thasos at approximately 2432 YBP, suggesting that the selective pressure from mumps could have been significant even earlier, and that mumps was not endemic in the Bronze Age but occurred as local and severe outbreaks.

The many unique SNPs within the different populations associated with protection against the same pathogen or disease(s) suggest that population-specific adaptations were common. In the MS selection analysis, we found one SNP linked to protection against HSV in the ANA path (rs2735076, HLA-A). In contrast, in the RA analysis, two SNPs were linked to protection against HSV and were selected in the ANA, CHG, WHG, EHG and pan-ancestry paths (rs9927316, LINC02132-LINC01082), and in the WHG (rs72887765, RGL2) path. The HSV-protective SNPs in the RA analysis were different from the one observed in the MS analysis. These results suggest that HSV circulated in all the analysed Bronze Age populations. Phylogenetic analyses have estimated that the European lineage of HSV originated approximately 5000 years before the present (YBP) (Guellil et al., 2022). This result suggests that the Bronze age lifestyles, increased population densities and population dispersals resulted in an increased incidence of HSV facilitating the generation of a European HSV lineage.

In the MS selection analysis, four of the 32 SNPs (~13%) were associated with protection against chickenpox or shingles/zoster (caused by VZV) and each were found in unique genes. One SNP was selected exclusively in the CHG path (rs7747873, HLA-E), one in the WHG and CHG paths (rs2735046, HLA-F), one in the EHG, WHG, and pan-ancestral paths (rs17576984, PGBD1), and one in the CHG and ANA path (rs9266773, MICA). Similarly, in the RA analysis, five SNPs were associated with protection against VZV-associated disease (9%). Of these SNPs, one was only associated with protection against VZV and was observed in the CHG (rs2523668, MICB) path, while the rest also have been associated with protection against other pathogens/diseases. The percentage of protective SNPs against VZV was higher than against any other specific pathogen in both the MS (4/32, 13%) and RA (5/55, 9%) analyses, making it tempting to speculate that the selective pressure on the Bronze Age populations from this virus might have been especially strong. VZV appears to be a uniquely human pathogen as no animal reservoir has been identified. Although VZV might have migrated with modern humans out of Africa, the greatest diversity is found in Europe and the VZV strains currently in existence seems to have originated in Europe about ~5000 years ago (Weinert et al., 2015; Pontremoli et al., 2019), i.e. during the Bronze age.

Our results highlight that interrogating SNPs associated with only one autoimmune disease risk can thwart the understanding of virus selection pressures in the analysed populations. We found no overlap between the many EBV- and VZV-associated protective SNPs in the MS and RA selection analysis.

Among the MS-risk SNPs, no protective alleles were found against measles (measles morbillivirus (MeV)). In contrast, among the RA-risk SNPs we identified one MeV protective SNPs in the EHG, ANA, CHG, and the pan-ancestry paths (rs707939, MSH5).

MeV is an exclusively human pathogen, and it is assumed, though not formally established, that it results from a spill-over from cattle infected with Rinderpest morbillivirus (RPV). MeV and RPV are closely related to Peste des petite ruminants virus (PPRV), which primarily infects wild and domestic goats and sheep, bharals, ibexes, and gazelles, but can cause infections in other hosts like camels and dogs (Dou et al., 2020). Phylogenetic analysis has estimated that the divergence between PPRV and the Mev/RPV ancestor happened at approximately 5221 YBP (95% HPD: 6655 - 3923 YBP), coinciding with an increase in pastoralism (Dux et al., 2020). The divergence between MeV and RPV is estimated to have occurred at about 2530 YBP (95% HPD: 3196 - 2187 YBP) (Dux et al., 2020). These estimates are calculated based on very few old MeV sequences and are uncertain. Although RPV primarily infects cattle, it also can infect humans. As the host range of the common ancestor to MeV/RPV is unknown, selective pressure from morbillivirus on ancient populations might predate the MeV/PRV split. Nevertheless, these results suggest that MeV, and possible other morbilliviruses, might have exerted a selective pressure on human populations since ~5000 YBP, and for sure since 2500 YBP, which is in line with the MeV-associated protective alleles identified in our RA-risk SNP analysis.

In the MS analysis, we observed a SNP linked to protection against spirochetes. It is unlikely to have been positively selected because it protects against syphilis; this disease is caused by treponema pallidum and the prevailing, but unproven, theory is that syphilis was brought into Europe (Naples, Italy) in 1493 AD by the ship from the New World (America). However, new studies have demonstrated a high diversity in the treponema pallidum family in Europe during the last ~1500 years and have estimated the origin of the European lineages at approximately 5000 YBP (Majander et al., 2020). Moreover, the spirochete protective SNP could also, or only, have been selected to combat Borrelia burgdorferi, a bacteria transmitted to humans through the bite of infected ticks causing Lyme disease.

We observed one SNP that exclusively protected against infection with mycobacteria including tuberculosis in WHG in the MS selection analyses (rs4902647, ZFP36F1) and one weakly protective allele against respiratory tuberculosis in the RA selection analysis which was selected in the EHG path (rs2596543, MICA). MICA has been demonstrated to be involved in the immune response to several bacterial infections apart from tuberculosis; these include brucellosis and listeriosis (Das et al., 2001).

HLA DRB1*15:01 is associated with protection against TB in European populations but not in other ethnic groups (Tervi et al., 2022). Studies have suggested a Neolithic emergence of the Mycobacterium tuberculosis complex (Bos et al., 2014; Sabin et al., 2020), with a particularly strong disease burden in the last 2,000 years (Kerner et al., 2021). TB can be transmitted through respiratory aerosols and humans can contract zoonotic TB from cattle (Mycobacterium bovis) through consumption of unpasteurised milk and handling sick animals (Gompo et al., 2020). Human TB can also, but less frequently, infect cattle (Lombard et al., 2021). Cold climate, overcrowding, and consumption of unpasteurized milk is known to facilitate transmission and lack of UV light and Vitamin D can increase the risk of TB reactivation (Fares 2011).

HLA DRB1*04:01 is associated with increased risk of RA and with better outcomes in, e.g., hepatitis C virus (HCV) (Hydes et al., 2015), Hepatitis B virus (HBV) (Yan et al., 2012), and SARS-CoV-2 infections (Langton et al., 2021), and with strong immune responses to influenza peptides (Danke & Kwok, 2003). We have found that RA risk was greatest in WHG and EHG and that the overall risk of RA has been under negative selection since the Bronze Age. It might be important to underscore that we estimate genetic risk in WHG and EHG, not disease incidence. While the genetic risk in WHG and EHG was high, their exposure to several respiratory or gastrointestinal pathogens linked to triggering RA was probably low. Consequently, their high genetic risk likely did not necessarily translate into high disease frequencies because of a lack of exposure to pathogen triggers. However, when these genetic risk factors were then present during the Bronze Age, the increased population sizes and the associated increased risk of transmission of 'crowd diseases', e.g., influenza, coronavirus infections, measles and morbilli, suddenly increased the risk of triggering RA pathogenesis. The inflammatory immune responses linked to RA pathogenesis have been estimated to approximately double the risk of a serious outcome of subsequent infections (Listing et al., 2013). Consequently, the effect of novel lifestyles during the Bronze Age might have resulted in a negative selection of alleles conferring a genetic risk of RA.

We found that most of the RA-risk SNPs linked to protection against pathogens and/or infectious diseases were protective against pathogens and/or diseases infecting mucosal surfaces. This result is somewhat surprising as multiple RA studies suggest that mucosal surfaces might be sites of inflammation and potential generation of RA-related autoimmunity in the preclinical period of the disease (Deane at al., 2017). However, this adaptation pattern to pathogens that might be disease triggers is in line with our observation that two MS-associated risk SNPs are also associated with protection against EBV, a major environmental MS risk factor. It is tempting to speculate that our RA results might reflect host adaptation in genes related to RA-risk that aim to reduce the risk of infection or serious illness by pathogens that might trigger RA. If this speculation is correct, then some of the SNPs linked to protection against a given pathogen might reveal which pathogens are most likely to trigger autoimmune disease. Determining whether these speculations are correct will require further studies.

Several SNPs were associated with protection against bacterial and viral pneumonia. One SNP was associated with protection against influenza in the MS analysis in the EHG and pan-ancestry paths, and one was identified only in WHG in the RA analysis. None of the SNPs selected in the MS and RA analysis overlapped.

The MS and the RA selection analyses revealed several protective alleles against parasites within many population paths suggesting that parasites exerted strong selective pressure on each population. None of these SNPs were overlapping between the MS and RA analyses. Likewise, several SNPs were associated with cellulitis and abscesses within the MS-risk SNPs (n=4), and within the RA-risk SNPs (n=6), but none overlapped. Cellulitis and abscesses are usually caused by bacteria, typically staphylococcus or streptococcus, that enter the skin and subcutaneous tissue through cuts and abrasions. Such lesions are likely to occur frequently, and our results suggest that strong selective pressures acted on reducing the harmful effects of these bacterial infections.

We found protective alleles against unique pathogens or diseases among the MS-risk and the RA-risk SNPs. The unique specific pathogens or diseases found among the MS-risk SNPs included streptococcus pneumonia (rs9296015, NOTCH4, WHG; rs3129934, TSBP1, WHG and pan-ancestry paths) and spirochetes (rs2248372, HCP5, MICB, WHG). Among the RA-risk SNPS, three SNPs reduced the risk of unique diseases or pathogens: pneumococcal pneumonia (rs2856822, HLA-DPA1, ANA, WHG), Entamoeba histolytica (rs3128947, HLA-DOA, WHG), and meningococcal infection (rs8075737, NEUROD2, WHG). Entamoeba histolytica causes amebiasis or amebic dysentery; today about 1/500 recorded infections are fatal.

Our analysis of MS- and RA-risk SNPs suggests that the growing population sizes and extensive population dispersals of the Bronze Age likely facilitated zoonosis (e.g., possibly MeV) and disease transmission of e.g., HSV and VZV. The infectious challenges from new pathogens and the more intense spread and exposure to old pathogens, e.g. through overcrowded living conditions, the use of manure and human waste as fertilisers and consumption of unpasteurised dairy products, likely resulted in the selection of strong anti-parasite Th2 responses and strong antiviral and antibacterial Th1 responses. Because Th2 responses hamper the efficiency of Th1 responses, an increase in the strength of the former will necessitate an increase in the latter if survival is to be ensured. While the immune response adaptations during the Bronze Age facilitated survival, today, in places with, e.g., better hygiene and less exposure to parasites, these strong immune responses can misfire and result in autoimmune diseases.
